# Disruption of cAMP Signalling and RNA Regulation Confers Cefiderocol Resistance in *Acinetobacter baumannii*

**DOI:** 10.64898/2026.10.08.755845

**Authors:** David M. P. De Oliveira, Ruby Heaton, Hayden Whyte, Larissa Buedenbender, Anuk D. Indraratna, Nguyen Thi Khanh Nhu, Minh-Duy Phan, Saleh F. Alquethamy, Yujiro Hirose, Budi Permana, Amy Wright, Anton Y. Peleg, Micaela Nochomovitz, Patrick N.A. Harris, Kay A. Ramsay, Philadelphia Holmes, Mark A. T. Blaskovich, David L. Paterson, John D. Boyce, Arun Everest-Dass, Mark A. Schembri, Jai J. Tree, Mark J. Walker, Brian M. Forde

**Author notes:** These authors contributed equally to this work (David M. P. De Oliveira, Brian M. Forde).

## Abstract

Cefiderocol is a siderophore-conjugated cephalosporin used to treat severe multidrug-resistant Gram-negative infections. Its recent prioritisation for US national preparedness underscores its clinical importance, yet resistance can emerge despite apparent in vitro susceptibility. Here, we identify two regulatory routes to cefiderocol resistance in carbapenem-resistant *Acinetobacter baumannii* (CRAB): mutation of the cAMP-producing adenylate cyclase CavA and truncation of the glycine-rich RNA-binding domain of the post-transcriptional regulator Hfq. These variants occurred in contemporary clinical isolates, conferred resistance in a murine CRAB lung-infection model and imposed minimal fitness costs. In *Escherichia coli*, deletion of the CavA orthologue *cyaA* increased competitive fitness during sub-inhibitory cefiderocol exposure. Hfq-CLASH and integrated omics revealed Hfq-dependent regulation of Fur-associated iron homeostasis in *A. baumannii.* Under iron limitation, truncated-Hfq induced a defence-primed state that attenuated siderophore uptake and reduced cefiderocol activity. These findings identify bacterial regulatory state and iron availability as determinants of cefiderocol susceptibility, with implications for resistance surveillance and susceptibility testing.

## Introduction

*Acinetobacter baumannii* is an adaptable, opportunistic pathogen responsible for an increasing burden of healthcare-associated infections worldwide^1–3^. Carbapenem-resistant *A. baumannii* (CRAB) is independently associated with elevated mortality, with invasive infections incurring death rates exceeding 40%, driven in part by the paucity of effective treatment options^4,5^. Compared to carbapenem-susceptible *A. baumannii*, CRAB imposes substantial healthcare costs through prolonged hospital stays, increased intensive care admissions, and limited therapeutic alternatives. Reflecting this threat, the World Health Organization has classified CRAB as a “critical priority” pathogen, underscoring the urgent need for novel therapeutics and improved surveillance strategies^6^.

Against the backdrop of a stagnating antibiotic development pipeline, Cefiderocol (Fetroja) remains the only stand-alone antibiotic (non β-lactam, β-lactamase combination) approved since 2019 for the treatment of complicated Gram-negative infections, including complicated urinary tract infections, bloodstream infections and severe CRAB infection^7^. Cefiderocol is a cephalosporin-siderophore conjugate antibiotic that displays high affinity for penicillin-binding proteins (PBPs), notably for PBP3, leading to inhibition of cell wall synthesis^8,9^. The chlorocatechol group on the cefiderocol C3-side-chain enables chelation of ferric (Fe³⁺) iron, allowing the drug to mimic bacterial siderophores and hijack iron uptake systems^10^. By exploiting host-induced iron limitation during infection, cefiderocol-Fe³⁺ complexes are actively transported across the outer membrane into the periplasm^11,12^, providing potent antibacterial activity against carbapenem-resistant Gram-negative pathogens.

While cefiderocol resistance has traditionally been attributed to mutations in TonB-dependent siderophore receptors (e.g., *pirA*), production of PER-1 β-lactamase (*bla_PER-1_*), alterations in PBP3 (*ftsI*), reduced outer membrane permeability or increased efflux, these changes do not fully account for clinical resistance and treatment failure^13–16^. Notably, mutations conferring cefiderocol resistance in *A. baumannii* are highly heterogeneous and non-conserved across isolates, suggesting that resistance may arise from pre-existing antimicrobial resistance determinants rather than *de novo* selection under antibiotic pressure^17^. Thus, our incomplete understanding of cefiderocol resistance mechanisms may complicate future efforts to predict and mitigate the emergence of resistance selected through cefiderocol exposure.

In this study, we have identified and experimentally confirmed novel mutations in the global regulators *cavA* (adenylate cyclase) and *hfq* (sRNA-mRNA chaperone) that confer cefiderocol resistance in CRAB upon exposure. Such mutations are also found in contemporary clinical CRAB isolates, indicating their epidemiological relevance beyond laboratory selection. Thus, our findings reveal previously unrecognised resistance mechanisms driven by global regulatory circuits, expanding the current resistance paradigm beyond traditional β-lactamase expression and porin loss. By establishing *cavA* and *hfq* genetic variants as clinically relevant determinants of cefiderocol resistance, we provide a rational foundation for developing targeted molecular markers to enable rapid, reliable detection of cefiderocol-resistant CRAB, addressing a critical knowledge gap in surveillance and guiding the design of improved diagnostic strategies.

## Results

### AB5075 rapidly develops resistance to cefiderocol upon exposure

To investigate genetic pathways contributing to cefiderocol resistance, we conducted resistance development studies using the CRAB clinical isolate AB5075, a strain characterised by multi-drug resistance, high virulence and genetic tractability^18^. Cefiderocol susceptibility was assessed using Clinical and Laboratory Standards Institute (CLSI) guidelines as the European Committee on Antimicrobial Susceptibility Testing (EUCAST) does not yet define a breakpoint for *Acinetobacter*^19,20^. AB5075 was susceptible (minimum inhibitory concentration [MIC] 0.5 µg/mL, CLSI susceptible breakpoint ≤ 4 µg/mL), but serial passage in sub-inhibitory cefiderocol produced a progressive, stepwise MIC increase over 13 days, culminating in a 512-fold rise to 128 µg/mL (Fig. 1A).Comparative whole-genome sequencing (WGS) analysis of the resistant mutants identified two sequentially acquired chromosomal mutations (Table S1). The first was a single base change in *cavA* (ABUW_2208), encoding the sole functional adenylate cyclase in AB5075^21^, giving rise to an F237V missense mutation. Secondary structure modelling localised this surface-exposed substitution within the C-terminal catalytic domain that converts ATP to cAMP (Fig. S1)^21^. The AB5075*^cavA^* mutant showed significantly reduced cAMP production relative to AB5075 wild-type (WT), indicating a critical role for residue F237 in CavA-mediated cAMP generation (Fig. S1D).

**Figure 1.**
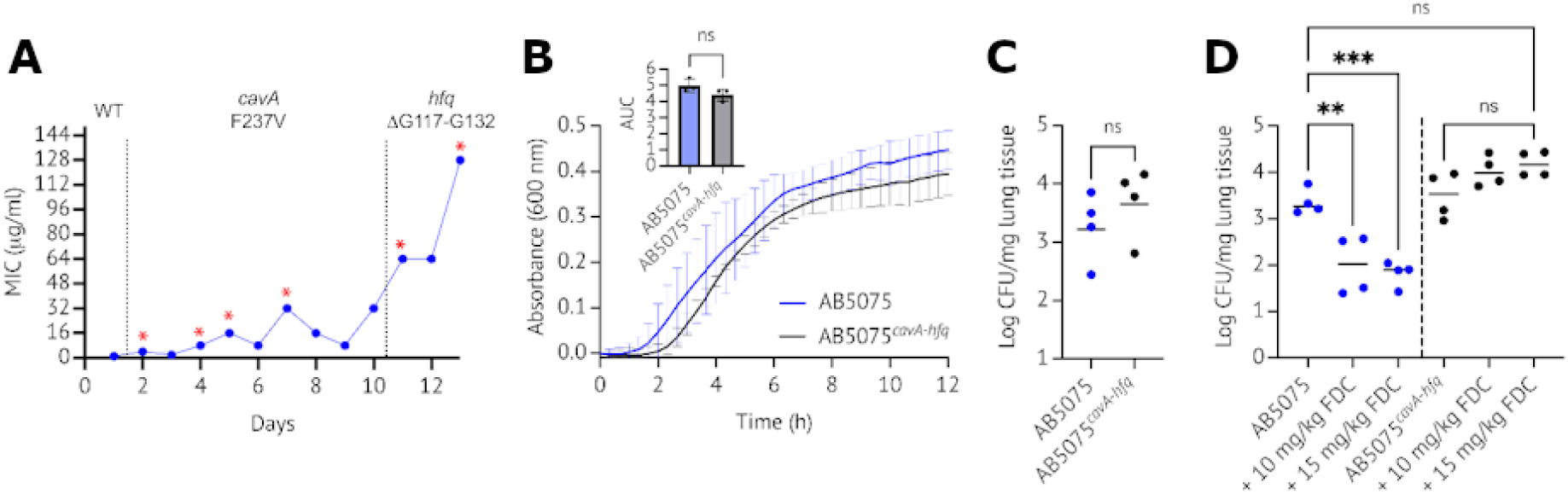
Mutations in adenylate cyclase (cavA) and RNA chaperone (hfq) genes confer high-level cefiderocol resistance. (A) Serial passage of susceptible AB5075 with cefiderocol in ID-CAMHB. Asterisks denote time points where single colonies were isolated from resistant cultures for whole-genome sequencing. Dotted lines indicate the period in time where each mutation was identified. (B) In vitro growth comparison of AB5075 and AB5075^cavA-hfq^ mutants in ID-CAMHB. The inset shows area under the curve (AUC) analysis for statistical significance. (C) In vivo fitness of AB5075^cavA-hfq^ in a murine model of A. baumannii lung infection. Cohorts of 12-week-old, sex-matched mice were intranasally challenged with 5 × 10⁷ CFU AB5075 WT or AB5075^cavA-hfq^; CFUs recovered from lungs after 24 h. (D) In vivo cefiderocol resistance assessment. Sex-matched BALB/c mice were intranasally challenged with 5 × 10⁷ CFU of either AB5075 WT or AB5075^cavA-hfq^, followed by intraperitoneal treatment with vehicle or cefiderocol (FDC) at either 10 mg/kg or 15 mg/kg. Treatments were administered at 0 and 6 h post-infection. CFUs recovered from lungs 24 hr post-challenge. Values for individual mice are shown (**P ≤ 0.01; ****P ≤ 0.0001; one-way ANOVA with Tukey’s multiple comparisons)

The second mutation was a deletion in *hfq* (ABUW_1609; ΔG117→G132), which encodes a major RNA chaperone involved in post-transcriptional gene regulation via sRNA–mRNA interactions^22^. *A. baumannii* Hfq has a long, disordered C-terminal tail of 7 glycine-rich repeats (GGFGGQ) required for high-affinity sRNA binding^22^; the deletion removed five of the seven repeats. Hfq truncation increased the cefiderocol MIC from 8 µg/mL in the *cavA* F237V first-step mutant to 128 µg/mL in the *hfq* deletion second-step mutant. The intrinsic disorder and flexibility of these motifs precluded resolution of structural differences using secondary structure modelling^22,23^. As with AB5075*^cavA^*, AB5075*^cavA-hfq^* demonstrated significantly reduced intracellular cAMP production relative to AB5075 WT (Fig. S1D).

### Mutations in CavA and Hfq confer cefiderocol resistance in vivo *without imposing a fitness* cost

Fitness assessment of AB5075*^cavA-hfq^* demonstrated equivalent growth rate compared to AB5075 WT when cultured in iron-depleted cation-adjusted Mueller Hinton broth (ID-CAMHB) (Fig. 1B). This absence of a fitness defect was further supported *in vivo*, where AB5075*^cavA-hfq^* showed no significant difference in bacterial clearance relative to AB5075 WT in a 24 h immunocompetent mouse model of CRAB lung infection (Fig. 1C). The cefiderocol resistance phenotype of AB5075*^cavA-hfq^* was also confirmed *in vivo*, whereby intraperitoneal administration of cefiderocol failed to provide therapeutic benefit in mice intranasally challenged with AB5075*^cavA-hfq^*. In contrast, cefiderocol demonstrated a significant therapeutic effect against AB5075 WT in this same lung infection model (Fig. 1D).

### Mutations in cavA and hfq drive convergent regulatory, proteomic and phenotypic remodelling

To define the effects of each mutation on global gene regulation and subsequent cefiderocol resistance, we performed comparative transcriptomic and proteomic analyses across AB5075 WT, AB5075*^cavA^* and AB5075*^cavA-hfq^*. In *A. baumannii*, CavA catalyses cAMP synthesis, which acts as a signalling molecule through binding to the cAMP receptor protein Vfr to regulate downstream gene expression^21^. Relative to AB5075 WT, AB5075*^cavA^* demonstrated increased expression of genes involved in exopolysaccharide production (*pgaABCD*), antibiotic tolerance (*ecnB*), and cell turgor and osmotic stress response (*aqpZ*)^24–26^, together with reduced expression of sulfur transport genes (*cysPTW* and *tauABCD*), suggesting altered sulfur metabolism (Fig. 2A; Table S2)^27^. The proteome mirrored these changes, with increased abundance of PgaAB, required for synthesis of the biofilm matrix polysaccharide poly-N-acetylglucosamine, and the CsuABCDE chaperone–usher pilus proteins, alongside reduced abundance of the biofilm negative regulator BigR and Vfr itself, consistent with reduced cAMP-dependent Vrf activity (Fig. 2C; Table S4). Notably, twitching motility proteins (PilBDGHIKOPQRTUWXYZ) were significantly depleted despite unchanged *pil* transcription, underscoring multi-layered post-transcriptional regulation. GO enrichment of low-abundance proteins in AB5075*^cavA^* identified changes in cell envelope remodelling and stress responses consistent with a biofilm-associated tolerance state (Fig. 2D). These molecular changes were reflected phenotypically, with both AB5075*^cavA^* and AB5075*^cavA-hfq^* producing significantly more biofilm than AB5075 WT at 24 h and showing significantly reduced twitching motility (Fig. S2A,B).

**Figure 2.**
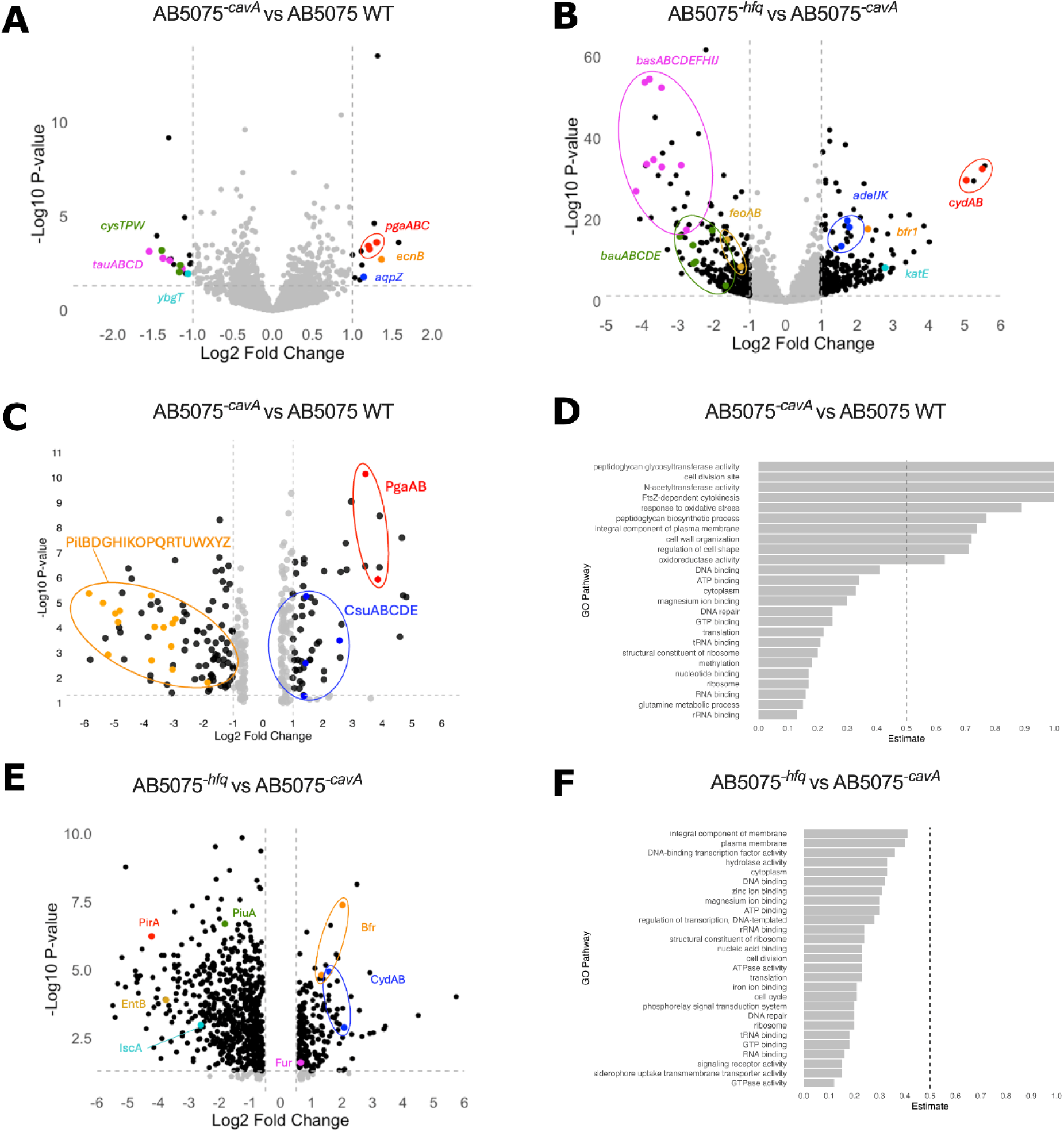
CavA F237V and Hfq truncation drive coordinated transcriptomic and proteomic remodelling in Acinetobacter baumannii AB5075. RNA-seq volcano plots compare (A) AB5075cavA with wild-type (WT) AB5075 and (B) AB5075cavA-hfq with AB5075cavA. Proteomic volcano plots and corresponding Gene Ontology (GO) enrichment analyses show the same comparisons in (C, D) and (E, F), respectively. All strains were grown in iron-depleted cation-adjusted Mueller–Hinton broth (ID-CAMHB). For RNA-seq (A, B), black points indicate genes with an absolute log2 fold change >1 and P <0.05. Highlighted genes are associated with biofilm formation (pgaABCD), stress responses (ecnB, aqpZ, katE), sulfur metabolism (cysPTW, tauABCD), respiration (ybgT, cydAB), iron acquisition and storage (bas, bau, feoAB, bfr), and antimicrobial resistance (adeIJK, ampC). RNA-seq data represent three independent biological replicates. For proteomics (C, E), black points indicate proteins with an absolute log2 fold change >0.5 and P <0.05. Highlighted proteins are associated with biofilm formation, adherence and motility (PgaAB, CsuABCDE, PilBDGHIKOPQRTUWXYZ), regulation (Vfr, BigR), iron homeostasis (PfeA, PiuC, BasGH, FeoAB, IscX and ErpA), oxidative stress (KatE), and drug efflux (AdeN and AdeIJK). In (D, F), grey bars show estimated GO-term activity; estimates <0.5 indicate repression and >0.5 indicate activation, with the dashed line marking 0.5.

Hfq is a critical RNA chaperone in *A. baumannii*, modulating growth, stress tolerance, virulence, biofilm formation and antibiotic resistance^22,28^. Comparative transcriptomic analysis of AB5075*^cavA^* and AB5075*^cavA-hfq^*, in which five of seven Hfq C-terminal repeats were deleted, revealed extensive transcriptional changes associated with Hfq truncation (Fig. 2B; Table S3). Consistent with these findings, proteomic analysis of AB5075*^cavA-hfq^* identified parallel changes alongside high-level cefiderocol resistance (Fig. 2E; Table S4). Two major mechanisms emerged. First, Hfq truncation enhanced efflux and β-lactamase activity. The resistance-nodulation-division efflux pump system *adeIJK* and the chromosomal β-lactamase *ampC* were significantly upregulated, with *ampC* upregulation previously associated with cefiderocol resistance and clinical treatment failure in *Enterobacter cloacae*^29^ (Table S3). Consistent with the transcriptomic data, AdeIJK proteins were elevated, while the AdeIJK negative regulator AdeN was reduced, indicating derepression of the AdeIJK efflux system ^30^.

Second, Hfq truncation impaired iron acquisition and siderophore-mediated uptake. Under iron-depleted conditions (ID-CAMHB), genes involved in acinetobactin biosynthesis (*bas* operon), siderophore receptor (*bau* operon) and TonB-dependent siderophore transport, including *pfeA*, which collectively mediate Fe³⁺ uptake and cefiderocol entry, were significantly downregulated^31,32^. These changes were mirrored at the protein level, with reduced abundance of the putative TonB energy transducer (UniProt A0A0D5YE91), the outer membrane siderophore receptors PfeA and PiuC, the acinetobactin permease BauC, and the Fe–S cluster assembly proteins ErpA and IscX (Fig. 2E; Table S4)^33^. GO enrichment further supported disruption of siderophore uptake transmembrane transport, TonB-dependent receptor activity, together with broader changes in DNA/RNA regulatory networks (Fig. 2F).

Additional changes were consistent with a broader envelope stress and metabolic adaptation, including increased *cydAB* expression, indicative of increased reliance on the cytochrome bd terminal oxidase^26,34^. Increased upregulation of the bacterioferritin gene (*bfr2*) and catalase gene (*katE*) expression were also observed, accompanied by elevated KatE peroxidase protein, consistent with enhanced iron storage and oxidative stress defence^35^. CsuA/B fimbrial and OmpA-family proteins were also increased, whereas genes associated with twitching motility, pilus assembly (*pil*) and competence (*com*) were reduced^36^.

Together, these findings support a regulatory role for Hfq in iron homeostasis under host-imposed nutritional immunity, consistent with Hfq-dependent post-transcriptional control of Fur, the master regulator of iron homeostasis, in *E. coli*^37,38^.

### Hfq interacts with mRNAs associated with iron homeostasis and drug efflux

To elucidate Hfq-dependent post-transcriptional networks underlying cefiderocol resistance, UV cross-linking, ligation and sequencing of hybrids (CLASH) analysis was performed using a closely related *A. baumannii* strain (AB307) with wild-type Hfq encoding seven glycine-rich repeats, and that contained a chromosomally tagged Hfq used as bait (Fig. 3A; Table S5)^39^. Hfq-CLASH sequencing of RNAs interacting with Hfq in iron-replete and iron-depleted media identified a Hfq binding site within the *fur* mRNA specifically under iron-replete conditions (Fig. 3A). Analysis of RNA-RNA interactions at this site identified an interaction with sRNA97 (ABBFA_004058_sRNA; also known as CsrB; 88 hybrid reads, FDR 4.03×10^−208^) consistent with post-transcriptional repression of *fur* by Hfq-sRNA97 (Table S5). Integration of the Hfq-CLASH data with RNA-seq from the AB5075*^cavA-hfq^* 2-repeat Hfq deletion mutant revealed that 47.3% of upregulated genes (>2 log_2_FC and *p*_adj_ <0.05) were detected in at least one sRNA-mRNA interaction, with the majority mediated by the abundant iron-limitation-induced sRNA97. These findings suggest that a substantial fraction of genes upregulated in the Hfq deletion mutant reflect loss of sRNA-mediated repression, predominantly via sRNA97. Conversely, many genes downregulated in the Hfq truncation mutant are known members of the Fur regulon. Loss of Hfq–sRNA97 repression of *fur* would be expected to increase Fur levels under iron-replete conditions, thereby enhancing repression of Fur-regulated genes.

**Figure 3.**
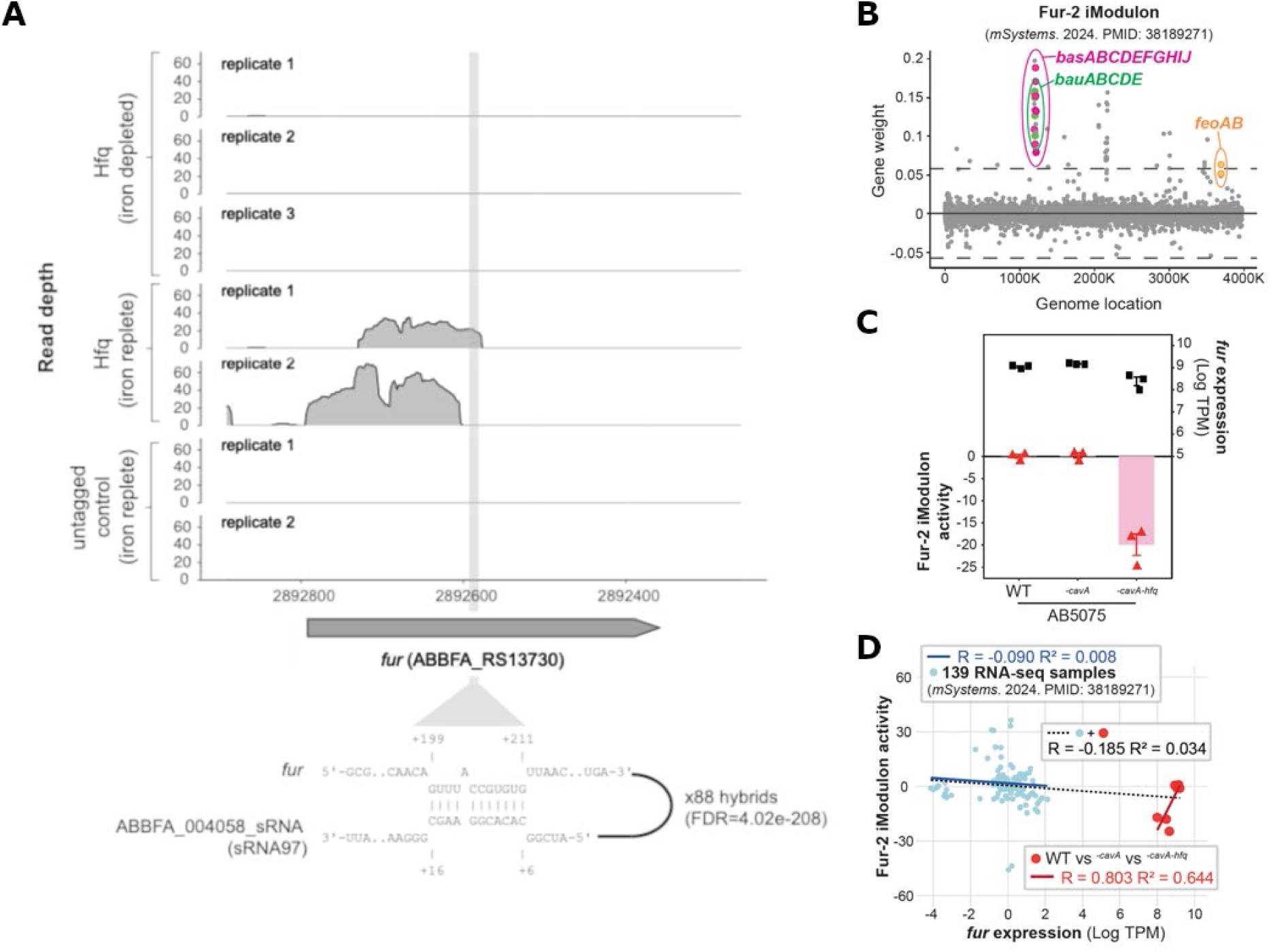
Hfq-CLASH reveals sRNA99-fur interaction and Fur-2 iModulon confirms post-transcriptional Fur dysregulation in AB5075^cavA-hfq^ truncation mutant. (A) Hfq-CLASH performed in chromosomally Hfq-tagged AB307 under iron-replete and iron-depleted conditions identified specific Hfq binding within the fur mRNA exclusively in iron-replete media. RNA–RNA hybrid analysis revealed a high-confidence interaction between fur and sRNA99 (ABBFA_004058_sRNA; also known as CsrB; 88 hybrid reads, FDR = 4.02 × 10⁻²⁰⁸), consistent with Hfq-dependent post-transcriptional regulation. Untagged controls showed no enrichment. (B) Fur-2 iModulon gene weights derived from the AB5075 AbaumPRECISE139 compendium (mSystems 2024; PMID: 38189271) highlight strong contributions from iron acquisition operons (basABCDEFJHI, bauABCDE, feoAB), whereas fur exhibits minimal weight. (C) Quantification of Fur-2 iModulon activity across RNA-seq datasets from AB5075 WT, AB5075^cavA^, and AB5075^cavA-hfq^. The hfq-truncation mutant (AB5075^cavA-hfq^) displays decreased Fur-2 activity relative to AB5075 WT and AB5075^cavA^ (Wilcoxon test, **P < 0.01) despite comparable fur transcript abundance (log TPM). Boxplots indicate median and interquartile range. (D) Correlation between fur expression (log TPM) and Fur-2 iModulon activity across 139 publicly available AB5075 RNA-seq datasets shows a weak negative relationship (R = −0.090). Inclusion of our datasets minimally alters this trend (R = −0.185). In contrast, within our dataset alone, Fur-2 iModulon activity positively correlates with fur expression (R = 0.803), indicating context-dependent decoupling of fur transcription and regulon activity.

Iron acquisition operons (*basABCDEFJHI*, *bauABCDE*, and *feoAB*) belong to the Fur regulon in *A. baumannii*. Transcriptome-wide independent component analysis (ICA), which delineates modules of co-expressed genes (iModulons), assigns these operons to the Fur-2 iModulon in AB5075 (Fig. 3B; Table S6)^40–42^. Because iron-acquisition operons dominate this iModulon, Fur-2 activity reports expression of Fur-repressed genes and is therefore expected to vary inversely with Fur activity. We thus quantified Fur-2 iModulon activity across our RNA-seq datasets and compared it with *fur* transcript abundance. Relative to AB5075 WT and AB5075*^cavA^*, the AB5075*^cavA-hfq^* mutant exhibited decreased Fur-2 iModulon activity despite comparable *fur* expression across strains (Fig. 3C; Table S7).

To determine whether Fur regulon activity correlates with *fur* transcript levels more broadly, we next analysed 139 publicly available AB5075 RNA-seq datasets. Across these datasets, Fur-2 activity showed only a weak negative correlation with *fur* expression overall (R = −0.090; Fig. 3D), which did not appreciably change with the inclusion of our datasets (R = −0.185). In contrast, within our dataset, Fur-2 activity positively correlated with *fur* expression (R = 0.803). Collectively, these findings indicate that Fur regulon activity is not dictated solely by *fur* transcription. Instead, post-transcriptional mechanisms, such as Hfq-sRNA97-mediated regulation of *fur*, likely contribute to iron homeostasis dysregulation in the AB5075 Hfq truncation mutant.

### CavA F237V mutations and Hfq polymorphisms in contemporary clinical isolates of A. baumannii

To determine the prevalence of CavA F237V point mutations and Hfq polymorphisms among clinical *A. baumannii* isolates, we performed a comparative alignment analysis using 43,061 publicly available genome sequences from NCBI (accessed: 25/07/2025). Comparative alignment revealed that the surface-exposed CavA F237V mutation was rare (13 of 43,601 genomes; 0.03%) and restricted to contemporary clinical isolates, with 12 of 13 occurrences found in isolates from the United States (U.S.) and a single isolate from China (Table S8). All *A. baumannii* isolates harbouring the CavA F237V mutation were collected between 2019 and 2024, consistent with recent selective pressures, such as antimicrobial exposure. The U.S.-derived strains were confined to Pasteur multilocus sequence types ST79, ST78, and ST499. These represent distinct epidemiological lineages known for their involvement in regional outbreaks and multidrug resistance but not classified among the classic high-risk international clones GC1 (ST1) or GC2 (ST2)^43,44^. Cefiderocol susceptibility data were not available for these isolates, and they were not accessible for phenotypic testing. As such, the contribution of CavA F237V to cefiderocol susceptibility in these backgrounds therefore remains to be established.

Analysis of Hfq across *A. baumannii* genomes revealed polymorphic diversity within a predominantly canonical seven-repeat glycine-rich domain within the C-terminal region. Consistent with the cefiderocol-resistant AB5075*^cavA^*^−*hfq*^ mutant identified in this study, approximately 5% of full-length Hfq sequences (n = 1,865 of 39,710) encode truncated variants, possessing only two of the typical seven glycine-rich RNA-binding repeats (Fig. S3).

### Hfq polymorphisms prime A. baumannii for increased fitness and adaptive antibiotic resistance under iron-depleted conditions

CLSI guidelines for cefiderocol susceptibility testing specify broth microdilution in ID-CAMHB prepared from standard CAMHB, to promote siderophore-mediated drug uptake^45^. To investigate how Hfq-dependent post-transcriptional regulation may modulate cefiderocol resistance in response to changing iron availability, clinical *A. baumannii* isolates encoding either truncated 2-repeat Hfq or canonical 7-repeat Hfq were pre-cultured to mid-log phase in iron-replete CAMHB or iron-depleted ID-CAMHB prior to MIC determination in ID-CAMHB. Consistent with the transcriptional and proteomic signature of dysregulated iron uptake described above (Fig. 2B, 2E), all clinical isolates carrying truncated 2-repeat Hfq remained cefiderocol-susceptible when primed in iron-replete medium (CAMHB). However, after growth in ID-CAMHB these isolates displayed increased MICs, with isolates M00376, M87341, M87136, and M88825 exceeding the CLSI resistance breakpoint of ≥4 µg/mL (Table 1). By contrast, clinical isolates expressing canonical 7-repeat Hfq showed no change in cefiderocol susceptibility regardless of pre-culture conditions, highlighting the combined impact of Hfq polymorphism and strain background on cefiderocol resistance outcomes. We term this condition a ‘defence-primed state’, in which iron restriction activates Hfq-dependent post-transcriptional remodelling of Fur-regulated iron acquisition before cefiderocol exposure, thereby reducing siderophore-mediated uptake and limiting cefiderocol entry to the bacterial cell.

**Table 1:**
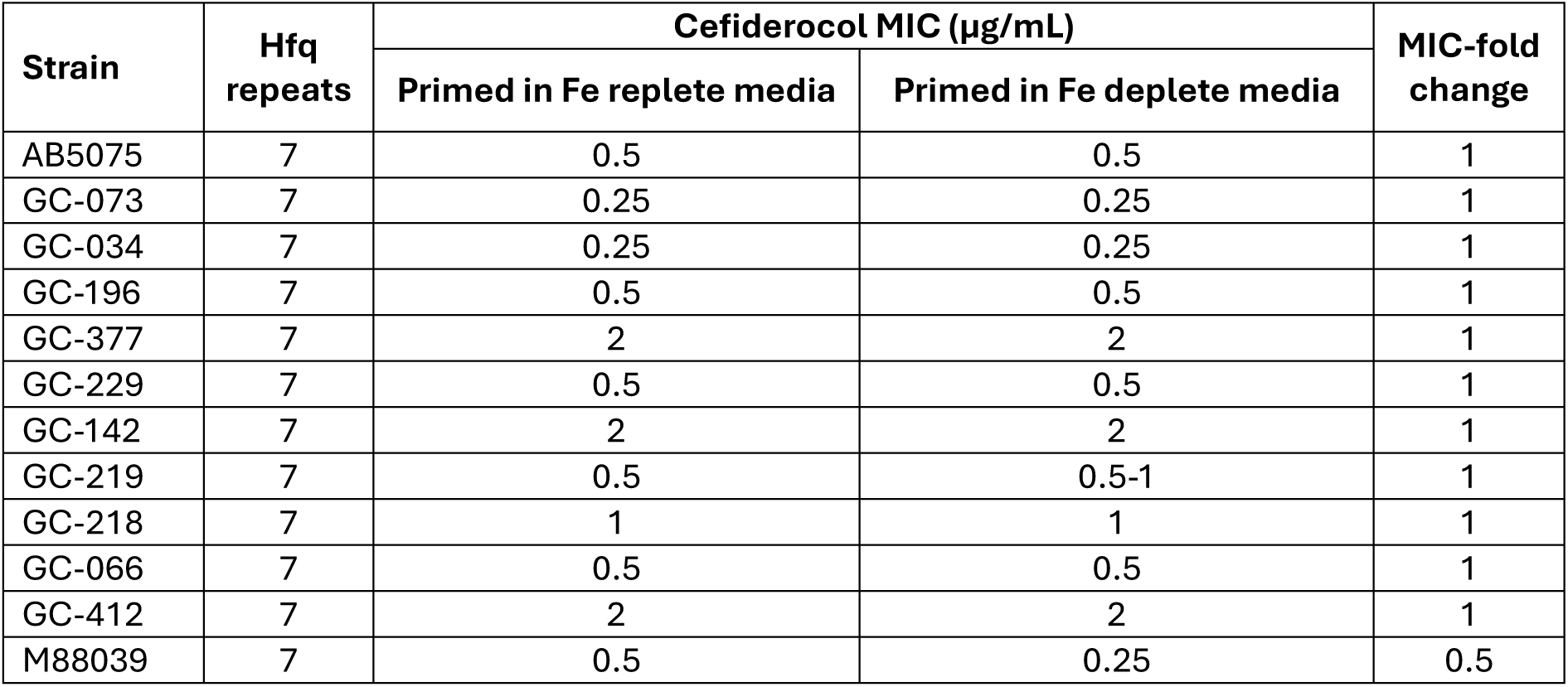
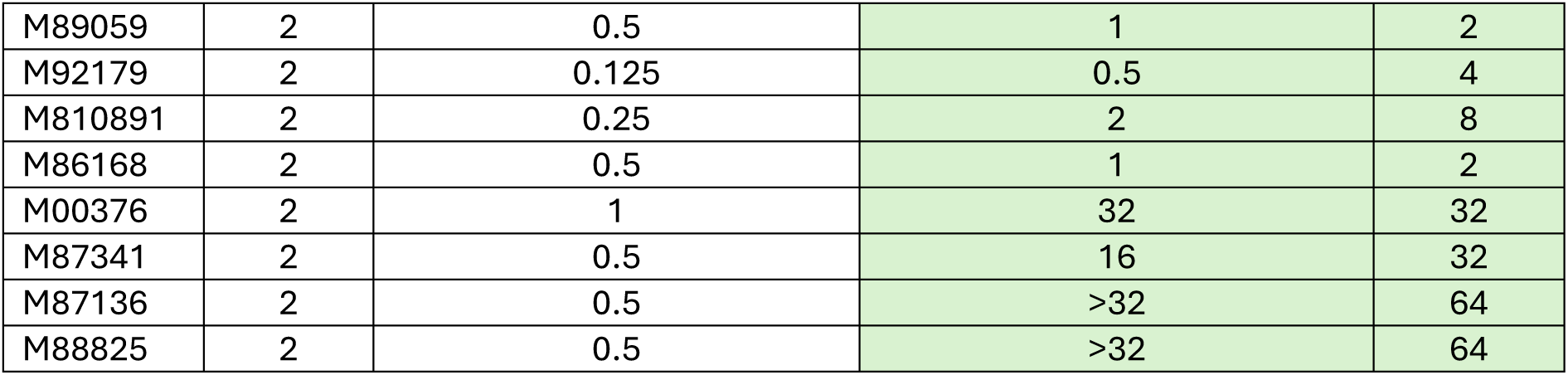
Iron-deplete conditions enhance cefiderocol resistance in CRAB isolates with truncated 2-repeat Hfq domains. A. baumannii isolates were sub-cultured and primed either in iron-replete media (CAMHB) or iron-deplete medium (ID-CAMHB) prior to MIC testing in ID-CAMHB in line with CLSI guidelines for cefiderocol susceptibility assessment. Increases in cefiderocol-resistance fold-change are highlighted in green.

To examine Hfq-dependent adaptation to iron limitation in more detail, quantitative real-time PCR targeting the acinetobactin transport genes *bauB* and *bauE* was performed after growth in iron-depleted conditions. Both *bauB* and *bauE* were selected as they are highly conserved across isolates AB5075, GC-073, GC-034, M00376, M87136 and M88825, and encode, respectively, the TonB-dependent outer-membrane receptor for ferric acinetobactin and the periplasmic siderophore-binding protein that delivers ferric acinetobactin to the inner-membrane transport complex^46,47^. Relative to iron-replete CAMHB, isolates harbouring canonical 7-repeat Hfq exhibited significantly greater induction of *bauB* and *bauE* in ID-CAMHB than isolates with truncated 2-repeat Hfq, suggesting that Hfq truncation attenuates activation of the acinetobactin uptake pathway under iron-limiting conditions (Fig. 4A). Collectively, these data support a model in which the 2-repeat Hfq variant disrupts normal regulation of iron uptake and storage, thereby impairing siderophore-dependent cefiderocol entry and favouring an adaptive, reversible resistance phenotype in low-iron environments. We further assessed this model by passaging 2-repeat Hfq isolates (M89059, M92179, M810891, M86168, M00376, M87341, M87136 and M88825) in ID-CAMHB with sub-inhibitory concentrations of cefiderocol for a period of 13 days. At 48h, post-cefiderocol exposure M00376, M87341, M87136 and M88825 exhibited increased MICs at or above the cefiderocol resistance breakpoint (MIC ≥4 µg/mL) (Fig. 4B). WGS of these derivatives revealed no chromosomal mutations relative to their parental strains (Table S9), consistent with a non-genetic adaptive response to iron-depleted cefiderocol-containing conditions rather than the acquisition of stable resistance mutations. These observations are compatible with cefiderocol heteroresistance. Subsequently, our findings have important implications for clinical cefiderocol susceptibility testing, indicating that Hfq polymorphisms and pre-test growth conditions can influence measured MICs and may unmask heteroresistance not captured by standard protocols.

**Figure 4.**
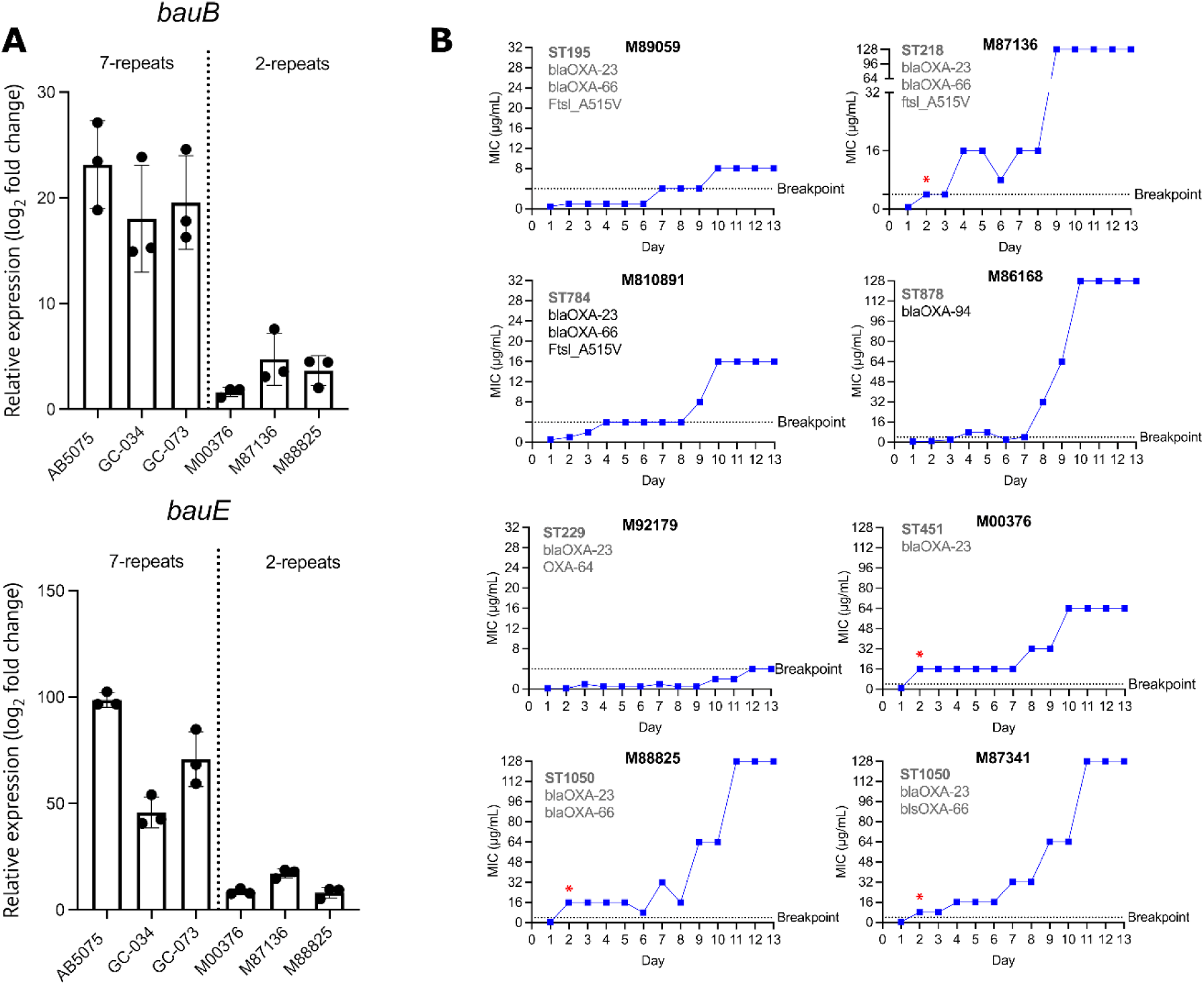
Truncated Hfq is associated with attenuated acinetobactin uptake gene induction and emergence of cefiderocol resistance. (A) Cefiderocol MIC trajectories during serial passage of eight initially susceptible, 2-repeat Hfq A. baumannii isolates in cefiderocol-containing ID-CAMHB. The dashed line indicates the cefiderocol resistance breakpoint, and red asterisks indicate time points at which colonies were selected for whole-genome sequencing. (B) Relative bauB and bauE expression in isolates encoding canonical 7-repeat or truncated 2-repeat Hfq following growth to mid-log phase in ID-CAMHB. Expression was calculated relative to growth in iron-replete CAMHB and normalized to gyrB using the ΔΔCT method. Bars show the mean ± SD of three independent biological replicates; assays were performed in technical triplicate.

### cavA mutations alter cefiderocol resistance in Escherichia coli

To investigate whether *cavA* or *hfq* mutations can confer cefiderocol resistance in other Gram-negative pathogens, transposon-directed insertion-site sequencing (TraDIS) was performed on *E. coli* strain MS7163 (ST95; serotype O45:K1:H7) under cefiderocol selection^48^. We plated approximately 2 × 10⁸ mutants on CAMHB agar supplemented with 0.125 µg/mL cefiderocol (2× MIC), yielding ∼4,736 resistant colonies, which we pooled for genomic DNA extraction and mini-Tn5 insertion-site sequencing.

TraDIS reads were significantly enriched in genes involved in iron uptake and cAMP signalling. Insertions were prominent in the *fecA* promoter (driving expression of this ferric ion importer)*, tonB* (encoding the TonB periplasm-spanning protein required for siderophore-iron transport), and *envZ* (encoding a sensor kinase that regulates outer membrane porin expression via OmpR)^49,50^. Notably, insertions were also detected in *cyaA* (the *A. baumannii cavA* orthologue, encoding adenylate cyclase) and *crp* (the *A. baumannii vfr* orthologue, encoding the cAMP-responsive transcriptional regulator), mirroring findings in cefiderocol-resistant *A. baumannii* AB5075 mutants and supporting a conserved role for cAMP signalling in cefiderocol resistance across Gram-negative species (Fig. 5A&5B; Fig. S4; Table S10). To validate the *cyaA* TraDIS hit, a targeted Δ*cyaA* mutant and a complemented strain were constructed in MS7163. Although the Δ*cyaA* mutant showed no measurable change in cefiderocol MIC compared to the parent strain in single-strain assays (Table S11), a more sensitive mixed 50:50 competition assay revealed a significant reduction in the mutant population in the absence of cefiderocol and complete competitive displacement by the Δ*cyaA* mutant in the presence of sub-MIC cefiderocol (Fig. 5C). These effects were abolished in the competition between the Δ*cyaA* complemented strain and MS7163, indicating that the phenotype was *cyaA*-specific (Fig. 5C). Consistent with this finding, a Δ*vfr* mutant generated in in the AB5075-related *A. baumannii* strain AB307-0294 showed a strong competitive advantage over the wild-type in the presence of sub-MIC cefiderocol concentrations, phenocopying the Δ*cyaA* mutant (Fig. 5D). Together, these finding support a model in which CavA influences cefiderocol tolerance through cAMP-dependent modulation of Vfr regulatory activity. No *hfq* insertions were detected, consistent with fundamental differences in Hfq structure and sRNA regulation between *E. coli* and *A. baumannii*, where *E. coli* Hfq lacks the glycine-rich C-terminal domain characteristic of *A. baumannii* Hfq.

**Figure 5.**
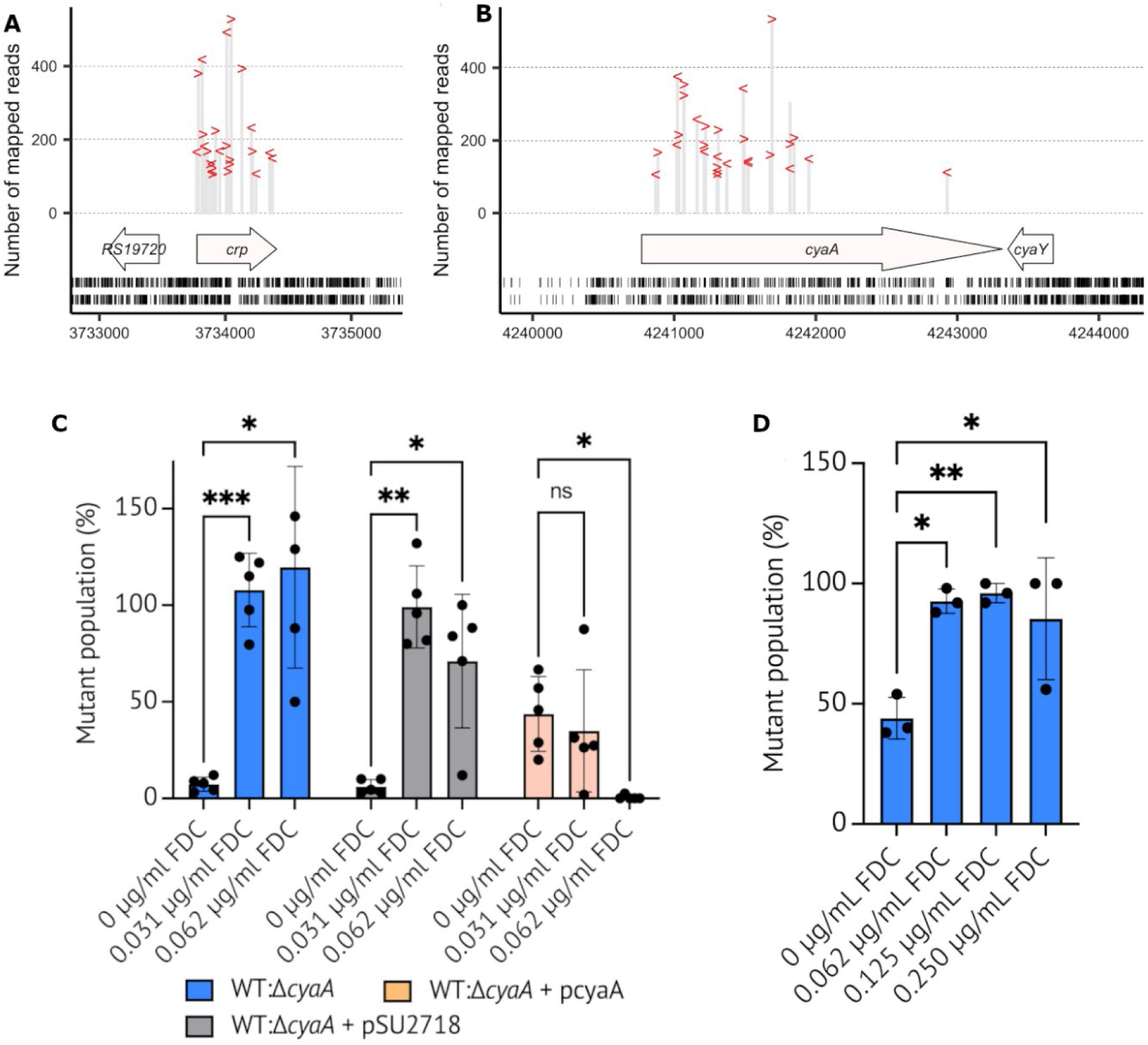
Disruption of cAMP signalling confers a selective advantage during cefiderocol exposure in Escherichia coli and Acinetobacter baumannii. Location and number of mapped reads per insertion site within the (A) crp and (B) cyaA genes of Escherichia coli strain MS7163. Each insertion site is represented by “>” (forward-strand insertion) or “<” (reverse-strand insertion). Horizontal arrows represent coding sequences. Insertion sites present in the input pool are indicated by short vertical lines below the coding sequences, with the upper and lower rows representing insertions on the forward and reverse strands, respectively. (C) Competition assays between the wild-type E. coli strain MS7163 and each of its derivatives, including the ΔcyaA mutant MS7163ΔcyaA (WT:ΔcyaA), the ΔcyaA mutant with an empty vector MS7163ΔcyaA + pSU2718 (WT:ΔcyaA), and the ΔcyaA mutant complemented with the cyaA gene on the pSU2718 vector MS7163ΔcyaA + pcyaA (WT:ΔcyaA + pcyaA), were conducted similarly to the broth microdilution method, where each competition mix was inoculated in ID-CAMHB in the presence of cefiderocol (0, 0.0313, 0.0625 µg/mL) and allowed to grow overnight. Data represent biological replicates. Statistical significance is indicated (****, P ≤ 0.0001; one-way ANOVA with Tukey’s multiple comparisons). (D) Competition assays between WT Acinetobacter baumannii AB307-0294 and an isogenic kanamycin-resistant Δvfr mutant. Strains were mixed at an approximately 1:1 ratio and grown for 20 h in ID-CAMHB containing 0, 0.062, 0.125 or 0.250 µg/mL FDC. The proportion of the Δvfr mutant was determined by plating each competition culture onto non-selective agar and patching 50 colonies onto kanamycin-containing agar to differentiate the mutant from the kanamycin-sensitive WT strain. Points represent independent biological replicates; bars show the mean and error bars indicate. Statistical significance was determined by one-way ANOVA with Tukey’s multiple-comparisons test: ns, not significant; ∗ P ≤ 0.05, ∗∗ P ≤ 0.01, ∗∗∗ P ≤ 0.001and ∗∗∗∗ P ≤ 0.0001.

## Discussion

Cefiderocol represents a promising last-line antibiotic against carbapenem-resistant Gram-negative bacteria, including CRAB^7,10,51,52^. However, trials such as CREDIBLE-CR reported high treatment failure and mortality in patients with CRAB infections despite apparent *in vitro* susceptibility^53,54^. This discrepancy suggests that clinically relevant adaptive resistance may arise under conditions not captured by conventional susceptibility workflows. Here, we show that cefiderocol resistance in *A. baumannii* can emerge rapidly through mutations in two global regulators, CavA and Hfq, which remodel cAMP signalling and post-transcriptional control of iron homeostasis, respectively. These changes imposed minimal fitness costs *in vitro* and *in vivo*, expanding the cefiderocol resistance paradigm beyond canonical β-lactamase-mediated resistance, porin alterations, and loss of individual TonB-dependent receptors.

CavA F237V emerged as an early mutation and conferred a moderate increase in cefiderocol MIC, but its effects were predominantly regulatory. Reduced cAMP production was accompanied by increased *pgaABCD* expression, elevated PgaAB and Csu pili abundance, enhanced biofilm biomass, and impaired twitching motility, consistent with a transition towards a more adherent, EPS-rich phenotype that may reduce cefiderocol penetration and promote antibiotic tolerance^55^. These findings align with prior reports linking cAMP–Vfr signalling, Csu pili and biofilm-associated tolerance to β-lactams and siderophore–antibiotics^21,56^, but here they arise following selection of a defined CavA variant during cefiderocol exposure. Notably, CavA F237V was identified in recent (2019–2024) outbreak-associated lineages (MLST78, MLST79 and MLST499) from the U.S. and China, indicating an epidemiologically relevant determinant beyond experimental evolution.

Hfq truncation, involving deletion of five of seven glycine-rich C-terminal repeats, emerged as a second-step mutation and drove high-level cefiderocol resistance. The truncated variant increased *adeIJK* and *ampC* expression while reducing the abundance of TonB-dependent receptors (PfeA and PiuC), together with components of acinetobactin and other siderophore uptake systems. Thus, Hfq truncation simultaneously enhances efflux and constrains ferric-siderophore import, broadening the established paradigm from loss of individual receptors (e.g., PiuA) to coordinated disruption of the broader iron-uptake network^33^. Hfq-CLASH in AB307 and iModulon analysis in AB5075 further implicated Hfq–sRNA97-mediated repression of *fur* as a post-transcriptional node connecting Hfq to iron homestasis. Fur is itself regulated post-translationally by its Fe²⁺ cofactor and autoregulates its own transcript, providing additional layers of control over Fur-mediated iron acquisition ^57,58^. Our data are therefore consistent with a model in which Hfq truncation perturbs post-transcriptional regulation of the Fur network, altering the balance between iron acquisition and iron conservation under iron limitation. Passaging of 2-repeat Hfq isolates in ID-CAMHB containing cefiderocol generated resistant derivatives without detectable chromosomal mutations, supporting a non-genetic, iron-responsive adaptive state that is compatible with heteroresistance^59,60^

A central finding of this study is that measured cefiderocol susceptibility in isolates carrying truncated-Hfq depends not only on the composition of the test medium, but also on the iron status of the medium used to prepare the inoculum. CLSI specifies determination of cefiderocol MICs in ID-CAMHB to promote siderophore-mediated drug uptake but does not standardise the iron conditions under which the inoculum is propagated before testing. Consistent with the altered iron-uptake state identified in the Hfq mutants, induction of *bauB* and *bauE* in ID-CAMHB was significantly attenuated in 2-repeat Hfq isolates, while PfeA and PiuC abundance were reduced at the protein level. Because cefiderocol exploits bacterial siderophore uptake pathways for entry, reduced expression of these systems provides a plausible mechanism for decreased drug uptake and increased MIC. We therefore define this as a defence-primed state in which prior iron restriction, in the context of Hfq truncation, remodels iron-acquisition pathways before cefiderocol exposure. Canonical seven-repeat isolates experienced the same iron restriction without exhibiting this ‘primed-state’ phenotype, indicating that the response is conditional on Hfq genotype. Because iron limitation is a characteristic of host nutritional immunity, these findings raise the possibility that standard susceptibility testing may underestimate cefiderocol resistance in isolates capable of entering this state, providing a potential mechanistic explanation for treatment failure despite *in vitro* susceptibility (Fig S5).

Truncated Hfq variants were identified in approximately 5% of available *A. baumannii* genomes, and the 2-repeat Hfq isolates with priming-dependent resistance also harbour *bla*_OXA-23_ and *bla*_OXA-66_, linking the Hfq genotype and β-lactamase resistance in clinically relevant lineages^33,60^. Hfq-dependent control of iron uptake and efflux therefore defines a lineage-specific regulatory landscape that can support or resist cefiderocol activity, helping reconcile reports of rapid on-therapy MIC escalation and heteroresistant subpopulations in CRAB with the often heterogeneous and non-conserved resistance mutations observed across isolates. Importantly, the prevelance of truncated Hfq variants in public genomes does not by itself establish their contribution to cefiderocol resistance in patients but identifies a potentially tractable genetic marker for further surveillance and phenotype investigation.

The involvement of these pathways beyond *A. baumannii* was supported by TraDIS experiments in *E. coli* MS7163. Cefiderocol selection enriched insertions in *cyaA* (*cavA* orthologue) and *crp* (*vrf* orthologue), as well as in iron-related loci (*fecA*, *tonB* and *envZ)*. No *hfq* insertions were recovered, consistent with the absence of the glycine-rich C-terminal characteristic of *A.* baumannii Hfq and suggesting that the cAMP-centred component of the phenotype may be more broadly conserved, whereas Hfq-dependent mechanism is more species-specific. A Tn5 insertion typically abolishes function, whereas the *A. baumannii* truncated Hfq variant likely retains core chaperone activity while selectively altering interaction with various sRNAs subsets. Functionally, the *E. coli* Δ*cyaA* mutant showed no change in cefiderocol MIC in single-strain assays yet outcompeted the WT strain when exposed to sub-MIC cefiderocol, indicating a fitness advantage under selective pressure. Similarly, a *Δvfr* mutant of *A. baumannii*, previously reported to phenocopy disruption of cAMP synthesis ^21^, demonstrated a strong competitive advantage over WT under sub-MIC cefiderocol but was not appreciably counter-selected in the absence of drug. These findings suggest that disruption of cAMP–Vfr signalling can alter competitive fitness specifically during cefiderocol exposure.

Together, our findings demonstrate that cefiderocol resistance in *A. baumannii* can arise through disruption of the global regulatory circuits involving CavA–cAMP signalling and Hfq-dependent control of iron homeostasis. We propose that the resulting phenotype is likely conditional on both iron availability and regulatory genotype, revealing an environmental dimension to cefiderocol susceptibility that is not captured by conventional MIC workflows. These findings support further evaluation of iron-modulated MIC testing, population analysis of isolates carrying potentially high-risk *cavA* or *hfq* alleles, and surveillance of these regulatory determinants in CRAB. Such approaches together with investigation of combination therapy in high-risk patients, may help preserve the clinical utility of cefiderocol.

## Materials and Methods

### Reagents, bacterial strains and culture conditions

Cefiderocol was purchased from NovaChem (cat. no. CS-0016784). All *A. baumannii* and *E. coli* strains employed in the study (Table S12) were grown in iron-depleted cation-adjusted Mueller-Hinton broth (ID-CAMHB) unless stated otherwise. CA-MHB (cat. no. 212322) was sourced from Becton Dickson and depletion of iron was carried out as per CLSI and EUCAST guidelines. Briefly, iron was removed from the prepared medium using Chele× 100 resin (Sigma-Aldrich, cat. no. C7901) with stirring for 2 h, followed by filtration to remove the resin. Non-iron cations (Ca²⁺, Mg²⁺, Zn²⁺) were supplemented to standard levels (Ca 20–25 mg/L, Mg 10–12.5 mg/L, Zn 0.5–1.0 mg/L). The ID-CAMHB media was then filtered sterilised and iron depletion to ≤0.03 μg/mL was confirmed by inductively coupled plasma mass spectrometry as previously described^61^. Bacterial colony forming units (CFU) enumeration was carried out on Luria Bertani broth (LB) agar. Bacteria were grown routinely at 37°C under aerobic conditions.

### Antimicrobial susceptibility testing

MICs and MIC breakpoints were determined by broth microdilution in ID-CAMHB in accordance with CLSI and EUCAST guidelines for cefiderocol susceptibility testing^19,62^. MIC assays were carried out using Nunclon Delta Surface 96-well microtiter plates in biological triplicate.

### Resistance development studies

Selection of cefiderocol resistance in AB5075 was undertaken as previously described. Briefly, all *A. baumannii* isolates were sequentially passaged in serially diluted concentrations of cefiderocol over a period of 13-days in ID-CAMHB. The highest cefiderocol concentration that demonstrated growth after each daily passage was diluted 1:250 into a new microtiter plate containing 2-fold dilutions of cefiderocol. Where resistance was observed to increase with each sequential passage, a minimum of 5 colonies were isolated for individual genome sequencing analysis.

### Genome sequencing analysis

DNA was extracted from overnight cultures of AB5075 WT or AB5075 mutants resistant to cefiderocol (5 x mutants per increase in MIC) using the DNeasy blood and tissue kit (cat. no. 69504 Qiagen) as per the manufacturer’s instructions. Paired-end Illumina DNA libraries were prepared using the Nextera DNA flex library prep kit (Illumina, Australia) with a modification of the starting input of 5 μL genomic DNA (gDNA) with samples of >20 ng/μL. WGS of pooled libraries was performed using the Illumina MiniSeq system and the high-output reagent kit (300 cycles). Illumina paired-end reads were assembled using SPAdes v3.14.1^63^ with default settings. Draft genome assemblies were ordered and orientated by aligning them to the complete genome of AB5075 using ragtag v1.1.0 ^64^. Pairwise alignments to AB5075 were visualized using the Artemis comparison tool (ACT) v18.1.0 ^65^. Whole-genome comparisons were then used to identify differences between WT and mutant AB5075 genomes. Insertion sequences were identified and annotated by a blastN comparison of draft genomes against the ISfinder database^66^. Single nucleotide polymorphism (SNP) profiling and determination of core genome SNPs was undertaken by aligning the trimmed Illumina sequencing reads for each sample to the complete genome of wild-type AB5075 (GenBank accession NZ_CP008706) using Snippy v4.4.1 with a minimum read coverage of 10× and minimum base quality of 20 that is required for a site to be considered.

### Analysis of Hfq C-terminal repeat domain copy number

To characterise the global diversity of the GGFGGQ hexapeptide tandem repeat domain in *A. baumannii* Hfq, the proteomes of 43,601 *A. baumannii* genomes were downloaded from the NCBI Protein database on 25 July 2025. Hfq was identified in each proteome by BLASTp as implemented in DIAMOND. Sequences flagged as partial in the record header were excluded from downstream analysis, yielding 39,710 full-length sequences. The copy number of the GGFGGQ hexapeptide motif in each sequence was determined by non-overlapping exact-match counting implemented in Python. The resulting copy-number distribution was visualised as a frequency histogram using matplotlib.

### Biofilm quantification

All *A. baumannii* isolates, including cefiderocol-resistant mutants, were cultured in sterile, high-binding 96-well polystyrene microtiter plates (Corning, cat. no. CLS3598). Overnight cultures were diluted in freshly prepared ID-CAMHB to an initial optical density of 0.1 at 600 nm (OD_600_ = 0.1) and incubated statically for 24 h at 37°C to allow biofilm formation. Biofilms were gently washed twice with phosphate-buffered saline (PBS, pH 7.4) to remove non-adherent cells. Adherent biofilms were stained with 0.1% (w/v) crystal violet for 15 min at room temperature, protected from light. Excess stain was removed by washing the wells three times with PBS, followed by air-drying. Crystal violet was solubilized with 95% ethanol (100 μL per well), and the resulting solution was transferred to a fresh microtiter plate and diluted 1:1 with sterile distilled H_2_O. Absorbance was measured at 570 nm with a microplate spectrophotometer to quantify biofilm biomass.

### Twitching motility

All strains were cultured overnight on LB agar. Sterile twitching medium (10 g/L tryptone, 5 g/L yeast extract, 10 g/L agar) was prepared, and 10 mL was poured into 90 mm Petri dishes and allowed to solidify and dry for 8 min. Three colonies from an overnight culture were stabbed through the agar to inoculate the plastic–medium interface, with inoculation sites evenly spaced across the plate. One plate was prepared per strain. Plates were incubated at 37°C for 24 h, after which the diameter of the twitching motility zone was measured. All twitching motility assays were performed in biological triplicate.

### Quantification of intracellular cyclic-AMP

Intracellular cAMP concentrations were determined as previously described^21^, with the following modifications. 15 mL bacterial cultures were grown to OD_600_ 0.7 and collected via centrifugation. Pellets were resuspended in 1mL PBS, transferred to lysing matrix B tubes, and lysed using a FastPrep-24 5G Homogeniser (MP Biomedicals). Bacterial cell lysates were centrifuged to remove cell debris, and the supernatant collected. cAMP was measured using the Cyclic AMP XP Assay Kit (Cell Signalling Technology) following the manufacturer’s protocol. cAMP concentrations were normalised to total protein content, determined via BCA assay (Thermo Fisher Scientific). All assays were performed in biological triplicate.

### Murine pulmonary infection model

For pulmonary infection, cohorts of 12-week-old sex-matched BALB/c mice were challenged intranasally with a 25 μL preparation of 1 × 10^8^ CFU of mid-log-phase *A. baumannii* AB5075 or mutant *A. baumannii* AB5075*^−cavA-hfq^*resistant to cefiderocol. For treatments, mice cohorts were treated with cefiderocol (10 mg/kg and 15 mg/kg intraperitoneal injection), or saline solution (0.9% [wt/vol]; intraperitoneal injection) in a 100 μL volume. Treatments were administered at 0 and 6 h postinfection. At 24 h postinfection, mice were euthanized, and the lungs were harvested and washed in sterile PBS. To determine viable bacteria in the lungs, tissue was homogenized and plated as described previously^61^.

### Transcriptome analysis

RNA was isolated using a RNeasy Plus kit (Qiagen) as described previously^67^. RNA-seq analysis was performed at the Australian Genome Research Facility. The cDNA library was prepared and assessed as described previously^67^. An average of 30 million reads per sample were generated, then trimmed using trimmomatic (v0.36), and mapped to the reference genome (*A. baumannii* AB5075 accession number NZ_CP008706) using bowtie2 (v2.3.4.2) with default parameters (–local –very-sensitive-local). Read counts were determined using featureCounts (Rsubread v1.28.1), and differential gene expression was analyzed with edgeR (v3.20.9) and limma-voom (v3.34.9) with the counts-per-million (cpm) threshold set at 1.0. Figures were generated using ggplot2 (v3.3.0) and ggrepel (v0.8.2) packages in Rstudio.

### Quantitative real-time PCR

Quantitative real-time PCR was carried out as described previously^61^. Relative gene expression was calculated by the threshold cycle (ΔΔCT) method using *gyrA* as the reference gene for all *A. baumannii* strains. All experiments were done in biological triplicates and measured in technical triplicates. Primers used for quantitative real-time PCR are provided in Table S13.

### Proteome analysis

The protocol for bacterial lysis was adapted from Tkalec et al 2024^68^. Briefly, frozen *A. baumannii* cell pellets (three biological replicates per sample) were lysed by boiling in lysis buffer (5% sodium dodecyl sulfate (BioRad), 100 mM Tris, pH 8.5) at 95°C with shaking at 750 RPM for 10 min. Samples were cooled on ice for 10 min and then subjected to a second round of boiling with shaking for 10 min. After cooling on ice, 100U Benzonase (Merck) per 50 µg protein were added to the sample and incubated for 10 min at room temperature. Protein pellets by adding nine volumes of ice-cold acetone (Sigma-Aldrich) to one volume of lysate, incubating overnight at -30°C and subsequent centrifugation at 15000 x *g* for 30 min at 4°C. Protein precipitates were then resolubilised in lysis buffer. Protein concentrations were determined by bicinchoninic acid assays (Thermo Fisher Scientific). Approximately 50 µg of protein per biological replicate were prepared for proteolytic digestion upon an S-Trap^TM^ 96-well Mini Plate ( Protifi). Initially, samples were reduced with 25 mM dithiothreitol (Roche) for 60 min at 60°C and then alkylated with 50 mM iodoacetamide (Merck) for 30 min in the dark. Samples were acidified with 1.2% phosphoric acid (Supelco) and diluted with seven sample volumes of S-Trap^TM^ wash buffer (90% methanol, 100mM triethylammonium bicarbonate (TEAB, Sigma-Aldrich), pH 8.0) and then loaded into the S-Trap^TM^ wells and washed thrice with 150 µL of S-Trap^TM^ wash buffer. Proteins were digested with 125 µL digestion buffer comprising 1.25 µg MS-grade Trypsin/LysC (Promega; 1:40, enzyme: protein ratio) in 50 mM TEAB, pH 8.0, overnight at 37°C. Peptides were then sequentially eluted with 80 µL of 50 mM TEAB, 0.2% formic acid in MS-grade water, and 50% acetonitrile. Eluates were pooled, dried under vacuum, and further desalted with C18 stage tips prepared in-house. After drying, samples were resuspended in MS grade water and 1 µL iRT (Biognosys AG) were added for LC–MS/MS analysis.

Digested proteins were analysed on a Vanquish Neo nanoLC coupled to an Orbitrap Eclipse Tribrid mass spectrometer with a FAIMS Pro Duo interface (Thermo Fisher Scientific). Peptides were chromatographically separated on a PicoChip® High Performance column (25 cm, 1.9 µm, 120 Å C18-AQ, NewObjective) over a 65-min gradient of buffer A (0.1% formic acid) and buffer B (80% acetonitrile with 0.1% formic acid) at a flow rate of 300 nL/min. Data acquisition was performed in Data-Independent Acquisition (DIA) mode utilizing the FAIMS Pro Duo interface set to a constant compensation voltage of −45 V. MS1 spectra were acquired at 60,000 resolution with a scan range from *m/z* 375 to 1500, normalised AGC target of 300%, and maximum injection time of 120 ms. For MS2 analysis, DIA scans were recorded at a resolution of 30,000 and an AGC target of 100%, with a maximum injection time of 54 ms. Fragmentation was induced with stepped HCD of collision energies of 25%, 30%, and 35%. Peptide data were searched in Spectronaut version 18.6.231227.55695 (Biognosys AG) by directDIA (library-free) analysis against the *Acinetobacter baumannii* UniProt database (UP000032746) with the following modifications: maximum of 2 missed cleavages; cysteine carbamidomethylation (fixed), and methionine oxidation (variable). Data were filtered at a false discovery rate (FDR) of 0.01 at the protein level.

### CLASH sequencing

The chromosomal copy of *hfq* in *A. baumannii* strain AB307-0294 *wzc*::*aph* was tagged with HTF (His_6_-TEV-FLAG). Hfq-CLASH was performed as described previously (PMID: 37925184) using iron-replete and iron-depleted LB media. For a detailed description of *A. baumannii* Hfq-CLASH refer to the Supplementary Methods. Briefly, *A*. *baumannii* AB307_0294 *wzc*::*aph* and AB307_0294 *wzc*::*aph hfq-*HTF were used to inoculate iron-repleted and depleted LB media and were grown with shaking at 37°C, 200 rpm to an OD_600_ of 0.6 prior to crosslinking by UV irradiation and immediate harvesting. Cells were lysed and debris clarified by centrifugation. Supernatants were added to an anti-FLAG M2 affinity gel (Sigma) and incubated at 4°C for 2 hr with gentle rotation. Hfq-RNA complexes were washed and the FLAG tags were cleaved by TEV protease digestion at 18°C with gentle rotation for 2 hr. Cleaved Hfq-RNA complexes were then treated with 0.15 U of RNace-IT (Agilent) for 7 min at 20°C to trim RNA. The Hfq-RNA complexes were added to pre-equilibrated magnetic Ni-NTA beads and incubated at 4°C with gentle rotation overnight. Following incubation with Ni-NTA beads, the Hfq-RNA were washed, and the 5′ RNA phosphate groups were removed by addition of 1 U of Calf Intestinal Alkaline Phosphatase (CIAP, Promega). RNA was radiolabelled by addition of 30 µCi of [γ-³²P]-ATP (Revvity Pty Ltd). Linker aptamers were ligated to the 3′ end using App-PE oligonucleotide (Integrated DNA Technologies), and to the 5′ end using unique barcoded index, and 40 U of T4 ssRNA ligase I (NEB). ^32^P-labelled Hfq-RNA complexes were eluted and resolved on NuPAGE 4–12% Bis-Tris gel (Invitrogen). Radiolabelled Hfq-RNA complexes were visualised by autoradiography and bands corresponding to Hfq–RNA were excised, collected, and digested with 100 µg Proteinase K (NEB) at 55°C for 2 hr with mixing. RNA was extracted with 3 M NaOAc and phenol:chloroform:isoamyl alcohol, followed by ethanol precipitation. cDNA libraries were constructed by reverse transcription and PCR amplification. PCR products were resolved on a 2% MetaPhor agarose gel (Lonza), and amplicons were excised and purified using a MinElute Gel Extraction Kit (Qiagen). Library concentrations were determined using the Qubit dsDNA assay (Thermo Fisher), pooled to equimolar concentrations, and sequenced on an Illumina NovaSeq X Plus platform. Data were analysed using the Hyb-CRAC-R pipeline ^69^ that incorporates pyCRAC software ^70^ for single read mapping and *hyb* software ^71^ for hybrid read mapping. Sequence data have been deposited at NCBI GEO under accession number GSE319194.

### iModulon Analysis and Code Availability

We utilized statistically independent components (M) previously identified through independent component analysis (ICA), along with normalized log TPM values (X) and their associated activity profiles (A) across various experimental conditions, all of which were obtained from iModulonDB ^41,42^.

In this study, we further incorporated newly obtained normalized log TPM values (X) and the corresponding activity profiles (A), calculated from RNA-seq count data for wild-type AB5075, AB5075*^cavA^*, and AB5075*^cavA-hfq^*, all grown in ID-CAMHB. Details of these combined datasets are provided in Supplementary Data 6 and 7. The Python package for analysing and visualising iModulons (PyModulon) is also available on GitHub (https://github.com/SBRG/pymodulon).

### Transposon-directed insertion-site sequencing

A total of 4736 cefiderocol-resistant transposon mutants were pooled, and DNA samples were extracted for transposon-directed insertion-site sequencing (TraDIS) analysis as previously described^72^. In brief, the Illumina library preparation was performed using the Nextera Flex DNA Sample Prep Kit (Illumina) following the manufacturer’s instructions with modifications for TraDIS and sequenced with 100 bp single-end reads on the Illumina MiSeq platform.

Spontaneous resistant mutants were sequenced using Illumina technology. The sequencing libraries were prepared using the Nextera DNA Library Prep Kit. The libraries were sequenced with 2 × 100 bp paired-end reads using the Illumina HiSeq 2500. Illumina reads were analysed using an in-house pipeline for quality control. High quality Illumina reads were mapped to the EC958 genome (HG941718) using SHRIMP 2.0 and nucleotide variations were identified using Nesoni (www.vicbioinformatics.com/software.nesoni.shtml)^73^.

### Growth competition assays

#### Escherichia coli

Fitness competitions were performed using a broth microdilution MIC-based protocol. For each competition, wild-type (WT) and mutant strains were combined at a 1:1 ratio to generate the initial inoculum, which was then grown in the presence of four concentrations of cefotaxime (Cef): 0 (no antibiotic), 0.25× MIC, 0.5× MIC, and 1× MIC, corresponding to 0, 0.03125, 0.0625, and 0.125 µg/mL Cef, respectively, for *E. coli*.

Because the WT strain is chloramphenicol (Cm)-sensitive whereas the mutant carries a Cm-resistance marker (CmR), the relative proportion of each strain within the competition was assessed by differential plating. On day 3 of the competition, aliquots were plated onto non-selective LB agar (total viable count) and onto LB agar supplemented with 30 µg/mL chloramphenicol (LB-Cm30; CmR count). The percentage of CmR colonies was then calculated as the number of colonies recovered on LB-Cm30 relative to the total number recovered on LB.

Competitions were performed for the following strain pairings: WT vs. WT(CmR) (fitness-neutral control), WT vs. mutant, WT vs. mutant carrying the empty vector control, and WT vs. mutant complemented with pcyaA.

#### Acinetobacter baumannii

Fitness competitions were performed in ID-CAMHB using *A. baumannii* AB307-0294 as the wild-type (WT) strain and an isogenic kanamycin-resistant Δ*vfr* mutant. For each competition, WT and Δ*vfr* strains were combined at an approximately 1:1 ratio and incubated for 20 h in the absence of cefiderocol or in the presence of 0.062, 0.125 or 0.250 µg/mL cefiderocol, 0.25× MIC, 0.5× MIC, and 1× MIC, respectively. Competitions were performed with three independent biological replicates.

After 20 h, cultures were serially diluted and plated onto non-selective LB agar. Following incubation, 50–100 colonies from each biological replicate and condition were patched onto non-selective LB agar and LB agar supplemented with kanamycin to differentiate the kanamycin-resistant Δ*vfr* mutant from the kanamycin-sensitive WT strain. The proportion of the Δ*vfr* mutant in each competition was calculated as the number of kanamycin-resistant colonies divided by the total number of colonies patched.

### Ethics

Animal experiments were performed according to the Australian code of practice for the care and use of animals for scientific purposes. Permission was obtained from the University of Queensland ethics committee (2022/AE000901).

### Statistical analysis

All experiments were undertaken either in technical duplicate or triplicate, with no less than 3 biological replicates. Data are presented as means ± SD. Bacterial dissemination data from *in vivo* studies are presented as geometric means. To compare means between more than two groups, a one-way ANOVA with *post hoc* (Tukey) was conducted. Statistical analysis was performed using GraphPad Prism software v9. A *p*-value of <0.05 was considered statistically significant.

## Data availability

Genome, RNA-seq and proteomic data sets have been deposited to NCBI under BioProject PRJNA1406814 and the ProteomeXchange repository (PXD073926). Raw Illumina sequence read data have been deposited to the Sequence Read Archive (SRA) under the accessions SRR36996861 to SRR36996895.

