## Supplementary material for "Disruption of cAMP Signalling and RNA Regulation Confers Cefiderocol Resistance in *Acinetobacter baumannii*": Table S

**Supplementary data**

**Supplementary tables**

***Table S1.*** *AB5075 mutant chromosomal differences as identified by Illumina whole genome sequencing.*

| **Locus_tag** | **Gene; Product** | **Mutation** |
| --- | --- | --- |
| ABUW_2208 | *cavA;* adenylate/guanylate cyclase domain-containing protein | F237V |
| ABUW_1609 | *hfq*; RNA chaperone | ΔG117→G132 |

***Table S2a. RNAseq AB5075^-cavA^ up-regulated genes relative to AB5075 WT.*** *Strains were cultured to OD_600nm_=0.5 in ID-CAMHB at 37ºC.*

| **Locus_tag** | **Log_2_FC** | **Pvalue** | **Gene** | **Product** |
| --- | --- | --- | --- | --- |
| ABUW_1560 | 1.577 | 0.00025027 | *pgaD* | Poly‑β‑1‑6‑N‑acetyl‑D‑glucosamine biosynthesis |
| ABUW_0687 | 1.362 | 0.00194423 | *ecnB* | Entericidin EcnB |
| ABUW_1439 | 1.312 | 2.84E-14 | *-* | Hypothetical protein |
| ABUW_1557 | 1.302 | 0.00024463 | *pgaA* | Outer membrane protein |
| ABUW_2216 | 1.271 | 2.36E-05 | *rpsT* | Ribosomal protein S20 |
| ABUW_1559 | 1.213 | 0.00055076 | *pgaC* | Glycosyl transferase, family 2 protein |
| ABUW_1558 | 1.198 | 0.00037606 | *pgaB* | Polysaccharide deacetylase |
| ABUW_0237 | 1.139 | 0.01690323 | *aqpZ* | Aquaporin Z |
| ABUW_1077 | 1.118 | 0.00395625 | - | Hypothetical protein |
| ABUW_2704 | 1.108 | 0.00071267 | - | Hypothetical protein |
| ABUW_2103 | 1.091 | 0.02417579 | - | Hypothetical protein |
| ABUW_3881 | 1.028 | 0.01849047 | - | Matrixin superfamily |
| ABUW_1561 | 1.000 | 0.00111767 | - | Hypothetical protein |

***Table S2b. RNAseq AB5075^-cavA^ down-regulated genes relative to AB5075 WT.*** *Strains were cultured to OD_600nm_=0.5 in ID-CAMHB at 37ºC.*

| **Locus_tag** | **Log_2_FC** | **Pvalue** | **Gene** | **Product** |
| --- | --- | --- | --- | --- |
| ABUW_2707 | -1.023 | 0.00296563 | *actP2* | Copper-translocating P-type ATPase |
| ABUW_0779 | -1.036 | 0.00351332 | *-* | Hypothetical protein |
| ABUW_0785 | -1.042 | 0.00115052 | *proB* | Glutamate-5-kinase |
| ABUW_1793 | -1.060 | 0.01131363 | *ybgT* | Cyd operon protein YbgT |
| ABUW_3789 | -1.094 | 0.01100455 | *dadA2* | D-amino acid dehydrogenase small subunit |
| ABUW_2338 | -1.104 | 1.15E-05 | *metQ1* | D-methionine transport protein |
| ABUW_2380 | -1.134 | 0.00607997 | *tauC* | Taurine transport system permease protein |
| ABUW_1018 | -1.158 | 0.0040161 | *cysW* | Sulfate ABC transporter, permease protein CysW |
| ABUW_2421 | -1.166 | 0.00398996 | *-* | ABC transporter, periplasmic binding protein |
| ABUW_1019 | -1.169 | 0.00903981 | *cysT* | Sulfate ABC transporter, permease protein cyst |
| ABUW_1020 | -1.241 | 0.00362179 | *-* | Alpha/beta hydrolase fold protein |
| ABUW_2420 | -1.273 | 0.00185141 | *-* | ABC transporter, periplasmic binding protein |
| ABUW_2379 | -1.289 | 0.0022245 | *tauD* | Taurine dioxygenase |
| ABUW_1795 | -1.296 | 0.00194528 | *cydA* | Cytochrome D ubiquinol oxidase subunit I |
| ABUW_0449 | -1.304 | 6.51E-10 | *-* | Hypothetical protein |
| ABUW_2381 | -1.376 | 0.00169785 | *tauB* | Taurine import ATP-binding protein |
| ABUW_1021 | -1.392 | 0.00064817 | *cysP* | Thiosulfate-binding protein |
| ABUW_3583 | -1.452 | 0.00010813 | *-* | Putative outer membrane protein W |
| ABUW_2382 | -1.548 | 0.0007354 | *tauA* | Taurine ABC transporter, periplasmic binding protein |

***Table S3a. RNAseq AB5075^-cavA-hfq^ up-regulated genes relative to AB5075^-cavA^.*** *Strains were cultured to OD_600nm_=0.5 in ID-CAMHB at 37ºC.*

| **Locus_tag** | **Log_2_FC** | **Pvalue** | **Gene** | **Product** |
| --- | --- | --- | --- | --- |
| ABUW_1792 | 5.554 | 6.46E-34 | *-* | Hypothetical protein |
| ABUW_1795 | 5.483 | 3.71E-33 | *cydA* | Cytochrome D ubiquinol oxidase subunit I |
| ABUW_1793 | 5.245 | 3.37E-30 | *ybgT* | Cyd operon protein YbgT |
| ABUW_1794 | 5.037 | 2.24E-30 | *cydB* | Cytochrome D ubiquinol oxidase, subunit II |
| ABUW_3583 | 4.005 | 3.02E-15 | *-* | Putative outer membrane protein W |
| ABUW_1796 | 3.858 | 3.65E-19 | *-* | Hypothetical protein |
| ABUW_1345 | 3.575 | 7.28E-13 | *-* | Peptidase U32 |
| ABUW_3508 | 3.452 | 6.98E-22 | *-* | Hypothetical protein |
| ABUW_1536 | 3.350 | 2.83E-10 | *-* | Hypothetical protein |
| ABUW_1943 | 3.001 | 3.53E-12 | *-* | Nicotinamide phosphoribosyltransferase |
| ABUW_2435 | 2.992 | 7.98E-09 | *-* | Oxidoreductase |
| ABUW_2065 | 2.959 | 8.93E-09 | *-* | Hypothetical protein |
| ABUW_1942 | 2.950 | 2.81E-13 | *-* | Ribose-phosphate pyrophosphokinase |
| ABUW_3125 | 2.934 | 1.44E-21 | *bfr2* | Bacterioferritin |
| ABUW_1466 | 2.916 | 7.75E-08 | *-* | Hypothetical protein |
| ABUW_0377 | 2.846 | 3.23E-17 | *hemF* | Coproporphyrinogen III oxidase, aerobic |
| ABUW_2436 | 2.775 | 6.63E-09 | *katE* | Catalase |
| ABUW_0672 | 2.675 | 9.54E-07 | *-* | Transposase inhibitor |
| ABUW_1227 | 2.673 | 5.13E-07 | *-* | Acyl-coa dehydrogenase |
| ABUW_3789 | 2.666 | 1.60E-09 | *dadA2* | D-amino acid dehydrogenase small subunit |
| ABUW_1471 | 2.640 | 3.26E-06 | *-* | Hypothetical protein |
| ABUW_2060 | 2.636 | 2.94E-08 | *-* | Hypothetical protein |
| ABUW_2064 | 2.604 | 5.48E-07 | *-* | Hypothetical protein |
| ABUW_1468 | 2.587 | 5.42E-07 | *-* | Lmbe-like protein |
| ABUW_0671 | 2.520 | 3.76E-18 | *-* | Transposase |
| ABUW_1469 | 2.515 | 2.61E-06 | *-* | Methyltransferase type 12 |
| ABUW_1344 | 2.468 | 3.05E-08 | *-* | Iron-sulfur cluster-binding protein |
| ABUW_1661 | 2.426 | 1.01E-15 | *-* | Universal stress protein family |
| ABUW_1467 | 2.401 | 4.23E-07 | *-* | Acyl-coa dehydrogenase |
| ABUW_2707 | 2.394 | 4.98E-12 | *actP2* | Copper-translocating P-type atpase |
| ABUW_0562 | 2.369 | 3.08E-06 | *-* | Transposase inhibitor |
| ABUW_2439 | 2.359 | 1.36E-05 | *-* | Hypothetical protein |
| ABUW_1526 | 2.353 | 6.62E-08 | *-* | Hypothetical protein |
| ABUW_3346 | 2.315 | 2.51E-10 | *acnA* | Aconitate hydratase 1 |
| ABUW_0306 | 2.294 | 1.92E-18 | *bfr1* | Bacterioferritin |
| ABUW_2540 | 2.258 | 3.81E-06 | *-* | Transposase |
| ABUW_3147 | 2.242 | 4.52E-09 | *-* | Hypothetical protein |
| ABUW_3788 | 2.212 | 3.03E-07 | *dadX* | Alanine racemase |
| ABUW_2440 | 2.194 | 3.87E-05 | - | Surface antigen |
| ABUW_2458 | 2.191 | 1.04E-10 | - | Oxidoreductase |
| ABUW_3397 | 2.184 | 1.46E-29 | *fumA* | Fumarate hydratase |
| ABUW_3429 | 2.182 | 8.76E-09 | *calB* | Aldehyde dehydrogenase |
| ABUW_3157 | 2.106 | 9.22E-06 | - | Hypothetical protein |
| ABUW_2708 | 2.084 | 9.35E-08 | - | Heavy metal transport/detoxification protein |
| ABUW_0255 | 2.080 | 2.92E-21 | - | Succinyl-coa:coenzyme A transferase |
| ABUW_2730 | 2.072 | 3.70E-07 | - | Ompa/motb |
| ABUW_1194 | 2.058 | 2.58E-09 | *ampC* | Beta-lactamase ADC7 |
| ABUW_3123 | 2.022 | 1.26E-07 | *otsA* | Trehalose-6-phosphate synthase |
| ABUW_0077 | 2.019 | 6.22E-07 | *hutU* | Urocanate hydratase |
| ABUW_3814 | 2.016 | 1.81E-10 | *lldP* | L-lactate permease |
| ABUW_2702 | 2.010 | 2.36E-05 | - | Hypothetical protein |
| ABUW_0078 | 2.002 | 5.13E-10 | *hutH* | Histidine ammonia-lyase |
| ABUW_3787 | 1.972 | 2.70E-06 | - | Endoribonuclease L-PSP family protein |
| ABUW_2433 | 1.954 | 1.18E-08 | - | KGG domain-containing protein |
| ABUW_2606 | 1.951 | 1.12E-07 | - | Glycoside hydrolase/deacetylase family protein |
| ABUW_3037 | 1.949 | 0.00019192 | - | Hemerythrin/HHE cation-binding motif- containing protein |
| ABUW_3582 | 1.948 | 1.98E-08 | - | Hypothetical protein |
| ABUW_1726 | 1.922 | 1.15E-08 | - | D-amino acid dehydrogenase 3 small subunit |
| ABUW_1787 | 1.918 | 2.03E-09 | - | Hypothetical protein |
| ABUW_2451 | 1.901 | 3.10E-23 | - | Transcriptional regulator acrr family |
| ABUW_3812 | 1.850 | 6.48E-12 | *lldD* | L-lactate dehydrogenase (cytochrome) |
| ABUW_2621 | 1.839 | 4.92E-07 | - | Hypothetical protein |
| ABUW_1216 | 1.825 | 1.05E-16 | *sodB* | Superoxide dismutase (Fe) |
| ABUW_0184 | 1.811 | 6.57E-12 | - | Na+/solute symporter |
| ABUW_2737 | 1.797 | 4.47E-05 | *clpB* | ATP-dependent chaperone ClpB |
| ABUW_0070 | 1.793 | 1.44E-06 | *fahA* | Fumarylacetoacetase |
| ABUW_1470 | 1.792 | 0.00012009 | - | Glycosyl transferase, family 2 |
| ABUW_0328 | 1.792 | 1.06E-05 | - | Hemerythrin |
| ABUW_1488 | 1.782 | 1.73E-08 | *csuA* | CsuA |
| ABUW_0843 | 1.778 | 7.16E-19 | *adeJ* | Multidrug efflux protein AdeJ |
| ABUW_0944 | 1.753 | 6.10E-07 | - | Oxidoreductase alpha (molybdopterin) subunit |
| ABUW_2319 | 1.749 | 4.99E-07 | - | Putative membrane protein |
| ABUW_1721 | 1.742 | 1.02E-08 | - | UDP-glucose 4-epimerase |
| ABUW_2389 | 1.731 | 1.40E-17 | *cioA* | Cytochrome bd ubiquinol oxidase, subunit I |
| ABUW_2706 | 1.727 | 3.96E-09 | *cueR* | Cu(I)-responsive transcriptional regulator |
| ABUW_0842 | 1.724 | 1.91E-20 | *adeK* | Multidrug efflux protein adek |
| ABUW_1593 | 1.722 | 2.78E-15 | *acnB* | Aconitate hydratase 2 |
| ABUW_2430 | 1.713 | 3.83E-08 | *0* | Transporter, lyse family |
| ABUW_2130 | 1.703 | 4.78E-05 | *acoB* | Acetoin:2,6-dichlorophenolindophenol oxidoreductase subunit beta |
| ABUW_1489 | 1.699 | 3.14E-06 | *csuB* | CsuB |
| ABUW_0469 | 1.695 | 5.99E-16 | - | Hypothetical protein |
| ABUW_2607 | 1.691 | 1.85E-09 | - | Hypothetical protein |
| ABUW_3881 | 1.679 | 0.00180374 | - | Matrixin superfamily |
| ABUW_1550 | 1.666 | 4.31E-39 | *cyoD* | Cytochrome O ubiquinol oxidase subunit IV |
| ABUW_2345 | 1.655 | NA | - | Hypothetical protein |
| ABUW_3124 | 1.647 | 3.26E-19 | - | Major facilitator superfamily MFS_1 |
| ABUW_1490 | 1.644 | 1.90E-06 | *csuC* | CsuC |
| ABUW_0183 | 1.641 | 1.71E-09 | - | Hypothetical protein |
| ABUW_2131 | 1.638 | 0.0003053 | *acoA* | Acetoin:2,6-dichlorophenolindophenol oxidoreductase alpha subunit |
| ABUW_0561 | 1.631 | 1.75E-05 | - | Transposase |
| ABUW_2437 | 1.617 | 0.00010409 | - | Heme oxygenase-like protein |
| ABUW_1886 | 1.614 | 1.48E-06 | *cpo* | Alpha/beta hydrolase |
| ABUW_0053 | 1.606 | 2.31E-05 | - | UPF0391 membrane protein |
| ABUW_3158 | 1.605 | 0.00010602 | - | Hypothetical protein |
| ABUW_3395 | 1.605 | 4.27E-08 | *ackA* | Acetate kinase |
| ABUW_2379 | 1.593 | 0.00016588 | *tauD* | Taurine dioxygenase |
| ABUW_2756 | 1.590 | 1.43E-05 | - | Hypothetical protein |
| ABUW_3252 | 1.589 | 3.31E-05 | *raiA* | Ribosomal subunit interface protein |
| ABUW_2605 | 1.587 | 6.49E-05 | *0* | Hypothetical protein |
| ABUW_1464 | 1.579 | 9.80E-10 | *mmsA2* | Methylmalonate-semialdehyde dehydrogenase |
| ABUW_1343 | 1.572 | 1.67E-08 | - | Small multidrug resistance protein |
| ABUW_1111 | 1.570 | NA | - | Aldehyde dehydrogenase |
| ABUW_0844 | 1.552 | 3.04E-14 | *adeI* | Multidrug efflux protein AdeI |
| ABUW_1764 | 1.550 | 1.38E-08 | - | GGDEF family protein |
| ABUW_2452 | 1.518 | 5.09E-22 | - | Isovaleryl-coa dehydrogenase |
| ABUW_2129 | 1.516 | 0.00010673 | *acoC* | Dihydrolipoamide acetyltransferase |
| ABUW_1074 | 1.502 | 9.12E-05 | *icd* | Isocitrate dehydrogenase, NADP-dependent |
| ABUW_1226 | 1.485 | 0.0003049 | - | 4-carboxymuconolactone decarboxylase |
| ABUW_2380 | 1.482 | 0.00035436 | *tauC* | Taurine transport system permease protein |
| ABUW_0133 | 1.482 | 9.09E-05 | - | Hypothetical protein |
| ABUW_0827 | 1.479 | NA | *pmrC* | Lipid A phosphoethanolamine transferase |
| ABUW_1540 | 1.478 | 1.02E-27 | - | Hypothetical protein |
| ABUW_2387 | 1.472 | 1.50E-06 | - | Hypothetical protein |
| ABUW_3798 | 1.460 | 6.30E-06 | - | Short-chain dehydrogenase/reductase SDR |
| ABUW_1905 | 1.460 | 0.00185413 | - | Isochorismatase hydrolase |
| ABUW_1491 | 1.448 | 6.30E-05 | *csuD* | CsuD |
| ABUW_1904 | 1.445 | 0.00216356 | - | Antibiotic biosynthesis monooxygenase |
| ABUW_2453 | 1.440 | 3.12E-11 | - | Methylcrotonoyl-coa carboxylase beta chain |
| ABUW_1668 | 1.430 | 2.66E-09 | - | Benzoate transporter |
| ABUW_2390 | 1.429 | 3.23E-10 | *cioB* | Cytochrome D ubiquinol oxidase, subunit II |
| ABUW_1891 | 1.424 | 0.00049953 | *0* | Thij/pfpi domain protein |
| ABUW_3813 | 1.417 | 7.80E-08 | *lldR* | L-lactate utilization transcriptional repressor |
| ABUW_1465 | 1.408 | 1.12E-13 | - | Gamma-aminobutyrate permease |
| ABUW_3231 | 1.402 | 9.30E-30 | *rpe* | Ribulose-phosphate 3-epimerase |
| ABUW_1573 | 1.397 | 0.00019797 | - | Acyl-coa dehydrogenase |
| ABUW_2341 | 1.395 | 3.37E-07 | - | D-methionine transport system permease protein MetI |
| ABUW_1703 | 1.391 | 0.00113245 | - | Hypothetical protein |
| ABUW_2391 | 1.385 | 5.18E-09 | - | Hypothetical protein |
| ABUW_1463 | 1.368 | 6.59E-07 | - | Omega-amino acid-pyruvate aminotransferase |
| ABUW_1143 | 1.366 | 2.41E-06 | - | Hypothetical protein |
| ABUW_1763 | 1.360 | 2.05E-05 | - | Uspa domain protein |
| ABUW_1015 | 1.360 | 0.0001507 | *carO* | Outer membrane protein CarO |
| ABUW_2381 | 1.356 | 0.00198357 | *tauB* | Taurine import ATP-binding protein |
| ABUW_0068 | 1.354 | 0.00015155 | - | Glyoxalase/bleomycin resistance protein/dioxygenase |
| ABUW_3469 | 1.349 | 8.67E-06 | *katG* | Catalase/peroxidase HPI |
| ABUW_2298 | 1.348 | 2.78E-07 | - | Hypothetical protein |
| ABUW_2123 | 1.346 | 1.08E-09 | - | Metallo-beta-lactamase family protein |
| ABUW_2456 | 1.346 | 0.00043627 | - | Hydroxymethylglutaryl-coa lyase |
| ABUW_2374 | 1.340 | 2.44E-10 | - | Isochorismatase hydrolase |
| ABUW_0646 | 1.337 | 0.00010124 | - | Hypothetical protein |
| ABUW_1768 | 1.324 | 7.19E-06 | - | Transcriptional regulator, marR-family |
| ABUW_3204 | 1.319 | 0.00047712 | - | Amidohydrolase |
| ABUW_1305 | 1.314 | 0.00420886 | - | Hypothetical protein |
| ABUW_0628 | 1.309 | 0.00061842 | - | Peroxidase |
| ABUW_3502 | 1.309 | 1.17E-17 | *etfD* | Electron transfer flavoprotein-ubiquinone oxidoreductase (EsvG) |
| ABUW_2388 | 1.295 | 0.00035729 | - | Hypothetical protein |
| ABUW_2594 | 1.291 | 3.16E-05 | - | Glutathione-dependent formaldehyde dehydrogenase |
| ABUW_1761 | 1.287 | 0.00124046 | - | Hypothetical protein |
| ABUW_2887 | 1.286 | 0.00116439 | - | Lipoprotein |
| ABUW_1659 | 1.285 | 0.00372279 | - | Hypothetical protein |
| ABUW_2122 | 1.284 | 4.56E-10 | - | Oxidoreductase |
| ABUW_2072 | 1.262 | 4.54E-12 | *ppk1* | Polyphosphate kinase |
| ABUW_0298 | 1.258 | 2.15E-12 | *gltB* | Glutamate synthase, large subunit |
| ABUW_0468 | 1.256 | 0.00073359 | - | Hypothetical protein |
| ABUW_0845 | 1.252 | 1.07E-09 | - | Hypothetical protein |
| ABUW_0987 | 1.246 | 7.06E-07 | - | Hypothetical protein |
| ABUW_2128 | 1.244 | 0.00024119 | *lpdA2* | Dihydrolipoamide dehydrogenase |
| ABUW_0885 | 1.238 | 0.00159856 | - | Biofilm-associated protein |
| ABUW_2163 | 1.230 | 5.59E-40 | - | Hypothetical protein |
| ABUW_1542 | 1.228 | 3.42E-10 | - | Transglutaminase |
| ABUW_3225 | 1.227 | 1.02E-42 | *sugE* | Suppressor of Gro |
| ABUW_1879 | 1.224 | NA | - | Hypothetical protein |
| ABUW_0376 | 1.221 | 1.09E-07 | *aroE* | Shikimate 5-dehydrogenase |
| ABUW_0859 | 1.218 | 5.15E-06 | - | Hypothetical protein |
| ABUW_2850 | 1.207 | 4.92E-12 | - | Lysine exporter protein ArgO |
| ABUW_1047 | 1.204 | 5.37E-06 | *uvrB* | Excinuclease ABC, B subunit |
| ABUW_1572 | 1.202 | 2.92E-05 | - | Short-chain dehydrogenase/reductase |
| ABUW_0299 | 1.200 | 2.43E-10 | *gltD* | Glutamate synthase, small subunit |
| ABUW_1629 | 1.198 | 6.37E-06 | - | Hypothetical protein |
| ABUW_2438 | 1.193 | 0.00487175 | *cinA1* | Competence/damage-inducible protein cina |
| ABUW_2900 | 1.193 | 9.57E-13 | - | Hypothetical protein |
| ABUW_1720 | 1.191 | 1.17E-05 | *tusD* | Sulfur relay protein TusD/DsrE |
| ABUW_0518 | 1.188 | 3.79E-10 | - | Hypothetical protein |
| ABUW_3352 | 1.185 | 0.00109054 | - | Putative transcriptional regulator |
| ABUW_2145 | 1.177 | 2.45E-07 | - | Hypothetical protein |
| ABUW_2684 | 1.177 | 1.67E-11 | - | Phage putative head morphogenesis protein |
| ABUW_2443 | 1.174 | 0.00566951 | - | Hypothetical protein |
| ABUW_2705 | 1.173 | 3.08E-06 | - | Hypothetical protein |
| ABUW_0932 | 1.170 | 1.14E-07 | - | Non-ribosomal peptide synthetase |
| ABUW_1018 | 1.169 | 0.00372715 | *cysW* | Sulfate ABC transporter, permease protein CysW |
| ABUW_2092 | 1.167 | 0.00043691 | *bdhA* | 3-hydroxybutyrate dehydrogenase |
| ABUW_2800 | 1.158 | 0.00058334 | - | Succinylornithine transaminase |
| ABUW_1732 | 1.155 | 7.89E-08 | *ntrC* | Nitrogen metabolism transcriptional regulator, ntrc, Fis Family |
| ABUW_3682 | 1.150 | 0.00546654 | - | Putative dihydrodipicolinate synthase |
| ABUW_3887 | 1.143 | 1.83E-08 | - | Transmembrane signal peptide protein |
| ABUW_1159 | 1.139 | 0.00181317 | - | Transcriptional regulator, TetR family |
| ABUW_0945 | 1.134 | 0.00102554 | *fdhD* | Formate dehydrogenase accessory protein FdhD |
| ABUW_1048 | 1.134 | 3.75E-05 | - | Outer membrane lipoprotein Blc |
| ABUW_2865 | 1.133 | 1.43E-05 | *lon* | ATP-dependent protease La |
| ABUW_1487 | 1.119 | 0.01960112 | - | CsuA/B |
| ABUW_3898 | 1.118 | 0.00290024 | - | Transglycosylase-associated protein |
| ABUW_1005 | 1.115 | 9.20E-30 | - | Hypothetical protein |
| ABUW_3019 | 1.114 | 1.80E-30 | *emrB* | Multidrug resistance protein B |
| ABUW_2382 | 1.109 | 0.01462439 | *tauA* | Taurine ABC transporter, periplasmic binding protein |
| ABUW_0433 | 1.108 | 1.43E-19 | - | Flavin-containing monooxygenase FMO |
| ABUW_0722 | 1.106 | 2.45E-18 | *cysH* | Phosphoadenosine phosphosulfate reductase |
| ABUW_2603 | 1.104 | 0.00310738 | *bccA* | Carbamoyl-phosphate synthase |
| ABUW_2724 | 1.104 | 0.00478896 | - | Hypothetical protein |
| ABUW_0721 | 1.104 | 1.16E-09 | *wax-dgaT* | Bifunctional protein wax ester synthase/acyl-coa diacylglycerol acyltransferase |
| ABUW_1165 | 1.099 | 1.11E-21 | - | Pirin domain protein |
| ABUW_2957 | 1.098 | 0.00036796 | - | Enoyl-coa hydratase/isomerase |
| ABUW_2318 | 1.097 | 0.00020037 | - | Cytosine/purines/uracil/thiamine/allantoin permease family protein |
| ABUW_1065 | 1.096 | 6.30E-05 | - | Hypothetical protein |
| ABUW_1379 | 1.095 | 0.00100031 | - | Hypothetical protein |
| ABUW_1675 | 1.094 | 3.56E-05 | - | Short-chain dehydrogenase/reductase |
| ABUW_2888 | 1.092 | 1.84E-10 | *aceA* | Isocitrate lyase |
| ABUW_1733 | 1.092 | 4.49E-10 | *ntrB* | Signal transduction histidine kinase, nitrogen specific, NtrB |
| ABUW_3351 | 1.091 | 0.00369157 | - | Heme oxygenase-like protein |
| ABUW_2709 | 1.088 | 0.00068546 | *benM* | Ben operon transcriptional regulator BenM |
| ABUW_1046 | 1.087 | 2.78E-06 | - | Hypothetical protein |
| ABUW_1873 | 1.082 | 0.00369164 | *pcaJ2* | 3-oxoadipate coa-transferase subunit B |
| ABUW_2058 | 1.082 | 0.00292518 | - | Hypothetical protein |
| ABUW_2232 | 1.078 | 0.00032209 | - | Biopolymer transport protein exbd/tolr |
| ABUW_3786 | 1.076 | 0.00483639 | *cycA2* | D-serine/D-alanine/glycine transporter |
| ABUW_1019 | 1.071 | 0.01661162 | *cysT* | Sulfate ABC transporter, permease protein CysT |
| ABUW_2358 | 1.070 | 0.00013135 | - | Proton/sodium-glutamate symport protein |
| ABUW_3579 | 1.068 | 0.00150503 | - | Hypothetical protein |
| ABUW_0069 | 1.067 | 0.00293277 | *maiA* | Maleylacetoacetate isomerase |
| ABUW_1790 | 1.067 | 2.26E-07 | - | Nitroreductase family protein |
| ABUW_1839 | 1.060 | 2.12E-05 | *pcaG* | Protocatechuate 3,4-dioxygenase, alpha subunit |
| ABUW_1021 | 1.056 | 0.00941188 | *cysP* | Thiosulfate-binding protein |
| ABUW_3053 | 1.055 | 1.04E-10 | *ftsK* | DNA translocase FtsK |
| ABUW_2159 | 1.051 | 2.43E-06 | - | Histidine triad protein |
| ABUW_2604 | 1.051 | 0.01771436 | - | Allophanate hydrolase |
| ABUW_3396 | 1.048 | 0.00045273 | *pta* | Phosphate acetyltransferase |
| ABUW_1020 | 1.048 | 0.01379323 | - | Alpha/beta hydrolase fold protein |
| ABUW_2992 | 1.044 | 0.01137228 | - | Hypothetical protein |
| ABUW_3103 | 1.043 | 1.21E-06 | *ribB2* | 3,4-dihydroxy-2-butanone 4-phosphate synthase |
| ABUW_1323 | 1.042 | 0.01724183 | - | Hypothetical protein |
| ABUW_1574 | 1.042 | 0.00216612 | - | Acyl-coa ligase |
| ABUW_1158 | 1.042 | 0.00043607 | - | Hypothetical protein |
| ABUW_1654 | 1.042 | 0.01320023 | - | Acetyltransferase, gnat family |
| ABUW_1142 | 1.041 | 5.73E-05 | *cstA* | Carbon starvation protein A |
| ABUW_1876 | 1.041 | 0.00150617 | *catC* | Muconolactone delta-isomerase |
| ABUW_2455 | 1.040 | 0.00106148 | - | Methylcrotonoyl-coa carboxylase subunit alpha |
| ABUW_0379 | 1.039 | 2.58E-37 | *dxs* | 1-deoxy-D-xylulose-5-phosphate synthase |
| ABUW_2097 | 1.024 | 8.58E-07 | *atoA* | 3-oxoacid coa-transferase, subunit B |
| ABUW_1683 | 1.023 | 8.05E-09 | - | Thioesterase family protein |
| ABUW_3156 | 1.022 | 3.30E-06 | - | Hypothetical protein |
| ABUW_1851 | 1.020 | 0.00130667 | - | Aminoglycoside phosphotransferase |
| ABUW_2686 | 1.017 | 0.0034193 | - | Hypothetical protein |
| ABUW_2720 | 1.017 | 1.57E-05 | - | Hypothetical protein |
| ABUW_2476 | 1.013 | 3.29E-11 | *dcaS* | Transcriptional regulator, IclR family |
| ABUW_3177 | 1.013 | 5.82E-10 | *nuoA* | NADH dehydrogenase I chain A |
| ABUW_0372 | 1.011 | 0.00126741 | - | Phage shock protein C |
| ABUW_3405 | 1.010 | 1.03E-07 | *leuA* | 2-isopropylmalate synthase |
| ABUW_1657 | 1.009 | 0.01109953 | - | Hypothetical protein |
| ABUW_2450 | 1.008 | 1.17E-05 | - | Long-chain fatty-acid-coa ligase |
| ABUW_0443 | 1.003 | 0.00556506 | *prlC* | Oligopeptidase A |
| ABUW_2218 | 1.003 | 2.08E-19 | *cysQ* | 3'(2'),5'-bisphosphate nucleotidase |
| ABUW_2091 | 1.002 | 7.72E-05 | - | Transcriptional regulator, AraC family |
| ABUW_1360 | 1.002 | 6.35E-08 | - | Hypothetical protein |

***Table S3b. RNAseq AB5075^-cavA-hfq^ down-regulated genes relative to AB5075^-cavA^.*** *Strains were cultured to OD_600nm_=0.5 in ID-CAMHB at 37ºC.*

| **Locus_tag** | **Log_2_FC** | **Pvalue** | **Gene** | **Product** |
| --- | --- | --- | --- | --- |
| ABUW_2784 | -1.004 | 0.01839215 | *-* | Hypothetical protein |
| ABUW_2921 | -1.005 | 0.02214311 | *-* | Hypothetical protein |
| ABUW_1387 | -1.010 | 0.0322071 | *-* | Hypothetical protein |
| ABUW_1246 | -1.011 | 5.95E-09 | *queF* | NADPH-dependent 7-cyano-7-deazaguanine reductase (EsvE1) |
| ABUW_0349 | -1.011 | 0.04984413 | *-* | Hypothetical protein |
| ABUW_3392 | -1.014 | 1.94E-13 | *gntT* | High affinity gluconate permease |
| ABUW_0245 | -1.014 | 2.84E-06 | *-* | Transcriptional regulator, XRE family |
| ABUW_0774 | -1.014 | 0.01944066 | *-* | Hypothetical protein |
| ABUW_1275 | -1.014 | 0.00316024 | *-* | Hypothetical protein |
| ABUW_0985 | -1.016 | 0.00427797 | *-* | Hypothetical protein |
| ABUW_1587 | -1.016 | 3.45E-07 | *-* | Hypothetical protein |
| ABUW_3710 | -1.017 | 0.00076219 | *-* | Putative acetyltransferase |
| ABUW_3439 | -1.017 | 5.48E-05 | *-* | Hypothetical protein |
| ABUW_0117 | -1.018 | 0.04362043 | *-* | Hypothetical protein |
| ABUW_3833 | -1.022 | 1.21E-09 | *ptk* | Tyrosine-protein kinase Ptk |
| ABUW_3312 | -1.023 | 0.01706421 | *pntB* | NAD(P) transhydrogenase subunit beta |
| ABUW_2915 | -1.023 | 3.02E-06 | *-* | Hypothetical protein |
| ABUW_0718 | -1.025 | 2.09E-07 | *maf1* | Septum formation protein Maf |
| ABUW_2048 | -1.027 | 0.03790923 | *-* | Phage replication protein |
| ABUW_0917 | -1.028 | 0.00595979 | *groS* | Chaperonin GroS |
| ABUW_1986 | -1.029 | 0.03543628 | *-* | Replication initiation factor |
| ABUW_2625 | -1.029 | 0.02697436 | *-* | Hypothetical protein |
| ABUW_3739 | -1.031 | 0.00023721 | *atpI* | ATP synthase F0, I subunit |
| ABUW_0118 | -1.033 | 0.05188653 | *-* | Hypothetical protein |
| ABUW_1221 | -1.035 | 5.95E-07 | *-* | Sensory box protein |
| ABUW_0266 | -1.036 | 0.0348633 | *czcC* | Cation efflux system protein |
| ABUW_0771 | -1.038 | 0.03943949 | *-* | Hypothetical protein |
| ABUW_1975 | -1.039 | NA | *adeB* | Multidrug efflux protein AdeB |
| ABUW_2513 | -1.044 | 0.03782334 | *csp2* | Cold-shock DNA-binding domain protein |
| ABUW_1386 | -1.044 | 0.03082676 | *-* | Hypothetical protein |
| ABUW_2444 | -1.046 | 0.00204667 | *-* | Hypothetical protein |
| ABUW_2021 | -1.047 | 0.02725473 | *-* | Hypothetical protein |
| ABUW_1901 | -1.049 | 0.04591768 | *-* | Hypothetical protein |
| ABUW_2261 | -1.050 | 8.37E-07 | *-* | MotA/TolQ/ExbB proton channel |
| ABUW_2256 | -1.055 | 1.41E-12 | *holB* | DNA polymerase III delta prime subunit |
| ABUW_0347 | -1.057 | 1.96E-05 | *-* | Amino-acid permease |
| ABUW_2041 | -1.060 | 0.03768963 | *-* | Hypothetical protein |
| ABUW_1991 | -1.060 | 0.03818416 | *-* | Hypothetical protein |
| ABUW_0792 | -1.063 | 0.01114931 | *-* | Hypothetical protein |
| ABUW_3424 | -1.063 | 2.12E-07 | *-* | Biopolymer transport protein ExbD/TolR |
| ABUW_2446 | -1.064 | 5.14E-05 | *-* | Transcriptional regulator, LysR family |
| ABUW_1268 | -1.064 | 0.01240653 | *-* | Hypothetical protein |
| ABUW_3391 | -1.068 | 1.13E-16 | *gntK* | Shikimate kinase |
| ABUW_3299 | -1.070 | 0.0009188 | *-* | DNA uptake protein |
| ABUW_1985 | -1.071 | 0.02128739 | *-* | Hypothetical protein |
| ABUW_0556 | -1.073 | 0.03486883 | *-* | Phage zinc-binding transcriptional activator |
| ABUW_1257 | -1.074 | 0.01911983 | *-* | Hypothetical protein |
| ABUW_1138 | -1.076 | 0.00633488 | *-* | Hypothetical protein |
| ABUW_0317 | -1.077 | 0.01997334 | *pilY* | Pilus assembly protein tip-associated adhesin PilY1 |
| ABUW_0756 | -1.078 | 0.00244326 | *-* | Hypothetical protein |
| ABUW_3051 | -1.078 | 8.20E-11 | *-* | Hypothetical protein |
| ABUW_2028 | -1.081 | 0.02731202 | *-* | Hypothetical protein |
| ABUW_3684 | -1.082 | 8.17E-05 | *-* | Hypothetical protein |
| ABUW_3482 | -1.085 | 2.50E-05 | *-* | Transcriptional regulator, HTH-type |
| ABUW_3356 | -1.087 | 4.25E-16 | *ilvN* | Acetolactate synthase, small subunit |
| ABUW_1408 | -1.088 | 0.01354428 | *-* | Hypothetical protein |
| ABUW_1681 | -1.091 | 0.0046425 | *-* | Hypothetical protein |
| ABUW_3118 | -1.094 | 2.03E-05 | *bioC* | Biotin biosynthesis protein BioC |
| ABUW_0757 | -1.097 | 0.00254452 | *-* | Hypothetical protein |
| ABUW_1253 | -1.098 | 0.00019832 | *-* | Hypothetical protein |
| ABUW_0319 | -1.098 | 0.01591741 | *comF* | Pilin like competence factor |
| ABUW_0677 | -1.100 | 0.00150628 | *-* | Hypothetical protein |
| ABUW_3780 | -1.101 | 8.71E-12 | *-* | Acyl-CoA dehydrogenase |
| ABUW_0235 | -1.102 | 2.70E-08 | *-* | Transporter LysE family |
| ABUW_0260 | -1.105 | 0.0059989 | *acr1* | Acyl coenzyme A reductase |
| ABUW_2216 | -1.117 | 0.00020698 | *rpsT* | Ribosomal protein S20 |
| ABUW_1412 | -1.121 | 0.00071648 | *-* | Hypothetical protein |
| ABUW_2994 | -1.122 | 0.01889015 | *-* | Hypothetical protein |
| ABUW_0267 | -1.123 | 0.02070968 | *czcB* | Cation efflux system protein (EsvF1) |
| ABUW_0772 | -1.124 | 5.57E-05 | *-* | Hypothetical protein |
| ABUW_1139 | -1.129 | 4.85E-10 | *-* | Hypothetical protein |
| ABUW_0776 | -1.130 | 0.00328123 | *-* | Hypothetical protein |
| ABUW_3117 | -1.132 | 1.11E-07 | *bioD* | Dethiobiotin synthase |
| ABUW_2861 | -1.140 | 0.0004683 | *-* | hypothetical protein |
| ABUW_3354 | -1.141 | 7.74E-11 | *-* | ggdef domain/eal domain protein |
| ABUW_0782 | -1.149 | 1.12E-06 | *-* | Prophage LambdaCh01, coat protein |
| ABUW_1925 | -1.157 | 3.45E-07 | *-* | Senescence marker protein-30 |
| ABUW_0274 | -1.158 | 4.22E-11 | *-* | Hypothetical protein |
| ABUW_3866 | -1.159 | 1.54E-05 | *-* | Hypothetical protein |
| ABUW_0698 | -1.159 | 9.84E-09 | *-* | Flavodoxin/nitric oxide synthase |
| ABUW_3722 | -1.163 | NA | *-* | Peptidoglycan-binding LysM |
| ABUW_2205 | -1.166 | 5.06E-07 | *hscB* | Fe-S protein assembly co-chaperone HscB |
| ABUW_1685 | -1.167 | 9.48E-15 | *-* | Hypothetical protein |
| ABUW_1233 | -1.170 | 3.23E-06 | *-* | OmpW family protein |
| ABUW_1523 | -1.171 | 0.01895414 | *-* | Transporter, major facilitator family |
| ABUW_2829 | -1.177 | 0.02016078 | *-* | Hypothetical protein |
| ABUW_3551 | -1.180 | 0.00574959 | *pilD* | type IV pilus prepilin peptidase PilD |
| ABUW_2652 | -1.184 | 0.00943323 | *-* | Hypothetical protein |
| ABUW_0351 | -1.184 | 1.05E-09 | *-* | Bifunctional succinylornithine transaminase/acetylornithine transaminase |
| ABUW_2445 | -1.194 | 0.00011699 | *-* | Hypothetical protein |
| ABUW_0739 | -1.201 | 0.006966 | *-* | Hypothetical protein |
| ABUW_1402 | -1.202 | 0.01406406 | *-* | Hypothetical protein |
| ABUW_0762 | -1.206 | 0.01241957 | *-* | Hypothetical protein |
| ABUW_1395 | -1.207 | 0.00948099 | *-* | Hypothetical protein |
| ABUW_1250 | -1.209 | 0.01357958 | *-* | Hypothetical protein |
| ABUW_2038 | -1.210 | 0.01236678 | *-* | Phage replication protein |
| ABUW_0770 | -1.210 | 0.00041899 | *-* | Hypothetical protein |
| ABUW_0936 | -1.213 | 0.01698158 | *-* | Hypothetical protein |
| ABUW_0784 | -1.216 | 2.63E-05 | *-* | Hypothetical protein |
| ABUW_0578 | -1.220 | 0.00082993 | *-* | Phage tail protein |
| ABUW_2782 | -1.224 | 0.01350494 | *hcaE* | Porin |
| ABUW_3362 | -1.227 | 1.39E-27 | *-* | Efflux transporter, RND family, MFP subunit |
| ABUW_2556 | -1.234 | 0.00918437 | *-* | Hypothetical protein |
| ABUW_1211 | -1.235 | 5.70E-07 | *-* | Hypothetical protein |
| ABUW_3779 | -1.235 | 5.75E-17 | *-* | Short-chain enoyl-CoA hydratase |
| ABUW_2353 | -1.237 | 0.00016024 | *-* | Lysine exporter protein |
| ABUW_2032 | -1.242 | 0.00418681 | *-* | Hypothetical protein |
| ABUW_3658 | -1.246 | 3.47E-05 | *-* | Hypothetical protein |
| ABUW_3632 | -1.247 | 3.55E-09 | *feoB* | Ferrous iron transport protein B |
| ABUW_1976 | -1.249 | 0.00437968 | *adeC* | Multidrug efflux protein AdeC |
| ABUW_2034 | -1.252 | 0.01276844 | *-* | Hypothetical protein |
| ABUW_3550 | -1.252 | 0.01329717 | *pilC* | Pilin biogenesis protein |
| ABUW_2044 | -1.253 | 0.01293475 | *-* | Hypothetical protein |
| ABUW_3425 | -1.256 | 7.19E-09 | *-* | MotA/TolQ/ExbB proton channel |
| ABUW_3719 | -1.259 | 9.11E-07 | *-* | TonB-dependent copper receptor |
| ABUW_1440 | -1.260 | 0.00441054 | *-* | Hypothetical protein |
| ABUW_3723 | -1.268 | 0.00458027 | *dprA* | DNA protecting protein DprA |
| ABUW_0313 | -1.268 | 0.00913966 | *fimT* | Pilin protein FimT |
| ABUW_0290 | -1.269 | 0.00876638 | *-* | Type IV pilus assembly protein PilM |
| ABUW_0348 | -1.270 | 0.00640489 | *-* | Hypothetical protein |
| ABUW_3314 | -1.275 | 0.00578677 | *pntA1* | NAD(P) transhydrogenase subunit alpha |
| ABUW_1276 | -1.277 | 0.0003 | *-* | Hypothetical protein |
| ABUW_1393 | -1.278 | 0.00163846 | *-* | Hypothetical protein |
| ABUW_1259 | -1.283 | 0.00304103 | *-* | Hypothetical protein |
| ABUW_3031 | -1.283 | 0.01218079 | *pilT* | Twitching mobility protein |
| ABUW_0751 | -1.283 | 0.00615542 | *-* | Hypothetical protein |
| ABUW_1324 | -1.285 | 0.00413342 | *-* | Hypothetical protein |
| ABUW_0777 | -1.286 | 7.42E-05 | *-* | Phage head morphogenesis protein |
| ABUW_1270 | -1.287 | 0.00512874 | *-* | Hypothetical protein |
| ABUW_0759 | -1.288 | 1.85E-05 | *-* | Hypothetical protein |
| ABUW_2521 | -1.288 | 0.00031225 | *-* | UvrD/REP helicase |
| ABUW_2047 | -1.289 | 0.00862687 | *-* | Hypothetical protein |
| ABUW_1388 | -1.299 | 0.0053978 | *-* | Hypothetical protein |
| ABUW_3549 | -1.304 | 0.01275652 | *pilB* | Type IV-A pilus assembly ATPase PilB |
| ABUW_0749 | -1.308 | 0.00703533 | *-* | Hypothetical protein |
| ABUW_0479 | -1.308 | 0.00019003 | *-* | Hypothetical protein |
| ABUW_1767 | -1.310 | 2.03E-08 | *-* | Hypothetical protein |
| ABUW_2962 | -1.321 | 9.96E-09 | *-* | YaeQ protein |
| ABUW_1390 | -1.324 | 0.00200821 | *-* | Hypothetical protein |
| ABUW_1641 | -1.330 | 0.00420321 | *-* | Hypothetical protein |
| ABUW_2938 | -1.336 | 4.56E-05 | *-* | Hypothetical protein |
| ABUW_3718 | -1.340 | 2.16E-07 | *-* | Hypothetical protein |
| ABUW_0353 | -1.347 | 5.10E-08 | *astD* | Succinylglutamic semialdehyde dehydrogenase |
| ABUW_0678 | -1.351 | NA | *pilG* | Type IV pilus response regulator receiver protein PilG |
| ABUW_0355 | -1.356 | 3.90E-08 | *astE* | Succinylglutamate desuccinylase |
| ABUW_1645 | -1.357 | NA | *-* | Transcriptional regulator, TetR family |
| ABUW_2185 | -1.363 | 2.40E-12 | *-* | Siderophore biosynthesis protein |
| ABUW_2321 | -1.373 | 6.83E-06 | *-* | Hypothetical protein |
| ABUW_0314 | -1.382 | 0.00710381 | *pilV* | type IV pilus modification protein PilV |
| ABUW_2051 | -1.382 | 0.00633476 | *-* | hypothetical protein |
| ABUW_0802 | -1.385 | 0.00081686 | *-* | hypothetical protein |
| ABUW_0807 | -1.385 | 3.01E-09 | *-* | hypothetical protein |
| ABUW_2975 | -1.391 | 2.98E-07 | *-* | homocysteine S-methyltransferase family protein |
| ABUW_1781 | -1.393 | 3.32E-26 | *-* | glutaredoxin-related protein |
| ABUW_0761 | -1.396 | 0.00308733 | *-* | hypothetical protein |
| ABUW_2976 | -1.399 | 1.23E-09 | *arcD* | arginine/ornithine antiporter |
| ABUW_1212 | -1.403 | 8.99E-07 | *-* | hypothetical protein |
| ABUW_2037 | -1.403 | 0.00399926 | *-* | hypothetical protein |
| ABUW_1401 | -1.404 | 0.0013534 | *-* | hypothetical protein |
| ABUW_1722 | -1.409 | 6.80E-07 | *fumC* | fumarate hydratase, class II |
| ABUW_2781 | -1.411 | 0.00456194 | *hcaD* | long-chain specific acyl-coa dehydrogenase |
| ABUW_2201 | -1.411 | 1.00E-07 | *iscR* | iscRSUA operon repressor |
| ABUW_0783 | -1.415 | 1.33E-05 | *-* | hypothetical protein |
| ABUW_0679 | -1.424 | 0.00542005 | *pilH* | type IV pilus response regulator protein PilH |
| ABUW_2517 | -1.434 | 0.00470923 | *-* | hypothetical protein |
| ABUW_3503 | -1.438 | 6.11E-09 | *-* | hypothetical protein |
| ABUW_2015 | -1.441 | 0.00064262 | *-* | hypothetical protein |
| ABUW_2828 | -1.447 | 8.30E-09 | *-* | TonB-dependent siderophore receptor |
| ABUW_0791 | -1.454 | 2.30E-05 | *-* | hypothetical protein |
| ABUW_0683 | -1.460 | 0.00321563 | *-* | hypothetical protein |
| ABUW_3032 | -1.460 | 0.00350306 | *pilU* | twitching motility protein |
| ABUW_1406 | -1.461 | 5.56E-07 | *-* | hypothetical protein |
| ABUW_0291 | -1.461 | 0.00375788 | *comN* | type 4 fimbrial biogenesis protein PilN |
| ABUW_0597 | -1.464 | 1.25E-13 | *-* | hypothetical protein |
| ABUW_2980 | -1.473 | 0.00092892 | *-* | hypothetical protein |
| ABUW_1698 | -1.474 | 3.13E-10 | *aroQ* | 3-dehydroquinate dehydratase, type II |
| ABUW_1394 | -1.475 | 0.0001128 | *-* | hypothetical protein |
| ABUW_1309 | -1.489 | 1.39E-05 | *-* | hypothetical protein |
| ABUW_1354 | -1.493 | 2.16E-13 | *-* | two-component system sensor protein |
| ABUW_3563 | -1.502 | 0.00136038 | *-* | transcriptional regulator, IclR family |
| ABUW_1247 | -1.503 | 2.50E-09 | *-* | ABC transporter permease protein |
| ABUW_1248 | -1.513 | 4.42E-08 | *-* | ABC transporter ATP-binding protein |
| ABUW_0680 | -1.527 | 0.00277398 | *pilI* | type IV pilus signal transduction protein PilI |
| ABUW_0116 | -1.529 | 3.62E-16 | *-* | hypothetical protein |
| ABUW_0354 | -1.537 | 1.57E-09 | *astB* | succinylarginine dihydrolase |
| ABUW_2860 | -1.542 | 1.41E-05 | *-* | hypothetical protein |
| ABUW_1974 | -1.548 | 0.00035988 | *adeA* | multidrug efflux protein AdeA |
| ABUW_0766 | -1.548 | 4.15E-05 | *-* | hypothetical protein |
| ABUW_2978 | -1.553 | 3.85E-09 | *-* | hypothetical protein |
| ABUW_3452 | -1.554 | 1.16E-05 | *-* | hypothetical protein |
| ABUW_0315 | -1.558 | 0.0022391 | *pilW* | pilus assembly protein PilW |
| ABUW_2004 | -1.562 | 0.00010869 | *-* | hypothetical protein |
| ABUW_0294 | -1.568 | 0.00077323 | *comQ* | fimbrial assembly protein PilQ |
| ABUW_0120 | -1.570 | 0.00046457 | *-* | hypothetical protein |
| ABUW_1999 | -1.574 | 0.00029342 | *-* | hypothetical protein |
| ABUW_1258 | -1.576 | 0.00016477 | *-* | hypothetical protein |
| ABUW_1273 | -1.580 | 3.65E-05 | *-* | hypothetical protein |
| ABUW_1411 | -1.584 | 9.19E-06 | *-* | hypothetical protein |
| ABUW_0159 | -1.596 | 1.32E-09 | *-* | TonB-dependent receptor protein |
| ABUW_3453 | -1.597 | 0.00039571 | *-* | hypothetical protein |
| ABUW_1307 | -1.600 | 1.71E-08 | *-* | hypothetical protein |
| ABUW_2276 | -1.623 | 9.07E-15 | *-* | transcriptional regulator, ArsR family |
| ABUW_0268 | -1.628 | 1.53E-08 | *czcA* | cation efflux system protein |
| ABUW_3641 | -1.628 | 0.00031647 | *pilR* | type 4 fimbriae expression regulatory protein PilR |
| ABUW_2042 | -1.634 | 0.00099436 | *-* | hypothetical protein |
| ABUW_3633 | -1.634 | 8.78E-16 | *feoA* | iron transporter |
| ABUW_0222 | -1.643 | 1.18E-06 | *-* | transcriptional regulator, TetR family |
| ABUW_0755 | -1.645 | 0.00032655 | *-* | hypothetical protein |
| ABUW_2186 | -1.646 | 5.00E-19 | *-* | siderophore biosynthesis protein |
| ABUW_0737 | -1.653 | 1.56E-13 | *-* | hypothetical protein |
| ABUW_2184 | -1.654 | 9.24E-18 | *-* | hypothetical protein |
| ABUW_0760 | -1.669 | 1.74E-13 | *-* | hypothetical protein |
| ABUW_1177 | -1.669 | 0.00015591 | *bauA* | ferric acinetobactin receptor |
| ABUW_1355 | -1.672 | 1.31E-08 | *-* | complement control module protein |
| ABUW_2985 | -1.677 | 0.0007242 | *-* | hypothetical protein |
| ABUW_0352 | -1.683 | 1.10E-11 | *astA1* | arginine N-succinyltransferase |
| ABUW_0681 | -1.685 | 0.00074872 | *pilJ* | type IV pilus methyl-accepting chemotaxis sensory transducer PilJ |
| ABUW_2035 | -1.686 | 0.00064148 | *-* | hypothetical protein |
| ABUW_3426 | -1.687 | 2.37E-17 | *-* | TonB protein |
| ABUW_0742 | -1.698 | 0.00011747 | *-* | hypothetical protein |
| ABUW_4090 | -1.706 | 0.00290779 | *-* | hypothetical protein |
| ABUW_0744 | -1.708 | 0.0001668 | *-* | hypothetical protein |
| ABUW_2981 | -1.716 | 0.0001802 | *-* | heme oxygenase |
| ABUW_2168 | -1.718 | 3.27E-07 | *-* | hypothetical protein |
| ABUW_2180 | -1.719 | 5.15E-19 | *-* | hypothetical protein |
| ABUW_0682 | -1.719 | 0.00044852 | *pilL* | type IV pilus hybrid sensor kinase/response regulator PilL |
| ABUW_1308 | -1.725 | 2.28E-06 | *-* | putative tail fiber |
| ABUW_3720 | -1.729 | 1.34E-31 | *-* | hypothetical protein |
| ABUW_0293 | -1.729 | 0.00047615 | *comL* | pilus assembly protein, PilQ |
| ABUW_2182 | -1.732 | 1.07E-15 | *-* | TonB-dependent receptor |
| ABUW_0763 | -1.751 | 4.53E-05 | *-* | hypothetical protein |
| ABUW_2918 | -1.775 | 1.88E-12 | *-* | rhomboid family protein |
| ABUW_0809 | -1.776 | 2.64E-05 | *-* | hypothetical protein |
| ABUW_1260 | -1.778 | 4.56E-05 | *-* | hypothetical protein |
| ABUW_0363 | -1.787 | 1.99E-07 | *-* | YCII-related protein |
| ABUW_2983 | -1.788 | 5.15E-05 | *-* | hypothetical protein |
| ABUW_1363 | -1.809 | 4.05E-22 | *-* | acyl-CoA dehydrogenase |
| ABUW_2045 | -1.815 | 0.00022062 | *-* | hypothetical protein |
| ABUW_0779 | -1.815 | 5.90E-07 | *-* | hypothetical protein |
| ABUW_0307 | -1.822 | 2.17E-08 | *-* | hypothetical protein |
| ABUW_0741 | -1.822 | 4.49E-05 | *-* | hypothetical protein |
| ABUW_1397 | -1.837 | 0.00022471 | *-* | hypothetical protein |
| ABUW_0735 | -1.847 | 5.73E-05 | *-* | hypothetical protein |
| ABUW_0753 | -1.847 | 3.06E-06 | *-* | hypothetical protein |
| ABUW_2167 | -1.854 | 1.49E-06 | *-* | iron-regulated membrane protein |
| ABUW_2166 | -1.872 | 2.58E-05 | *-* | hypothetical protein |
| ABUW_0292 | -1.876 | 0.00014371 | *comO* | pilus assembly protein, PilO |
| ABUW_2207 | -1.897 | 9.81E-16 | *fdx* | ferredoxin, 2Fe-2S type |
| ABUW_0780 | -1.951 | 4.33E-13 | *-* | hypothetical protein |
| ABUW_0750 | -1.954 | 1.28E-05 | *-* | hypothetical protein |
| ABUW_0758 | -1.964 | 2.39E-08 | *-* | hypothetical protein |
| ABUW_0752 | -1.968 | 9.28E-06 | *-* | replicative DNA helicase |
| ABUW_1272 | -2.005 | 7.62E-07 | *-* | hypothetical protein |
| ABUW_0734 | -2.040 | 4.07E-17 | *-* | hypothetical protein |
| ABUW_1396 | -2.045 | 4.41E-10 | *-* | hypothetical protein |
| ABUW_2010 | -2.048 | 1.81E-06 | *-* | hypothetical protein |
| ABUW_1176 | -2.048 | 4.18E-18 | *bauB* | ferric acinetobactin transport system periplasmic binding protein |
| ABUW_2181 | -2.053 | 1.75E-19 | *-* | hypothetical protein |
| ABUW_1269 | -2.053 | 2.07E-06 | *-* | hypothetical protein |
| ABUW_1439 | -2.061 | 6.77E-16 | *-* | hypothetical protein |
| ABUW_3465 | -2.074 | 3.86E-24 | *-* | transcriptional regulator, LysR family |
| ABUW_2982 | -2.084 | NA | *-* | TonB family protein |
| ABUW_2039 | -2.085 | 1.20E-08 | *-* | hypothetical protein |
| ABUW_2179 | -2.103 | 1.19E-24 | *-* | hypothetical protein |
| ABUW_0304 | -2.114 | 1.84E-05 | *-* | type IV pilin structural subunit |
| ABUW_0754 | -2.116 | 4.20E-07 | *-* | hypothetical protein |
| ABUW_2165 | -2.151 | 3.47E-09 | *-* | outer-membrane receptor for Fe(III)-coprogen,Fe(III)-ferrioxamine B and Fe(III)-rhodotrulic acid |
| ABUW_0803 | -2.161 | 2.02E-07 | *-* | hypothetical protein |
| ABUW_0781 | -2.174 | 6.71E-19 | *-* | hypothetical protein |
| ABUW_1325 | -2.187 | 2.31E-08 | *-* | hypothetical protein |
| ABUW_1364 | -2.199 | 4.05E-15 | *-* | alpha/beta hydrolase fold protein |
| ABUW_2183 | -2.223 | 2.23E-62 | *-* | dimethylmenaquinone methyltransferase |
| ABUW_2049 | -2.271 | 7.57E-08 | *-* | hypothetical protein |
| ABUW_0790 | -2.279 | 5.92E-11 | *-* | putative tail fiber |
| ABUW_2979 | -2.283 | 1.13E-09 | *-* | hypothetical protein |
| ABUW_0768 | -2.294 | 6.54E-08 | *-* | hypothetical protein |
| ABUW_1255 | -2.312 | 4.72E-10 | *-* | hypothetical protein |
| ABUW_2188 | -2.325 | 5.30E-15 | *-* | hypothetical protein |
| ABUW_0743 | -2.333 | 1.47E-07 | *-* | hypothetical protein |
| ABUW_2187 | -2.345 | 1.01E-19 | *-* | transporter, major facilitator family |
| ABUW_0767 | -2.354 | 6.88E-08 | *-* | hypothetical protein |
| ABUW_0789 | -2.370 | 4.34E-15 | *-* | hypothetical protein |
| ABUW_1598 | -2.409 | 3.59E-13 | *-* | hypothetical protein |
| ABUW_1166 | -2.434 | 7.96E-42 | *-* | threonine ammonia-lyase |
| ABUW_1655 | -2.479 | 1.98E-08 | *-* | TonB-dependent siderophore receptor |
| ABUW_1174 | -2.510 | 2.21E-10 | *bauC* | ferric acinetobactin transport system permease |
| ABUW_1173 | -2.571 | 5.66E-10 | *bauD* | ferric acinetobactin transport system permease |
| ABUW_1175 | -2.587 | 1.94E-14 | *bauE* | ferric acinetobactin transport system ATP-binding protein |
| ABUW_0748 | -2.658 | 9.31E-10 | *-* | hypothetical protein |
| ABUW_2189 | -2.695 | 1.85E-19 | *-* | IucA/IucC-family aerobactin siderophore biosynthesis component |
| ABUW_2074 | -2.736 | 3.57E-27 | *-* | transcriptional regulator, fur family |
| ABUW_1170 | -2.767 | 3.15E-18 | *basB* | non-ribosomal peptide synthetase |
| ABUW_2916 | -2.806 | 2.71E-16 | *pfeA* | TonB dependent outer membrane siderophore receptor protein |
| ABUW_1167 | -2.817 | 5.64E-24 | *-* | hypothetical protein |
| ABUW_0740 | -2.886 | 3.35E-11 | *-* | hypothetical protein |
| ABUW_3403 | -2.901 | 3.27E-17 | *-* | TonB-dependent receptor |
| ABUW_1251 | -2.901 | 1.57E-07 | *-* | hypothetical protein |
| ABUW_1178 | -2.914 | 4.57E-34 | *basC* | nonribosomal peptide synthetase BasC |
| ABUW_1168 | -2.952 | 1.29E-16 | *bauF* | siderophore-interacting protein |
| ABUW_2986 | -2.966 | 1.65E-16 | *-* | putative transmembrane sensor protein FecR |
| ABUW_1172 | -2.974 | 8.68E-13 | *-* | hypothetical protein |
| ABUW_0143 | -3.021 | 5.60E-20 | *-* | TonB-dependent receptor protein |
| ABUW_2987 | -3.052 | 1.99E-19 | *-* | RNA polymerase sigma factor FecI |
| ABUW_2075 | -3.056 | 1.34E-31 | *-* | isochorismatase |
| ABUW_1183 | -3.109 | 2.55E-23 | *-* | hypothetical protein |
| ABUW_1185 | -3.176 | 1.47E-39 | *-* | ABC transporter |
| ABUW_3464 | -3.215 | 1.66E-29 | *-* | hypothetical protein |
| ABUW_3564 | -3.297 | 2.21E-13 | *alkB* | DNA repair system |
| ABUW_1184 | -3.428 | 4.69E-37 | *-* | ABC transporter, ATP-binding protein |
| ABUW_1169 | -3.454 | 1.20E-33 | *basA* | non-ribosomal peptide synthetase |
| ABUW_1186 | -3.461 | 4.36E-53 | *basH* | thioesterase |
| ABUW_1182 | -3.553 | 1.44E-31 | *basG* | histidine decarboxylase |
| ABUW_2076 | -3.640 | 7.75E-46 | *-* | 2,3-dihydro-2,3-dihydroxybenzoate dehydrogenase |
| ABUW_1179 | -3.685 | 1.93E-35 | *basD* | nonribosomal peptide synthetase BasD |
| ABUW_1171 | -3.754 | 3.41E-21 | *-* | hypothetical protein |
| ABUW_1188 | -3.799 | 3.51E-55 | *basJ* | isochorismate synthetase |
| ABUW_1181 | -3.879 | 2.58E-34 | *basF* | 2,3 dihydro-2,3 dihydroxybenzoate synthase |
| ABUW_2178 | -3.905 | 5.99E-34 | *-* | rhizobactin siderophore biosynthesis protein RhbD |
| ABUW_1187 | -3.932 | 2.09E-54 | *basI* | phosphopantetheinyl transferase |
| ABUW_3565 | -4.066 | 7.38E-21 | *ada* | methylated-DNA-protein-cysteine methyltransferase transcription regulator |
| ABUW_1180 | -4.174 | 1.08E-27 | *basE* | 2,3-dihydroxybenzoate-AMP ligase |

***Table S4a. Proteomic analysis of AB5075^-cavA^ expressed proteins with increased relative abundance to AB5075 WT.*** *Strains were cultured to OD_600nm_=0.5 in ID-CAMHB at 37ºC.*

| **UniProt ID** | **Protein Descriptions** | **Log_2_ FC** | **Pvalue** | **Genes** |
| --- | --- | --- | --- | --- |
| U5QBM2 | Signal peptidase I | 6.133 | 0.00235785 | lepB |
| A0A0D5YIQ5 | Fimbria/pilus outer membrane usher protein | 4.808 | 5.90E-06 | ABUW_2054 |
| A0A0D5YIL4 | Fimbrial protein | 4.701 | 5.08E-06 | ABUW_2055 |
| V5VD49 | Fimbria/pilus periplasmic chaperone | 4.659 | 2.49E-08 | fimC_2 |
| V5VCZ7 | Uncharacterized protein | 4.592 | 0.00022724 | A7M90_12425 |
| A0A0D5YHJ2 | DUF4882 domain-containing protein | 3.911 | 3.34E-09 | ABUW_2056 |
| V5VCT5 | Ferrous iron transporter B | 3.904 | 3.83E-07 | hifA |
| A0A0D5YHR1 | Poly-beta-1,6-N-acetyl-D-glucosamine N-deacetylase PgaB | 3.851 | 1.16E-06 | pgaB |
| A0A0D5YG83 | Poly-beta-1,6-N-acetyl-D-glucosamine synthase | 3.612 | 0.06651501 | pgaC |
| A0A0D5YG80 | Outer membrane protein | 3.439 | 7.10E-11 | pgaA |
| A0A059ZTI4 | Secreted protein | 3.426 | 3.44E-07 | A7M90_15590 |
| A0A0D5YE22 | Putative membrane protein | 2.961 | 8.94E-10 | ABR2091_3300 |
| X2FEH6 | Bacterial conjugation TrbI-like protein | 2.810 | 4.73E-07 | traB |
| A0A0D5YNZ3 | Matrixin superfamily | 2.777 | 4.14E-08 | ABUW_3881 |
| U5QBD8 | TraE | 2.634 | 7.21E-05 | traE |
| U5QEZ8 | TraK | 2.605 | 5.85E-07 | traK |
| U5QES9 | Membrane lipoprotein lipid attachment site | 2.588 | 0.00153715 | traV |
| V5VB91 | Csu fimbrial major subunit CsuAB | 2.568 | 0.0003211 | csuAB |
| U5QEU2 | Lipoprotein | 2.243 | 0.05043482 | ABUW_4035 |
| A0A0D5YF17 | Rubredoxin-NAD(+) reductase | 2.197 | 0.06059514 | ABUW_1151 |
| A0A0B3YZD4 | Thiol:disulfide interchange protein | 2.049 | 5.40E-07 | dsbC_3 |
| V5VDM1 | Fimbrial protein | 2.043 | 0.0053496 | ABUW_2313 |
| U5QBB6 | Conserved protein involved in tellurite resistance | 2.038 | 0.01163685 | ABUW_4028 |
| V5VFG4 | 3-oxoacyl-[acyl-carrier-protein] reductase FabG | 1.906 | 0.13613227 | fabG_5 |
| A0A081H3B8 | Chloride channel protein | 1.885 | 0.00036009 | clcA |
| A0A0B4B5C5 | Uncharacterized protein | 1.814 | 0.0004303 | A7M90_15725 |
| V5V9K5 | Anti-sigma factor antagonist | 1.813 | 0.0710558 | mlaB |
| U5QBP3 | Uncharacterized protein | 1.741 | 9.20E-06 | A7M90_15640 |
| U5QBF7 | Uncharacterized protein | 1.641 | 0.0007869 | ABUW_4029 |
| U5QEP9 | AAA-like domain protein | 1.633 | 2.53E-05 | traC |
| U5QBA4 | TraN | 1.596 | 6.14E-06 | traN |
| A0A059ZHN2 | Uncharacterized protein | 1.575 | 0.00183847 | ABUW_0356 |
| U5QBN0 | Uncharacterized protein | 1.574 | 0.00193202 | ABUW_4031 |
| A0A0D5YH83 | LysR family transcriptional regulator | 1.559 | 0.07328427 | ABUW_1936 |
| A0A0D5YG16 | CsuC | 1.447 | 5.51E-06 | ABUW_1490 |
| A0A0D5YG17 | CsuE | 1.422 | 0.00255934 | csuE |
| A0A0D5YHJ9 | CsuD | 1.376 | 0.05269778 | csuD |
| A0A0D5YKU0 | Rhs element Vgr protein, putative | 1.376 | 1.75E-07 | ABUW_2817 |
| A0A0J8TTU0 | Transmembrane protein | 1.372 | 2.63E-07 | ABUW_0243 |
| U5QEW5 | Uncharacterized protein | 1.369 | 6.57E-06 | A7M90_15580 |
| A0A0D5YJN1 | Aromatic amino acid transport protein AroP | 1.357 | 0.04875376 | ABUW_2808 |
| A0A0D5YMF9 | 4'-phosphopantetheinyl transferase | 1.356 | 0.0040247 | ABUW_3372 |
| A0A0D5YKY5 | YD repeat protein | 1.347 | 2.16E-06 | ABUW_2816 |
| A0A0D5YM95 | SDR family NAD(P)-dependent oxidoreductase | 1.322 | 3.32E-05 | ABUW_3798 |
| A0A0Q1IMM6 | Alpha-ketoglutarate permease | 1.314 | 0.01695151 | pcaT |
| X2FAQ7 | Ribbon-helix-helix protein CopG domain-containing protein | 1.296 | 1.31E-05 | ABUW_4074 |
| A0A090C0Q9 | DUF4184 domain-containing protein | 1.287 | 0.00220317 | ABUW_0599 |
| A0A0D5YD47 | Intermembrane phospholipid transport system permease protein MlaE | 1.253 | 0.00011767 | mlaE |
| V5VBN5 | Alpha/beta fold hydrolase | 1.239 | 0.01378453 | A7M90_04225 |
| X2FIS0 | TraH | 1.230 | 0.00476793 | traH |
| A0A059ZNI7 | RNA polymerase factor sigma-70 | 1.217 | 0.02695772 | sigX |
| A0A0D5YF70 | Dipeptide/tripeptide permease | 1.171 | 0.00190531 | ABUW_1206 |
| V5V921 | ABC transporter periplasmic substrate-binding protein | 1.106 | 6.59E-07 | mlaD |
| A0A0D5YHD2 | Uncharacterized protein | 1.102 | 0.00509742 | ABUW_1990 |
| V5V9K7 | Large-conductance mechanosensitive channel | 1.075 | 2.36E-07 | mscL |
| A0A0D5YEC9 | Uncharacterized protein | 1.073 | 0.00467805 | ABUW_0902 |
| A0A0D5YDX9 | Toluene tolerance efflux transporter | 1.073 | 3.65E-05 | mlaF |
| V5VAI1 | Acidic transcription factor A | 1.063 | 0.00096776 | A7M90_18910 |
| V5VBV2 | Type II toxin-antitoxin system HicA family toxin | 1.052 | 0.0019303 | A7M90_03990 |
| A0A0B4AU91 | Uncharacterized protein | 1.047 | 0.02588043 | A7M90_15760 |
| A0A0D5YNQ0 | Low temperature requirement protein A | 1.010 | 0.00027872 | ltrA |
| A0A0D5YLB1 | Cobalamin biosynthesis protein CobW | 1.008 | 4.42E-06 | ABUW_2964 |
| A0A0D5YDV8 | Peptidase S8 | 1.000 | 0.00990688 | ABUW_0358 |
| V5VAL7 | DUF5713 family protein | 0.996 | 0.11975235 | A7M90_19015 |
| V5VI75 | 2-amino-4-hydroxy-6-hydroxymethyldihydropteridine pyrophosphokinase | 0.989 | 0.10102498 | folK |
| U5QBJ7 | TraD | 0.982 | 6.39E-06 | trwB |
| A0A0D5YKU8 | Protein tolA | 0.981 | 0.00064039 | ABUW_2724 |
| A0A098SCQ1 | Signal peptide protein | 0.979 | 4.32E-06 | A7M90_16810 |
| C0JBD5 | Orf4 | 0.976 | 8.98E-08 | orf4 |
| A0A0D5YLF7 | Catalase-peroxidase | 0.957 | 1.58E-05 | katG |
| V5VAM6 | Diacylglycerol kinase | 0.950 | 0.00186712 | bmrU |
| V5V9K9 | ABC transporter substrate-binding protein | 0.949 | 3.24E-06 | mlaC |
| V5VB00 | DUF541 domain-containing protein | 0.946 | 4.18E-10 | ABUW_1108 |
| U5QBF1 | Conjugal transfer protein TraG | 0.934 | 5.68E-05 | traG |
| A0A0D5YJY0 | Lipoprotein | 0.932 | 1.02E-07 | A7M90_09100 |
| V5VHB1 | Cation:proton antiporter | 0.919 | 0.00150417 | mrpF |
| A0A077GKT8 | Anthranilate synthase | 0.916 | 7.44E-05 | A7M90_11190 |
| A0A0D5YI18 | 3-carboxy-cis,cis-muconate cycloisomerase | 0.910 | 0.07685523 | pcaB |
| A0A0D5YEW8 | Coproporphyrinogen III oxidase | 0.892 | 0.00016871 | ABUW_0684 |
| U5QBG2 | Toxin | 0.872 | 1.56E-05 | ABUW_4003 |
| A0A077G9D7 | Transglycosylase | 0.869 | 1.95E-05 | slt_1 |
| A0A090B4N0 | DUF2147 domain-containing protein | 0.855 | 2.64E-09 | A7M90_06480 |
| V5VGU2 | Ribosome hibernation promoting factor | 0.845 | 0.02321902 | raiA |
| U5QBD2 | Uncharacterized protein | 0.843 | 1.27E-05 | ABUW_4033 |
| V5VGM5 | Integral membrane protein | 0.829 | 0.03066171 | A7M90_00065 |
| A0A0D5YMA4 | Ohr family peroxiredoxin | 0.817 | 0.09804677 | ABUW_3265 |
| A0A0D5YF26 | DUF7944 domain-containing protein | 0.810 | 1.95E-07 | ABUW_1128 |
| U5QBB1 | DUF1173 domain-containing protein | 0.803 | 0.0088218 | A7M90_15735 |
| A0A0D5YDM0 | Sulfate transporter | 0.802 | 6.30E-05 | ABUW_0259 |
| A0A0D5YJ82 | HTH hxlR-type domain-containing protein | 0.795 | 0.00546754 | ABUW_2117 |
| A0A0D7T9H5 | MATE efflux family protein | 0.790 | 0.05236474 | A7M90_07395 |
| A0A0D5YK54 | Sel1 repeat family | 0.790 | 0.00011642 | ABUW_2601 |
| V5VGY0 | DNA gyrase inhibitor YacG | 0.782 | 0.0859372 | yacG |
| A0A0D5YPB6 | Uncharacterized protein | 0.776 | 0.00014191 | ABUW_4120 |
| A0A0D8V3K3 | HTH-type transcriptional repressor NemR | 0.776 | 4.55E-07 | nemR_3 |
| V5VA92 | EcnB | 0.776 | 8.25E-05 | ecnB |
| A0A0D5YNE6 | Zinc ribbon-containing protein | 0.774 | 0.0009393 | ABUW_3804 |
| V5VH02 | Chaperone modulator CbpM | 0.773 | 1.93E-05 | cbpM |
| A0A0D5YDP6 | Uncharacterized protein | 0.769 | 0.05596491 | ABUW_0236 |
| A0A0D5YGS6 | Peptidase S1 and S6 | 0.767 | 9.73E-05 | algW |
| A0A0D5YIN1 | L-sorbosone dehydrogenase | 0.761 | 2.76E-06 | sndH2 |
| A0A0D5YDY0 | Probable membrane transporter protein | 0.754 | 0.0335304 | yfcA |
| A0A0D5YLK9 | RNA polymerase sigma factor FecI | 0.751 | 0.02718243 | ABUW_2987 |
| A0A1A1SVI5 | Rubredoxin | 0.737 | 0.05097911 | rubA |
| A0A0D5YJG3 | 17 kDa surface antigen | 0.734 | 0.00384007 | ABUW_2678 |
| A0A0D5YH21 | Omega-amino acid--pyruvate aminotransferase | 0.729 | 0.05996386 | ABUW_1463 |
| A0A0D5YI69 | TetR family transcriptional regulator | 0.725 | 0.01990673 | ABUW_1895 |
| V5VDY0 | DUF1049 domain-containing protein | 0.718 | 2.63E-05 | A7M90_11725 |
| A0A0D5YHT3 | Potassium-transporting ATPase KdpC subunit | 0.718 | 2.22E-05 | kdpC |
| V5XWX9 | GrpB family protein | 0.714 | 9.99E-07 | ABUW_2288 |
| A0A059ZLN5 | 4-hydroxyphenylacetate 3-monooxygenase reductase component | 0.708 | 0.0177142 | rutF |
| A0A081GU02 | Carbapenem susceptibility porin CarO | 0.707 | 3.91E-08 | carO |
| A0A0D5YHX0 | Quinoprotein glucose dehydrogenase | 0.707 | 0.00010765 | gdhB2 |
| A0A0D5YII0 | MFS permease | 0.694 | 0.11037668 | ABUW_2403 |
| A0A0D5YNJ1 | TetR family transcriptional regulator | 0.683 | 0.01347395 | ABUW_3716 |
| A0A0D5YJP6 | RHS repeat-associated core domain-containing protein | 0.682 | 6.61E-05 | ABUW_2830 |
| V5VI42 | DoxX family membrane protein | 0.682 | 4.06E-05 | yqjF |
| A0A0D5YMH1 | HdeD family acid-resistance protein | 0.678 | 8.04E-06 | ABUW_3438 |
| A0A090B0M2 | Sel1 repeat family protein | 0.673 | 0.00088913 | A7M90_17020 |
| V5V9U8 | Acyl-CoA dehydrogenase | 0.673 | 0.00157599 | ABUW_0487 |
| V5VBY5 | Cation:proton antiporter | 0.670 | 2.08E-05 | nhaP |
| A0A0D5YCN3 | Helix-turn-helix domain-containing protein | 0.666 | 0.09876955 | ABUW_0245 |
| A0A090BC42 | Small-conductance mechanosensitive channel | 0.665 | 8.90E-05 | A7M90_13550 |
| A0A0D5YIN6 | Quinoprotein glucose dehydrogenase-B | 0.665 | 6.96E-07 | gdhB1 |
| A0A0D5YI08 | (2Fe-2S)-binding protein | 0.664 | 0.00547145 | ABUW_2184 |
| A0A0D5YCB0 | PGAP1 family protein | 0.663 | 0.00013531 | ABUW_0122 |
| Q2FD94 | Efflux pump membrane transporter | 0.662 | 4.72E-05 | adeJ |
| A0A081GZA3 | Pyrroloquinoline-quinone synthase | 0.662 | 5.27E-08 | pqqC |
| A0A0D5YMN7 | Curved DNA-binding protein | 0.661 | 1.92E-05 | cbpA |
| A0A081H921 | Amino acid efflux pump, RhtB family | 0.657 | 0.00025229 | A7M90_13625 |
| A0A0D5YDB3 | Esterase | 0.649 | 0.00608846 | ABUW_0123 |
| V5VJ35 | Lipoprotein, putative | 0.648 | 0.00064615 | ABR2091_0174 |
| V5V9V3 | ClpXP protease specificity-enhancing factor | 0.645 | 3.04E-05 | sspB |
| V5V8S0 | LemA | 0.636 | 6.96E-05 | A7M90_14370 |
| Q5I2N2 | Beta-lactamase OXA-51-like protein | 0.636 | 5.60E-07 | bla-OXA-69 |
| D6NSM8 | Beta-lactamase | 0.635 | 1.68E-05 | ampC |
| A0A059ZTE6 | Uncharacterized protein | 0.635 | 0.08393919 | A7M90_15810 |
| A0A0D5YL68 | RHS repeat protein | 0.632 | 3.59E-07 | ABUW_2831 |
| A0A0D5YJ98 | Lipoyl synthase | 0.629 | 0.01755241 | lipA2 |
| A0A0D5YD80 | Uncharacterized protein | 0.627 | 0.00012844 | ABUW_0490 |
| A0A0C0B9L4 | Thiol:disulfide interchange protein | 0.622 | 4.40E-06 | dsbC_2 |
| A0A0D5YL42 | Anti-anti-sigma regulatory factor (Antagonist of anti-sigma factor) | 0.620 | 0.16358757 | ABUW_3261 |
| A0A0D8GPY7 | Right-handed parallel beta-helix repeat-containing protein | 0.617 | 0.00077141 | ABUW_2660 |
| A0A0D5YP08 | Conjugal transfer protein TraF | 0.617 | 0.00325023 | traF |
| A0A0D8G6G4 | Major facilitator superfamily MFS_1 | 0.616 | 0.06977286 | nreB |
| A0A0D5YEF4 | SgcJ/EcaC family oxidoreductase | 0.615 | 0.00265882 | ABUW_0931 |
| V5VE35 | Uncharacterized protein | 0.615 | 0.0203505 | A7M90_05605 |
| V5V8V2 | Ion channel | 0.615 | 0.01026436 | A7M90_07030 |
| A0A0D5YFC4 | Patatin-like phospholipase family protein | 0.615 | 2.14E-05 | ABUW_0823 |
| A0A0D5YJZ5 | Signal peptide-containing protein | 0.613 | 0.01280852 | ABUW_2832 |
| A0A0D5YIN9 | LysR family transcriptional regulator | 0.602 | 0.00291889 | ABUW_2080 |
| A0A0D5YIU8 | Uncharacterized protein | 0.602 | 0.08302887 | ABUW_2094 |
| A0A0D5YIN2 | Oxidoreductase | 0.600 | 0.0005098 | ABUW_2458 |
| V5VHT9 | DUF445 domain-containing protein | 0.598 | 0.00255469 | A7M90_08395 |
| V5VGD7 | GlmZ(SRNA)-inactivating NTPase | 0.597 | 0.0003108 | rapZ |
| A0A0D5YFE3 | Sulfur carrier protein FdhD | 0.596 | 0.01038935 | fdhD |
| A0A0D5YIX7 | DUF937 domain-containing protein | 0.594 | 0.00010294 | ABUW_2161 |
| A0A0D8GPZ2 | Lipoprotein, putative | 0.592 | 0.00028405 | ABUW_2654 |
| A0A0D5YD57 | Uncharacterized protein | 0.591 | 0.00177264 | ABUW_0460 |
| A0A0D5YEU8 | GFA family protein | 0.591 | 1.28E-05 | ABUW_1042 |

***Table S4b. Proteomic analysis of AB5075^-cavA^ expressed proteins with decreased relative abundance to AB5075 WT.*** *Strains were cultured to OD_600nm_=0.5 in ID-CAMHB at 37ºC.*

| **UniProt ID** | **Protein Descriptions** | **Log_2_ FC** | **Pvalue** | **Genes** |
| --- | --- | --- | --- | --- |
| A0A0D5YCX7 | Prepilin-type N-terminal cleavage/methylation domain-containing protein | -9.013 | 1.48E-06 | ABUW_0304 |
| A0A059ZR03 | Pilus assembly protein | -6.046 | 0.01444621 | pilX |
| A0A0D5YFD1 | Chemotaxis protein CheA | -5.843 | 4.22E-06 | pilL |
| V5VA86 | Chemotaxis protein | -5.8 | 0.00186429 | pilJ |
| A0A0D5YLN2 | Type IV-A pilus assembly ATPase PilB | -5.371 | 1.01E-05 | pilB |
| A0A0D5YIK7 | KGG domain-containing protein | -5.293 | 0.00025863 | ABUW_2433 |
| A0A0D5YLB6 | Twitching mobility protein | -5.201 | 0.00118496 | pilT |
| A0A0D5YDX1 | Pilus assembly protein tip-associated adhesin PilY1 | -4.96 | 2.63E-05 | pilY |
| A0A0D5YND1 | Response regulator | -4.867 | 5.89E-05 | pilR |
| A0A0D5YDQ7 | CheW-like domain-containing protein | -4.826 | 9.72E-05 | ABUW_0683 |
| V5VA30 | Alkaline phosphatase synthesis transcriptional regulatory protein PhoP | -4.807 | 2.09E-05 | pilG |
| A0A0D5YK07 | Chemotaxis protein | -4.787 | 0.00014983 | ABUW_2842 |
| A0A0D5YMI4 | FimV domain-containing protein | -4.727 | 3.98E-06 | ABUW_3453 |
| A0A0D5YFC5 | Uncharacterized protein | -4.662 | 0.00181284 | ABUW_0677 |
| A0A0D5YCR0 | Type 4 fimbrial biogenesis protein PilN | -4.52 | 4.24E-07 | comN |
| A0A0D5YHE2 | Uncharacterized protein | -4.412 | 1.09E-06 | ABUW_1641 |
| A0A077GRE9 | Competence protein A | -4.081 | 2.52E-05 | A7M90_14315 |
| A0A0D5YK33 | Uncharacterized protein | -3.859 | 0.00023143 | ABUW_2434 |
| A0A0D5YDS3 | Pilin like competence factor | -3.779 | 0.00803335 | comE |
| A0A0Q2HWF1 | Hemerythrin | -3.773 | 0.01282174 | A7M90_16365 |
| A0A0D5YEX4 | Lipoprotein | -3.761 | 1.19E-05 | ABUW_0514 |
| V5V999 | Pilus assembly protein, PilO | -3.75 | 5.14E-06 | pilO |
| A0A086HYC5 | PilT/PilU family type 4a pilus ATPase | -3.742 | 0.00199977 | pilU |
| A0A0D5YCW8 | Fimbrial assembly protein PilQ | -3.639 | 9.14E-05 | comQ |
| A0A0D5YE91 | Energy transducer TonB | -3.539 | 8.97E-06 | ABR2091_3221 |
| A0A0D5YME7 | LysM peptidoglycan-binding domain-containing protein | -3.454 | 1.83E-05 | ABUW_3722 |
| A0A0D8GGD3 | Chemotaxis protein CheW | -3.336 | 9.57E-05 | pilI |
| V5VHX5 | GlsB/YeaQ/YmgE family stress response membrane protein | -3.222 | 0.01880432 | A7M90_19275 |
| A0A0D5YJA5 | Pilus assembly protein | -3.082 | 0.00055469 | pilZ |
| A0A0D5YH74 | Type II secretion system protein E | -3.062 | 0.04003033 | ABUW_1351 |
| A0A0D5YM11 | Prepilin leader peptidase/N-methyltransferase | -3.039 | 0.00464763 | pilD |
| V5V9M9 | PilH | -3.037 | 6.63E-05 | pilH |
| V5VAN6 | Signal peptide | -3.017 | 0.00313801 | A7M90_10385 |
| A0A0D5YK97 | Glutathione-dependent formaldehyde dehydrogenase | -2.953 | 2.00E-07 | ABUW_2594 |
| A0A0D5YDP9 | Pilus assembly protein PilP | -2.942 | 4.43E-05 | comL |
| E9L098 | StrB | -2.692 | 5.38E-05 | strB |
| V5VC75 | Fimbrial major subunit CsuA/B family protein | -2.69 | 1.23E-05 | A7M90_01585 |
| A0A0D5YCY9 | Pilin like competence factor | -2.642 | 2.35E-05 | comF |
| A0A0D5YNC1 | DNA-processing protein DprA | -2.631 | 0.00076724 | dprA |
| V5VEG3 | General stress protein 39 | -2.591 | 0.00034117 | ydaD |
| A0A0D5YIU7 | Surface antigen | -2.568 | 2.96E-05 | ABUW_2440 |
| A0A6H3SVU9 | 4-carboxymuconolactone decarboxylase | -2.289 | 0.01134774 | ABR2091_1362 |
| A0A0D5YEH0 | Single-stranded DNA-binding protein | -2.237 | 0.00043448 | ssb |
| A0A0D5YJ84 | MotA/TolQ/ExbB proton channel | -2.212 | 0.0001296 | ABUW_2261 |
| A0A0D5YCJ1 | Flavoprotein | -2.207 | 0.00013158 | ABR2091_3523 |
| A0A0D5YLX0 | DUF3592 domain-containing protein | -2.153 | 0.00337623 | ABUW_3191 |
| A0A0D5YJV8 | Monooxygenase | -2.044 | 0.00985054 | luxA |
| V5V8Y6 | GDT1 family protein | -2.043 | 0.16039878 | A7M90_07160 |
| A0A0D5YF34 | EAL domain-containing protein | -1.995 | 0.00015456 | ABUW_1138 |
| A0A0D5YLN9 | Putative cold shock protein | -1.921 | 0.00318641 | ABUW_3162 |
| V5VH04 | Pilin biogenesis protein | -1.864 | 0.00033556 | epsF_1 |
| A0A0D8G9M9 | PilW family protein | -1.861 | 0.01524085 | pilW |
| A0A0D5YNQ3 | Putative oxidoreductase | -1.838 | 0.03511904 | ABUW_3843 |
| A0A0D5YFZ2 | Acyl-CoA dehydrogenase | -1.82 | 1.94E-05 | ABUW_1467 |
| A0AA90HUQ3 | Carnitine monooxygenase oxygenase subunit | -1.793 | 9.28E-05 | cntA |
| A0A0D5YH26 | Carbohydrate porin | -1.792 | 0.0277246 | quiX |
| A0A0D5YDC7 | 3'-5' exonuclease | -1.788 | 0.01325361 | ABUW_0541 |
| A0A077GCA9 | Carnitine dehydratase | -1.751 | 0.00770389 | fldA |
| A0A0D5YF35 | Capsid protein | -1.734 | 0.00753699 | ABUW_0581 |
| A0A0D5YMB0 | DUF4124 domain-containing protein | -1.726 | 0.00074179 | ABUW_3358 |
| A0A0D5YJV9 | SPOR domain-containing protein | -1.69 | 0.0091241 | ABR2091_0979 |
| A0A0D5YMN6 | Uncharacterized protein | -1.688 | 8.74E-06 | ABUW_3452 |
| A0A0D5YIG5 | 4-hydroxybenzoate transporter | -1.671 | 0.01285717 | pcaK1 |
| A0A0D5YJP1 | ATP-binding cassette domain-containing protein | -1.67 | 0.00185271 | ABUW_2422 |
| A0A0D5YIJ7 | ABC transporter permease subunit | -1.644 | 0.00169623 | ABUW_2423 |
| A0A0D5YLK1 | Trehalose 6-phosphate phosphatase | -1.594 | 0.00117194 | otsB |
| A0A0D5YGE2 | Fimbria/pilus periplasmic chaperone | -1.589 | 5.51E-07 | ABUW_1632 |
| V5VBM8 | DUF2171 domain-containing protein | -1.587 | 0.00039618 | A7M90_04275 |
| A0A0D5YGT2 | Abasic site processing protein | -1.526 | 0.00125028 | ABUW_1761 |
| A0A0D5YLX1 | Uncharacterized protein | -1.502 | 0.00189069 | ABUW_3503 |
| A0A0D5YKI0 | DUF4142 domain-containing protein | -1.465 | 1.52E-07 | ABUW_2679 |
| A0A0D5YEF7 | Biofilm-associated protein | -1.452 | 4.84E-09 | ABUW_0885 |
| A0A0D5YGQ5 | Extracellular serine protease | -1.418 | 0.02033475 | ABUW_1759 |
| V5VA16 | Uncharacterized protein | -1.383 | 4.96E-07 | A7M90_18985 |
| V5VJF6 | RcnB family protein | -1.382 | 0.00026369 | ABUW_3851 |
| V5VEP6 | Lipoprotein | -1.379 | 7.95E-05 | A7M90_03345 |
| V5VF22 | Uncharacterized protein | -1.355 | 1.41E-06 | A7M90_06705 |
| A0A059ZMV3 | Ethanolamine utilization protein EutQ | -1.35 | 0.0030571 | ABR2091_2644 |
| V5VEN3 | Transposase | -1.323 | 0.00183548 | A7M90_06685 |
| A0A0D5YLD0 | Uncharacterized protein | -1.275 | 0.01944493 | ABUW_3439 |
| V5VHC7 | Cytokinin riboside 5'-monophosphate phosphoribohydrolase | -1.261 | 0.02447429 | yvdD_1 |
| V5VAU0 | DUF4845 domain-containing protein | -1.255 | 1.32E-05 | A7M90_09940 |
| V5VBB1 | DUF2147 domain-containing protein | -1.252 | 0.00078086 | ABR2091_3185 |
| A0A059ZG43 | Methyltransferase | -1.243 | 2.86E-06 | ABUW_1469 |
| V5VCN3 | Protocatechuate 3,4-dioxygenase subunit alpha | -1.211 | 0.00039937 | pcaG |
| V5VHP0 | DUF4199 domain-containing protein | -1.187 | 0.02012464 | ABR2091_0186 |
| A0A059ZPD0 | Acetoin dehydrogenase operon transcriptional activator AcoR | -1.175 | 0.00247979 | acoR_1 |
| A0A0D5YGZ4 | 3-dehydroquinate dehydratase | -1.16 | 3.34E-07 | aroD |
| A0A0D5YGK6 | GTPase | -1.132 | 0.00106318 | ABUW_1139 |
| Q2FD35 | Aminoglycoside 3'-phosphotransferase | -1.119 | 1.76E-06 | strA |
| A0A0D5YIR0 | Catalase | -1.108 | 0.01266128 | ABUW_2059 |
| V5VAE3 | EsvJ | -1.09 | 0.16405168 | lytM_1 |
| A0A0D5YKX4 | ComEA family DNA-binding protein | -1.051 | 0.00013033 | ABUW_3299 |
| A0A0D5YMZ1 | Uncharacterized protein | -1.028 | 0.00901267 | ABUW_3577 |
| A0A0D7TFH6 | Cold-shock DNA-binding domain protein | -1.017 | 0.00348418 | csp2 |
| A0A059ZLA3 | Dihydrodipicolinate synthase family protein | -1.017 | 0.00013806 | ABUW_3682 |
| A0A0D5YI79 | STAS domain-containing protein | -1.01 | 0.00405292 | ABUW_1760 |
| A0A0D5YJK9 | Taurine ABC transporter substrate-binding protein | -0.993 | 0.000337 | tauA |
| A0A0D5YJY3 | Hotdog fold thioesterase | -0.97 | 0.02480247 | paaI1 |
| Q2FD60 | Amino acid permease | -0.956 | 0.00026405 | cycA_2 |
| A0A0D5YIG9 | 3-dehydroshikimate dehydratase | -0.92 | 0.00083402 | quiC |
| A0A0D5YEF0 | Lytic transglycosylase, catalytic | -0.918 | 0.00095406 | ABUW_0925 |
| A0A0D5YME8 | TonB-dependent receptor | -0.865 | 8.89E-05 | ABUW_3403 |
| V5VAT6 | ABC transporter permease | -0.862 | 4.23E-05 | sbp_1 |
| A0A0D8ENM3 | Energy transducer TonB | -0.859 | 0.00019991 | ABUW_2982 |
| A0A0D5YK34 | Heme oxygenase | -0.858 | 0.00051794 | ABUW_2981 |
| A0A0E1FJA9 | LemA family | -0.858 | 0.01645243 | ABUW_0534 |
| A0A0D5YJW7 | TonB dependent outer membrane siderophore receptor protein | -0.857 | 3.83E-05 | pfeA |
| A0A0D8GDK3 | AraC family transcriptional regulator | -0.855 | 0.00096192 | virS_10 |
| V5VHL8 | glutathione transferase | -0.848 | 8.12E-06 | dcmA |
| A0A0D7TN80 | DUF454 domain-containing protein | -0.842 | 0.00147567 | ybaN_1 |
| A0A0D5YDS1 | Uncharacterized protein | -0.839 | 0.0004367 | ABUW_0263 |
| A0A0D5YKF5 | 2Fe-2S iron-sulfur cluster binding domain-containing protein | -0.834 | 0.02131405 | benC |
| V5VGD1 | 'Cold-shock' DNA-binding domain protein | -0.817 | 0.01594103 | cspV |
| A0A0D5YHT1 | PEGA domain-containing protein | -0.814 | 0.00017265 | ABUW_1769 |
| A0A0D5YKC4 | Aldehyde dehydrogenase | -0.811 | 1.36E-05 | gabD3 |
| V5VAX2 | Amino acid transporter | -0.807 | 0.14884857 | rocC_2 |
| A0A0D5YE13 | Uncharacterized protein | -0.807 | 0.00288607 | ABUW_0764 |
| V5VIF8 | Alkanesulfonate monooxygenase | -0.801 | 0.00028768 | ssuD |
| A0A0D5YI68 | Protocatechuate 3,4-dioxygenase subunit beta | -0.798 | 7.95E-06 | pcaH |
| A0A0D5YKS5 | Salicylaldehyde dehydrogenase | -0.79 | 0.02562639 | ydcW |
| A0A0E1FH39 | DNA-binding protein | -0.781 | 0.0005037 | ABR2091_2400 |
| A0A0D5YL43 | Transcriptional Regulator, LysR family | -0.776 | 0.03584835 | ABUW_3344 |
| A0A0D5YL73 | Hemophilin receptor | -0.774 | 0.00049329 | hphR |
| V5VEI6 | HAS ABC exporter outer membrane component | -0.772 | 3.24E-06 | bepC_1 |
| A0A0D5YMQ6 | Fatty acid desaturase | -0.768 | 0.07266641 | ABUW_3842 |
| A0A0D5YGR5 | protein-N(pi)-phosphohistidine--D-fructose phosphotransferase | -0.767 | 0.02576984 | fruA |
| A0A081HAI4 | DUF2846 domain-containing protein | -0.763 | 0.01667883 | A7M90_00590 |
| V5VA62 | Aspartyl/glutamyl-tRNA(Asn/Gln) amidotransferase subunit C | -0.757 | 0.06268032 | gatC |
| A0A0D5YHU5 | Iron-regulated membrane protein | -0.753 | 0.00532399 | ABUW_2167 |
| A0A0D5YLG6 | Heme oxygenase-like protein | -0.75 | 0.00287779 | ABUW_3351 |
| A0A0D5YDJ8 | 4-aminobutyrate transaminase | -0.749 | 4.20E-07 | gabT |
| A0A0D5YDD1 | TonB-dependent receptor protein | -0.746 | 0.00044925 | ABUW_0143 |
| A0A098SFJ9 | Alpha-ketoglutarate-dependent dioxygenase AlkB | -0.746 | 0.07292763 | alkB |
| A0A098SH53 | ATP-binding cassette domain-containing protein | -0.745 | 0.00148566 | tauB |
| A0A0D5YF97;A0A0D5YGP7 | Uncharacterized protein | -0.737 | 0.05492797 | ABUW_0792;ABUW_1310 |
| V5VDI9 | Iron-containing redox enzyme family protein | -0.731 | 0.00071046 | A7M90_12265 |
| V5V960 | Endonuclease | -0.73 | 0.00017102 | A7M90_15425 |
| V5VF83 | Cyclic AMP receptor protein | -0.73 | 0.00039371 | crp |
| V5VFD0 | Transcriptional regulator | -0.721 | 0.00116372 | bigR_1 |
| V5VHL0 | Aldolase | -0.721 | 0.00175543 | A7M90_17700 |
| A0A0D5YKZ7 | Uncharacterized protein | -0.716 | 0.00195592 | ABUW_2833 |
| A0A0D5YK64 | SMI1/KNR4 family protein | -0.71 | 0.0724303 | ABUW_2611 |
| A0A0D5YEX0 | DNA replication protein DnaD | -0.695 | 0.01336001 | ABUW_0751 |
| A0A0D5YIC0 | D-methionine transport protein | -0.69 | 0.00986934 | metQ1 |
| G1C763 | D-ala-D-ala-carboxypeptidase penicillin-binding protein | -0.682 | 2.01E-05 | dacD |
| A0A0D5YC67 | Urocanate hydratase | -0.677 | 1.39E-05 | hutU |
| A0A0D5YE30 | DNA 5'-3' helicase | -0.668 | 0.01031339 | ABUW_0752 |
| A0A0D5YL06 | Glucarate dehydratase | -0.667 | 0.0144576 | gudD |
| A0A059ZT40 | Cytochrome B | -0.667 | 0.02463006 | cybC |
| Q5DL40 | CysA | -0.659 | 0.00051862 | cysA |
| V5VDA3 | Iron transporter | -0.657 | 0.08943464 | A7M90_12010 |
| A0A077GKA3 | Carnitine monooxygenase reductase subunit | -0.655 | 0.00794438 | cntB |
| V5V9W5 | Acyl-CoA desaturase | -0.653 | 0.00091493 | A7M90_17285 |
| V5VFQ0 | CitMHS family transporter | -0.651 | 0.00053054 | citN |
| V5VFS6 | Aspartate/glutamate leucyltransferase | -0.648 | 0.01525501 | ate |
| A0A0D5YIA7 | Uncharacterized protein | -0.647 | 0.00052446 | ABUW_2317 |
| A0A0D5YJ00 | Uncharacterized protein | -0.647 | 0.04234809 | ABUW_2144 |
| A0A059ZUC3 | Lipoprotein | -0.644 | 0.00565246 | ABUW_1597 |
| V5VIX7 | Phosphodiesterase | -0.642 | 0.01048538 | cpdA |
| A0A0D5YM34 | NADH-quinone oxidoreductase subunit J | -0.639 | 0.05473809 | nuoJ |
| V5VHI6 | Phosphomethylpyrimidine synthase | -0.636 | 0.00132884 | thiC |
| V5XVS7 | Epoxyqueuosine reductase | -0.63 | 0.02304019 | queG |
| V5VGN6 | Large ribosomal subunit protein bL35 | -0.625 | 0.00251573 | rpmI |
| A0A059ZGT5 | HTH-type transcriptional regulator HmrR | -0.622 | 0.01203487 | hmrR_3 |
| V5VEW6 | diguanylate cyclase | -0.617 | 0.06421382 | cph2_5 |
| A0A0D5YJ17 | Outer-membrane receptor for Fe(III)-coprogen,Fe(III)-ferrioxamine B and Fe(III)-rhodotrulic acid | -0.609 | 2.55E-05 | ABUW_2165 |
| A0A0D5YMR3 | Heme chaperone HemW | -0.605 | 0.02885861 | hemW |
| V5VGS0 | Nuclear transport factor 2 family protein | -0.604 | 0.13814088 | A7M90_09040 |
| A0A0D5YGI5 | Uncharacterized protein | -0.604 | 0.0625741 | ABUW_0788 |
| A0A0E1FKE5 | Ureidoglycolate hydrolase | -0.603 | 0.00342734 | allA |
| A0A0D5YGJ2 | Heme oxygenase-like protein | -0.602 | 0.00600547 | ABUW_1693 |
| A0A0D5YM90 | TonB-dependent siderophore receptor | -0.602 | 0.00134671 | ABUW_3792 |
| A0A1A9LAX8 | 2OG-Fe(II) oxygenase superfamily protein | -0.592 | 0.00047203 | piuC |
| V5V813 | 4-hydroxyphenylpyruvate dioxygenase | -0.588 | 8.58E-05 | hppD |
| A0A0D5YHL8 | TonB-dependent siderophore receptor | -0.588 | 5.51E-05 | ABUW_1655 |
| A0A0D5YEB8 | Sulfate ABC transporter substrate-binding protein | -0.587 | 0.00033807 | sbp |
| A0A0D5YHN9 | Uncharacterized protein | -0.583 | 0.00155498 | ABUW_2058 |

***Table S4c. Proteomic analysis of AB5075^-cavA-hfq^ expressed proteins with increased relative abundance to AB5075^-cavA^.*** *Strains were cultured to OD_600nm_=0.5 in ID-CAMHB at 37ºC.*

| **UniProt ID** | **Protein Descriptions** | **Log_2_ FC** | **Pvalue** | **Genes** |
| --- | --- | --- | --- | --- |
| V5VES8 | Lysozyme | 4.015 | 0.001849 | tssE |
| A0A0D5YKR2 | Uncharacterized protein | 2.652 | 0.10124241 | ABUW_2797 |
| A0A0D5YHJ2 | DUF4882 domain-containing protein | 2.442 | 9.29E-05 | ABUW_2056 |
| A0A0D5YEW8 | Coproporphyrinogen III oxidase | 2.123 | 0.00047388 | ABUW_0684 |
| V5V8Y6 | GDT1 family protein | 2.108 | 0.13165937 | A7M90_07160 |
| A0A0D5YGD3 | Peptidase C39 | 2.026 | 0.00555837 | ABUW_1225 |
| E9L098 | StrB | 2.011 | 0.15725188 | strB |
| A0A059ZLA3 | Dihydrodipicolinate synthase family protein | 2.002 | 0.11403528 | ABUW_3682 |
| U5QBL2 | TraU | 1.940 | 0.01308073 | traU |
| A0A0D5YFJ5 | Biofilm-associated protein | 1.931 | 2.04E-07 | ABUW_0916 |
| V5VIK3 | DUF1345 domain-containing protein | 1.910 | 0.00201145 | A7M90_00965 |
| A0A0D5YL68 | RHS repeat protein | 1.892 | 6.88E-05 | ABUW_2831 |
| A0A0C0ABZ5 | Uncharacterized protein | 1.883 | 0.00323521 | ABUW_2829 |
| Q2FD95 | Acriflavine resistance protein A | 1.856 | 1.13E-07 | adeI |
| V5V9B8 | DNA nickase | 1.816 | 0.00023574 | A7M90_14505 |
| A0A090B4N0 | DUF2147 domain-containing protein | 1.798 | 0.00023936 | A7M90_06480 |
| V5VHP0 | DUF4199 domain-containing protein | 1.756 | 0.00027894 | ABR2091_0186 |
| Q2FD94 | Efflux pump membrane transporter | 1.720 | 3.53E-08 | adeJ |
| A0A0D5YEN8 | Alpha/beta fold hydrolase | 1.689 | 2.06E-10 | ABUW_0685 |
| A0A0D5YIW1 | Uncharacterized protein | 1.665 | 0.00057699 | ABUW_2145 |
| A0A0D5YMI7 | Matrixin family metalloprotease | 1.652 | 0.13742203 | ABUW_3880 |
| V5VBI8 | Transport permease protein | 1.636 | 0.06217058 | yadH |
| A0A0D5YL31 | Uncharacterized protein | 1.628 | 1.44E-06 | ABUW_2940 |
| V5VC75 | Fimbrial major subunit CsuA/B family protein | 1.626 | 0.00034819 | A7M90_01585 |
| A0A0D5YI82 | Amidohydrolase | 1.607 | 0.05185471 | ABUW_1910 |
| A0A0D5YKI0 | DUF4142 domain-containing protein | 1.580 | 1.43E-07 | ABUW_2679 |
| Q24LT6 | AdeK | 1.516 | 1.20E-06 | adeK |
| A0A081GYG5 | Cation:dicarboxylase symporter family transporter | 1.485 | 3.16E-06 | ABUW_2358 |
| A0A0D5YGE2 | Fimbria/pilus periplasmic chaperone | 1.472 | 3.11E-05 | ABUW_1632 |
| A0A0D5YKY0 | Lipoprotein | 1.471 | 1.06E-05 | ABUW_2811 |
| A0A059ZUC3 | Lipoprotein | 1.459 | 0.00764107 | ABUW_1597 |
| A0A0D5YJK8 | OmpA family protein | 1.430 | 0.00107736 | ABUW_2730 |
| U5QBF7 | Uncharacterized protein | 1.408 | 0.00527861 | ABUW_4029 |
| A0A0D8V3K3 | HTH-type transcriptional repressor NemR | 1.398 | 0.00014652 | nemR_3 |
| A0A0D5YH96 | Circularly permuted type 2 ATP-grasp protein | 1.396 | 8.57E-05 | ABUW_1540 |
| A0A0D5YIB7 | Type III restriction enzyme, res subunit | 1.380 | 1.31E-06 | ABUW_1958 |
| A0A0D7QWA3 | HTH-type transcriptional regulator DmlR | 1.380 | 1.60E-09 | dmlR_2 |
| A0A0D5YKA4 | Catalase-related peroxidase | 1.342 | 0.00013136 | ABUW_2504 |
| V5VDM1 | Fimbrial protein | 1.342 | 0.06792229 | ABUW_2313 |
| A0A0D5YJP6 | RHS repeat-associated core domain-containing protein | 1.341 | 4.53E-06 | ABUW_2830 |
| A0A0D5YJC6 | GCN5-related N-acetyltransferase | 1.330 | 0.05862357 | ABUW_2635 |
| A0A0D5YKS8 | FilC | 1.290 | 7.86E-05 | filC |
| A0A0D5YKP8 | Amidohydrolase | 1.272 | 3.58E-06 | ABUW_3204 |
| A0A0D5YP62 | Bcr/CflA family efflux transporter | 1.272 | 0.07398522 | cmlA |
| A0A0D5YLG6 | Heme oxygenase-like protein | 1.266 | 8.03E-05 | ABUW_3351 |
| V5VBZ0 | Metalloprotease | 1.256 | 4.20E-06 | ABUW_0888 |
| A0A0D5YEK2 | Polysaccharide biosynthesis protein | 1.253 | 0.00523159 | ABUW_0934 |
| A0A0B3YZD4 | Thiol:disulfide interchange protein | 1.249 | 0.00159588 | dsbC_3 |
| U5QEZ8 | TraK | 1.225 | 0.01075114 | traK |
| A0A0B4A3C4 | Fumarate lyase | 1.216 | 0.00021473 | aspA |
| A0A0D5YLE7 | Lipoprotein | 1.215 | 8.33E-05 | ABUW_3459 |
| A0A0D5YM95 | SDR family NAD(P)-dependent oxidoreductase | 1.212 | 3.41E-05 | ABUW_3798 |
| A0A0D5YHN9 | Uncharacterized protein | 1.203 | 0.00012544 | ABUW_2058 |
| U5QEU2 | Lipoprotein | 1.187 | 0.04297918 | ABUW_4035 |
| A0A059ZMM7 | DUF1311 domain-containing protein | 1.174 | 5.11E-06 | A7M90_00885 |
| V5VEP6 | Lipoprotein | 1.166 | 2.75E-06 | A7M90_03345 |
| A0A0D5YN64 | TonB-dependent copper receptor | 1.166 | 4.35E-09 | ABUW_3719 |
| A0A0D5YH40 | DUF403 domain-containing protein | 1.161 | 2.21E-06 | ABUW_1541 |
| A0A0D5YKR7 | Pyrroline-5-carboxylate reductase | 1.159 | 0.00010393 | proC |
| A0A0D5YC67 | Urocanate hydratase | 1.143 | 0.0011333 | hutU |
| V5VGH4 | proton-translocating NAD(P)(+) transhydrogenase | 1.139 | 0.01654434 | pntA |
| A0A059ZTG3 | Uncharacterized protein | 1.129 | 0.00018858 | A7M90_15715 |
| B0FND5 | dihydrofolate reductase | 1.123 | 0.00402312 | dfrVII |
| A0A0D5YDJ8 | 4-aminobutyrate transaminase | 1.123 | 0.00051309 | gabT |
| A0A0D5YIG5 | 4-hydroxybenzoate transporter | 1.122 | 0.04806895 | pcaK1 |
| V5VA61 | Type II secretion system protein H | 1.111 | 0.00108242 | ABR2091_3081 |
| U5QBB1 | DUF1173 domain-containing protein | 1.110 | 0.09987588 | A7M90_15735 |
| A0A0D5YKC1 | N-acetyltransferase | 1.106 | 3.20E-05 | ABUW_2620 |
| A0A0E1FK68 | EamA family transporter | 1.091 | 0.0199452 | ABUW_0215 |
| V5V8K4 | Transporter | 1.084 | 0.00252535 | A7M90_07820 |
| X2FEH6 | Bacterial conjugation TrbI-like protein | 1.081 | 0.00509468 | traB |
| A0A0D5YKG0 | DUF4882 domain-containing protein | 1.074 | 0.00038704 | ABUW_2658 |
| U5QBD8 | TraE | 1.073 | 0.00268716 | traE |
| A0A0D5YDM0 | Sulfate transporter | 1.072 | 4.88E-05 | ABUW_0259 |
| U5QES9 | Membrane lipoprotein lipid attachment site | 1.062 | 0.04467313 | traV |
| C5HUY1 | beta-lactamase | 1.060 | 0.00517351 | blaGES-11 |
| U5QBG2 | Toxin | 1.044 | 1.65E-06 | ABUW_4003 |
| A0A0D8GPM3 | Lipoprotein, putative | 1.043 | 0.00333211 | ABUW_2656 |
| A0A0D5YFZ2 | Acyl-CoA dehydrogenase | 1.031 | 0.00018483 | ABUW_1467 |
| A0A0D8GPT4 | DUF1049 domain-containing protein | 1.029 | 0.0705493 | ABUW_2674 |
| A0A0D5YHQ4 | 5'-nucleotidase | 1.026 | 0.00070306 | ABUW_2073 |
| U5QBN9 | Uncharacterized protein | 1.025 | 0.00083595 | ABUW_4082 |
| A0A0D5YIH8 | TetR family transcriptional regulator | 1.017 | 0.00104673 | ABUW_2325 |
| A0A0D5YK41 | L-serine dehydratase | 1.017 | 0.00063505 | sdaA |
| A0A0D5YN51 | 2'-aminoglycoside nucleotidyltransferase AadB | 1.003 | 0.0091837 | aadB |
| A0A0D5YJS4 | DUF3139 domain-containing protein | 1.002 | 0.02462622 | ABUW_2795 |
| A0A0D5YLM2 | Cation:proton antiporter | 0.998 | 0.00088614 | phaD |
| V5VCT5 | Ferrous iron transporter B | 0.996 | 0.01290626 | hifA |
| U5QBH9 | TraW | 0.992 | 0.0110538 | traW |
| A0A059ZGT5 | HTH-type transcriptional regulator HmrR | 0.988 | 0.13112914 | hmrR_3 |
| A0A0D5YF78 | Uncharacterized protein | 0.988 | 0.00025317 | ABUW_0771 |
| A0A0X3QPX2 | Glyoxalase | 0.974 | 3.56E-06 | A7M90_16310 |
| X2FIS0 | TraH | 0.971 | 0.0115711 | traH |
| U5QBD2 | Uncharacterized protein | 0.970 | 1.00E-05 | ABUW_4033 |
| A0A0D5YGW6 | methylmalonate-semialdehyde dehydrogenase (CoA acylating) | 0.969 | 0.00010014 | mmsA2 |
| A0A0D5YLK1 | Trehalose 6-phosphate phosphatase | 0.945 | 0.10824616 | otsB |
| A0A0D5YDS8 | Uncharacterized protein | 0.942 | 2.05E-05 | ABUW_0709 |
| A0A0D7U1B8 | ABC transporter ATP-binding protein | 0.931 | 1.65E-05 | gsiA_3 |
| A0A0D5YH21 | Omega-amino acid--pyruvate aminotransferase | 0.914 | 0.00528348 | ABUW_1463 |
| A0A0D5YNZ3 | Matrixin superfamily | 0.911 | 0.02010104 | ABUW_3881 |
| A0A315RS97 | L-lactate permease | 0.906 | 0.01868983 | lldP |
| U5QBM2 | Signal peptidase I | 0.898 | 0.00045279 | lepB |
| A0A081H4A8 | ABC transporter permease | 0.897 | 0.00034744 | gsiC |
| A0A0D5YFT2 | Crispr-associated protein, Csy4 family | 0.890 | 0.0001394 | csy4 |
| A0A098SC94 | GFA family protein | 0.888 | 0.12781264 | ABUW_0479 |
| A0A0D5YGZ0 | 3-oxoadipate enol-lactonase | 0.881 | 0.00011882 | pcaD1 |
| V5XWX9 | GrpB family protein | 0.878 | 0.0003852 | ABUW_2288 |
| U5QBP3 | Uncharacterized protein | 0.878 | 0.00454834 | A7M90_15640 |
| V5VDL8 | General secretion pathway protein G | 0.876 | 0.12486656 | gspG |
| A0A0R4J6G8 | 3-isopropylmalate dehydratase large subunit | 0.874 | 5.64E-05 | leuC |
| A0A0D5YI91 | DUF2314 domain-containing protein | 0.873 | 0.0004327 | ABUW_1920 |
| A0A0D5YCT2 | DUF805 domain-containing protein | 0.871 | 0.0013095 | ABUW_0248 |
| A0A0D5YKZ7 | Uncharacterized protein | 0.861 | 0.01091275 | ABUW_2833 |
| U5QBN0 | Uncharacterized protein | 0.860 | 0.11870629 | ABUW_4031 |
| A0A0D5YK75 | Peroxiredoxin | 0.858 | 0.00590938 | ABUW_3027 |
| A0A059ZML9 | Uncharacterized protein | 0.846 | 0.00117327 | A7M90_02010 |
| A0A0D5YJT9 | DcaP-like protein | 0.841 | 0.06853512 | ABUW_2329 |
| A0A0D5YJ25 | SMI1/KNR4 family protein | 0.839 | 3.64E-05 | ABUW_2612 |
| U5QBA4 | TraN | 0.839 | 0.00114904 | traN |
| A0A0D5YIC2 | Acetyltransferase | 0.837 | 0.00023862 | ABUW_1964 |
| Q7AZE4 | Dihydropteroate synthase | 0.837 | 0.01656499 | sul1 |
| A0A0D5YJL2 | Uncharacterized protein | 0.829 | 0.00277643 | ABUW_2387 |
| V5VHV5 | MAPEG family | 0.827 | 0.00010325 | A7M90_17675 |
| V5VEN3 | Transposase | 0.826 | 0.08348198 | A7M90_06685 |
| V5VEI5 | Phenylacetic acid degradation protein | 0.816 | 0.06357277 | A7M90_05745 |
| A0A077GKT8 | Anthranilate synthase | 0.815 | 0.00031366 | A7M90_11190 |
| A0A090BC97 | FilD | 0.814 | 0.06120202 | fadL |
| A0A0D5YL84 | Uncharacterized conserved protein | 0.814 | 1.37E-06 | ABUW_2995 |
| A0A0D5YG02 | Csy3 family crispr-associated protein | 0.813 | 3.78E-06 | csy3 |
| Q2FD35 | Aminoglycoside 3'-phosphotransferase | 0.810 | 0.00237031 | strA |
| A0A0D5YEF7 | Biofilm-associated protein | 0.808 | 8.37E-05 | ABUW_0885 |
| A0A0D8GPZ2 | Lipoprotein, putative | 0.807 | 0.00089697 | ABUW_2654 |
| V5VBM8 | DUF2171 domain-containing protein | 0.806 | 0.00250679 | A7M90_04275 |
| A0A0J8TTU0 | Transmembrane protein | 0.801 | 0.00029907 | ABUW_0243 |
| A0A0D5YCF4 | DUF1311 domain-containing protein | 0.798 | 1.47E-05 | ABUW_0120 |
| A0A0D5YIT6 | Aminotransferase | 0.797 | 4.42E-06 | ABUW_2519 |
| A0A0D5YIF0 | Amidohydrolase | 0.792 | 0.00012786 | ABUW_2373 |
| V5V964 | Acetyl-CoA hydrolase | 0.789 | 0.00058949 | cat1 |
| A0A0D5YMI9 | Copper-translocating P-type ATPase | 0.785 | 0.00473898 | actP1 |
| V5VER8 | Divalent metal cation transporter | 0.781 | 0.0001907 | A7M90_06400 |
| V5VGD1 | 'Cold-shock' DNA-binding domain protein | 0.781 | 0.01749304 | cspV |
| A0A0D5YJY0 | Lipoprotein | 0.780 | 0.00017984 | A7M90_09100 |
| U5QBF1 | Conjugal transfer protein TraG | 0.778 | 0.01252528 | traG |
| U5QEP9 | AAA-like domain protein | 0.775 | 0.0047097 | traC |
| A0A0D5YH31 | Alpha/beta fold hydrolase | 0.775 | 0.08225812 | cpo |
| A0A0D5YF62 | Ferric acinetobactin transport system permease | 0.768 | 0.02323266 | bauC |
| A0A0D5YMN4 | Bacterial transferase hexapeptide (Three repeats) family protein | 0.767 | 2.08E-07 | ABUW_3822 |
| A0A0D5YFE8 | DUF4850 domain-containing protein | 0.763 | 0.07342181 | ABUW_0708 |
| A0A0D5YI64 | Beta-ketoadipyl-CoA thiolase | 0.763 | 8.96E-05 | pcaF1 |
| A0A0D5YIC8 | Lipoprotein | 0.761 | 0.00054338 | ABUW_1919 |
| A0A0E1JRA6 | Catalase | 0.758 | 0.02523141 | katE |
| A0A0D5YK99 | Aldose 1-epimerase | 0.756 | 0.00105576 | ABUW_2933 |
| A0A0D5YJB4 | Putative class A beta-lactamase | 0.754 | 0.00070014 | ABUW_2619 |
| A0A0D5YHW1 | Uncharacterized protein | 0.752 | 4.31E-05 | ABUW_1752 |
| A0A0D5YKJ9 | Biotin carboxylase | 0.752 | 0.00069339 | bccA |
| A0A059ZTI4 | Secreted protein | 0.752 | 0.01783766 | A7M90_15590 |
| A0A0D5YJA2 | 5-oxoprolinase/urea amidolyase family protein | 0.751 | 3.46E-06 | ABUW_2604 |
| V5VG26 | NADH-quinone oxidoreductase subunit N | 0.748 | 0.07605588 | nuoN |
| A0A0D5YDA8 | Lipoprotein | 0.745 | 3.89E-05 | ABUW_0118 |
| V5VA62 | Aspartyl/glutamyl-tRNA(Asn/Gln) amidotransferase subunit C | 0.742 | 0.00126559 | gatC |
| V5VE93 | DUF6438 domain-containing protein | 0.742 | 0.00440273 | A7M90_02595 |
| A0A0D5YDI2 | diguanylate cyclase | 0.741 | 0.01824943 | ABUW_0188 |
| A0A0D5YIF9;V5VCY7 | 3-oxoacid CoA-transferase subunit A | 0.739 | 0.00491741 | pcaI1;pcaI_2 |
| A0A0D5YFC3 | Endolytic peptidoglycan transglycosylase RlpA | 0.738 | 0.01416196 | rlpA |
| A0A0D5YK59 | 5-oxoprolinase subunit A | 0.729 | 0.00309834 | pxpA |
| A0A0D5YJ87 | Oxidoreductase | 0.729 | 0.05880372 | ABUW_2122 |
| A0A0D5YD70 | DUF4256 domain-containing protein | 0.728 | 0.00816115 | ABUW_0480 |
| A0A0D8GDK3 | AraC family transcriptional regulator | 0.726 | 0.14333745 | virS_10 |
| A0A0D5YF96 | VOC family protein | 0.721 | 0.00575638 | ABUW_0646 |
| A0A0D5YJ75 | Heme-binding protein A | 0.708 | 5.26E-05 | ABUW_2225 |
| A0A0D5YJX5 | AAA family ATPase | 0.703 | 0.00197737 | ABUW_2810 |
| A0A0D5YIG8 | PH domain-containing protein | 0.702 | 0.0122516 | ABUW_1963 |
| A0A059ZFG8 | Uncharacterized protein | 0.702 | 0.00288658 | ABUW_3070 |
| A0A0B9W852 | Heme-binding protein | 0.695 | 2.42E-05 | ABUW_2226 |
| V5VEA4 | PAAR domain-containing protein | 0.694 | 0.00011983 | ABR2091_1303 |
| A0A0D5YK55 | 3-methylglutaconyl-CoA hydratase | 0.687 | 0.00011824 | mgh |
| U5QBJ7 | TraD | 0.684 | 8.37E-05 | trwB |
| A0A0D5YDU4 | SH3 domain-containing protein | 0.683 | 0.00023899 | ABUW_0121 |
| A0A0D5YDS1 | Uncharacterized protein | 0.679 | 1.03E-05 | ABUW_0263 |
| A0A0D5YFS0 | Insulinase family protein | 0.678 | 0.00015812 | ABUW_1082 |
| A0A0D5YJ93 | Transcriptional regulator | 0.675 | 0.05237405 | ABUW_2682 |
| V5VHL3 | Glutamine-dependent NAD(+) synthetase | 0.670 | 0.00024363 | nadE |
| A0A0D5YIN6 | Quinoprotein glucose dehydrogenase-B | 0.669 | 5.54E-05 | gdhB1 |
| A0A0D5YFQ0 | UDP-N-acetylglucosamine 2-epimerase (non-hydrolyzing) | 0.664 | 0.02543377 | wecB |
| A0A0D5YN58 | Cysteine synthase | 0.663 | 0.00394838 | cysK |
| A0A0D5YI14 | Iron-containing alcohol dehydrogenase | 0.660 | 0.00070613 | ABUW_1684 |
| A0A0D5YGN2 | Uncharacterized protein | 0.659 | 0.00265405 | ABUW_1295 |
| A0A0D5YGJ2 | Heme oxygenase-like protein | 0.658 | 0.00053399 | ABUW_1693 |
| A0A0D5YKL7 | L-gulonate 3-dehydrogenase | 0.658 | 0.00338357 | ABUW_2623 |
| A0A0D5YHT7 | CENP-V/GFA domain-containing protein | 0.657 | 0.00057594 | ABUW_2157 |
| A0A0D5YEU8 | GFA family protein | 0.657 | 0.00284416 | ABUW_1042 |
| A0A0D5YI49 | Zn-dependent oligopeptidase | 0.649 | 0.00010515 | ABUW_2224 |
| A0A0D5YK78 | Exo-alpha-sialidase | 0.644 | 0.0079562 | ABUW_2626 |
| A0A0D5YLN8 | Glutathione hydrolase proenzyme | 0.640 | 0.00490139 | ggt |
| A0A0D8GPX3 | Uncharacterized protein | 0.639 | 3.12E-05 | ABUW_2675 |
| A0A0D5YH32 | MBL fold metallo-hydrolase | 0.637 | 0.00036076 | ABUW_1847 |
| A0A0D8G6V8 | KTSC domain-containing protein | 0.636 | 0.00604136 | ABUW_1262 |
| A0A0D5YDK9 | histidine kinase | 0.633 | 0.00922362 | ABUW_0585 |
| A0A086I1Y6 | Succinate dehydrogenase cytochrome b556 subunit | 0.633 | 0.00154113 | sdhC |
| A0A0D5YH17 | 3-oxoacid CoA-transferase subunit B | 0.631 | 0.00712808 | pcaJ1 |
| C0L0K5 | Carbapenem-associated resistance protein | 0.630 | 2.23E-05 | carO |
| A0A059ZI87 | DUF808 domain-containing protein | 0.624 | 0.00579103 | A7M90_18835 |
| A0A0D5YJD4 | Phosphodiester glycosidase domain-containing protein | 0.623 | 0.01104947 | ABUW_2291 |
| U5QET5 | YcxB family protein | 0.619 | 0.0313593 | ABUW_4008 |
| A0A0D5YJK4 | Glutathione S-transferase | 0.614 | 0.00011811 | gst |
| A0A0D5YEE4 | Metal-dependent hydrolase of the beta-lactamase superfamily | 0.611 | 0.00042801 | ABUW_0920 |
| A0A077GRI0 | Aldehyde dehydrogenase | 0.610 | 8.75E-05 | gapN |
| A0A059ZBD3 | DUF421 domain-containing protein | 0.610 | 0.00072098 | A7M90_02515 |
| A0A0D5YD36 | Glutamate racemase | 0.606 | 0.01114074 | murI1 |
| A0A0D5YEA9 | NAD-dependent DNA ligase | 0.605 | 3.84E-05 | ABUW_0553 |
| A0A0D5YIX7 | DUF937 domain-containing protein | 0.604 | 0.0790285 | ABUW_2161 |
| V5V8H4 | Hydrolase | 0.604 | 0.00037165 | ycaC_1 |
| A0A0D5YMQ7 | Acetate kinase | 0.599 | 0.00014437 | ackA |
| A0A098SJS3 | Type II secretion system protein I | 0.598 | 0.00879525 | gspI |
| A0A090C225 | TIGR01777 family oxidoreductase | 0.595 | 4.23E-05 | ABUW_3024 |
| A0A0D5YIA7 | Uncharacterized protein | 0.591 | 0.00120849 | ABUW_2317 |
| A0A0D7TSQ3 | NAD(P)-dependent oxidoreductase | 0.589 | 2.51E-06 | ydfG |
| A0A059ZN16 | Acyltransferase | 0.587 | 9.16E-05 | A7M90_11305 |
| V5VEG3 | General stress protein 39 | 0.585 | 0.00065817 | ydaD |
| A0A1A5UUC6 | Cold shock domain-containing protein | 0.583 | 0.01640734 | cspG |

***Table S4d. Proteomic analysis of AB5075^-cavA-hfq^ expressed proteins with decreased relative abundance to AB5075^-cavA^.*** *Strains were cultured to OD_600nm_=0.5 in ID-CAMHB at 37ºC.*

| **UniProt ID** | **Protein** | **Log_2_ FC** | **Pvalue** | **Gene** |
| --- | --- | --- | --- | --- |
| A0A098SIV4 | Aliphatic sulfonates transport permease protein SsuC | -12.215 | 0.01601826 | tauC |
| A0A0D5YJ02 | Uncharacterized protein | -10.741 | 6.48E-08 | ABUW_1997 |
| V5VCD4 | Multidrug efflux transcriptional regulator AdeN | -10.603 | 7.71E-07 | adeN |
| A0A098SFJ9 | Alpha-ketoglutarate-dependent dioxygenase AlkB | -9.957 | 6.39E-05 | alkB |
| A0A059ZPH9 | MFS transporter | -9.667 | 0.05102885 | ABUW_3317 |
| A0A0D5YLB0 | Alkaline phosphatase | -9.614 | 6.64E-05 | ABUW_3414 |
| A0A0D5YLX0 | DUF3592 domain-containing protein | -8.435 | 0.01620554 | ABUW_3191 |
| A0A0D5YE91 | Energy transducer TonB | -7.744 | 3.46E-09 | ABR2091_3221 |
| A0A0D8G9M9 | PilW family protein | -7.673 | 0.10313484 | pilW |
| A0A0D5YN54 | methylated-DNA--[protein]-cysteine S-methyltransferase | -7.549 | 4.26E-06 | ada |
| A0A0Q2HWF1 | Hemerythrin | -7.278 | 0.00095475 | A7M90_16365 |
| V5VB91 | Csu fimbrial major subunit CsuAB | -6.781 | 7.05E-05 | csuAB |
| A0A090BDJ4 | Metal-dependent hydrolase | -6.576 | 0.02715201 | A7M90_16470 |
| A0A0D5YD32 | Putative DNA modification/repair radical SAM protein | -5.484 | 0.00024073 | ABUW_0369 |
| A0A0D5YM64 | Alginate biosynthesis protein | -5.065 | 0.04727242 | algZ |
| A0A0B9XMY1 | Phage protein | -4.446 | 2.11E-08 | ABUW_0734 |
| A0A0D5YND1 | Response regulator | -4.18 | 0.00035015 | pilR |
| A0A0D5YGW0 | Uncharacterized protein | -4.155 | 1.88E-08 | ABUW_1389 |
| A0A0D5YJW7 | TonB dependent outer membrane siderophore receptor protein | -3.983 | 3.66E-05 | pfeA |
| V5VFG4 | 3-oxoacyl-[acyl-carrier-protein] reductase FabG | -3.953 | 0.01104089 | fabG_5 |
| V5VGV5 | UPF0102 protein A7M90_09240 | -3.903 | 0.01314197 | A7M90_09240 |
| A0A0D5YE49 | Amino acid permease | -3.739 | 3.27E-05 | aroP2 |
| U5QEQ7 | DNA polymerase IV | -3.67 | 4.36E-05 | umuC |
| A0A059ZFB6 | DUF1449 domain-containing protein | -3.658 | 0.00148245 | yqiJ |
| A0A0D5YEX0 | DNA replication protein DnaD | -3.521 | 0.00285024 | ABUW_0751 |
| A0A0D5YK39 | Putative transmembrane sensor protein FecR | -3.51 | 0.00020254 | ABUW_2986 |
| A0A0D5YJQ9 | Lysine exporter protein ArgO | -3.396 | 0.05629334 | ABUW_2850 |
| A0A0D5YG90 | Glycine zipper domain-containing protein | -3.368 | 0.00036855 | A7M90_03805 |
| A0A0D5YDD5 | Uncharacterized protein | -3.341 | 0.04782506 | ABUW_0546 |
| A0A077GZE2 | Lipoprotein | -3.308 | 0.01716216 | ABUW_1788 |
| A0A0D5YFD1 | Chemotaxis protein CheA | -3.23 | 9.60E-05 | pilL |
| A0A0D5YMI4 | FimV domain-containing protein | -3.21 | 1.68E-09 | ABUW_3453 |
| A0A0D5YLU2 | Zinc protease | -3.209 | 5.23E-05 | ABUW_3068 |
| A0A0D5YDC7 | 3'-5' exonuclease | -3.106 | 1.14E-05 | ABUW_0541 |
| A0A059ZFZ5 | NirD/YgiW/YdeI family stress tolerance protein | -3.06 | 6.23E-08 | ygiW |
| A0A0D5YJA0 | Type II secretion system protein J | -3.05 | 0.00095188 | gspJ |
| A0A0D5YH05 | Uncharacterized protein | -2.964 | 0.06056475 | ABUW_1278 |
| A0A0D5YIP3 | Uncharacterized protein | -2.87 | 7.54E-09 | ABUW_2034 |
| A0A0D5YDD1 | TonB-dependent receptor protein | -2.869 | 3.38E-06 | ABUW_0143 |
| A0ABF7PMH8 | Sulfurtransferase | -2.827 | 0.02348922 | tusE |
| A0A0B9VUF6 | Uncharacterized protein | -2.814 | 1.51E-08 | ABUW_0739 |
| V5VH04 | Pilin biogenesis protein | -2.768 | 0.02101397 | epsF_1 |
| A0A077GPV7 | 2-(1,2-epoxy-1,2-dihydrophenyl)acetyl-CoA isomerase | -2.754 | 1.90E-05 | paaB |
| A0A0D5YH74 | Type II secretion system protein E | -2.74 | 0.02047659 | ABUW_1351 |
| A0A0D5YGR0 | Diguanylate cyclase | -2.7 | 0.0002452 | ABUW_1764 |
| A0A0D5YM11 | Prepilin leader peptidase/N-methyltransferase | -2.689 | 0.00943153 | pilD |
| A0A0D5YLK9 | RNA polymerase sigma factor FecI | -2.594 | 0.03156646 | ABUW_2987 |
| A0A0D5YH83 | LysR family transcriptional regulator | -2.59 | 0.05694934 | ABUW_1936 |
| A0A0D5YF17 | Rubredoxin-NAD(+) reductase | -2.48 | 0.02853435 | ABUW_1151 |
| V5VA30 | Alkaline phosphatase synthesis transcriptional regulatory protein PhoP | -2.45 | 1.70E-12 | pilG |
| A0A0D5YME8 | TonB-dependent receptor | -2.435 | 1.41E-10 | ABUW_3403 |
| A0A0D5YD12 | Uncharacterized protein | -2.412 | 9.58E-05 | ABUW_0344 |
| V5VI75 | 2-amino-4-hydroxy-6-hydroxymethyldihydropteridine pyrophosphokinase | -2.391 | 0.00020831 | folK |
| A0A0D5YHX5 | DUF2750 domain-containing protein | -2.36 | 0.06250653 | ABUW_1784 |
| A0A0D5YK64 | SMI1/KNR4 family protein | -2.294 | 0.00017662 | ABUW_2611 |
| A0A0D5YJ17 | Outer-membrane receptor for Fe(III)-coprogen,Fe(III)-ferrioxamine B and Fe(III)-rhodotrulic acid | -2.292 | 1.14E-07 | ABUW_2165 |
| A0A0D5YEH5 | Uncharacterized protein | -2.271 | 0.00691436 | ABUW_0547 |
| A0A0D5YMT3 | Uncharacterized protein | -2.261 | 0.00117228 | ABUW_4081 |
| A0A0D5YFN7 | Phage family protein | -2.233 | 1.46E-06 | ABUW_0780 |
| A0A0D5YJN1 | Aromatic amino acid transport protein AroP | -2.22 | 0.01456522 | ABUW_2808 |
| A0A0D5YHJ9 | CsuD | -2.219 | 0.02050059 | csuD |
| V5VHC7 | Cytokinin riboside 5'-monophosphate phosphoribohydrolase | -2.2 | 0.04495725 | yvdD_1 |
| A0A089XXH6 | Chloramphenicol resistance pump cmr | -2.135 | 2.32E-05 | mdfA |
| A0A0D5YJV9 | SPOR domain-containing protein | -2.134 | 0.00016582 | ABR2091_0979 |
| A0A0D5YD94 | Acyl-CoA dehydrogenase | -2.13 | 0.00405569 | fadE |
| A0A0D8FIG7 | Uncharacterized protein | -2.124 | 0.002013 | ABUW_1719 |
| A0A0D5YCC2 | diguanylate cyclase | -2.078 | 0.05267215 | ABUW_0137 |
| A0A0D5YM34 | NADH-quinone oxidoreductase subunit J | -2.041 | 3.56E-05 | nuoJ |
| A0A0D5YDN0 | Gamma-glutamylcyclotransferase | -2.033 | 5.78E-05 | ABUW_0657 |
| A0A0D5YNN7 | AMP-binding protein | -2.031 | 0.00037973 | ABUW_3773 |
| A0A6H3SVU9 | 4-carboxymuconolactone decarboxylase | -2.01 | 0.01444698 | ABR2091_1362 |
| A0A0D5YI79 | STAS domain-containing protein | -2.01 | 0.00062626 | ABUW_1760 |
| V5VGL3 | Fe-S assembly protein IscX | -1.998 | 0.00158787 | iscX |
| A0A0D5YK00 | ABC transporter permease | -1.996 | 0.00526982 | ABUW_2837 |
| A0A0D5YLF8 | Uncharacterized protein | -1.961 | 0.09004097 | ABUW_2932 |
| A0A0D5YFK7 | Amino acid adenylation domain-containing protein | -1.947 | 0.06883816 | ABUW_0932 |
| V5VBK4 | Acetyltransferase | -1.909 | 0.00751639 | ABR2091_2278 |
| A0A0D5YHE2 | Uncharacterized protein | -1.9 | 9.71E-14 | ABUW_1641 |
| A0A0D5YJ46 | ATP-binding cassette domain-containing protein | -1.881 | 0.00022784 | modC |
| A0A0D5YJ38 | Phenylacetic acid degradation bifunctional protein PaaZ | -1.87 | 0.00274041 | paaN |
| A0A0D5YK91 | FMN dependent NADH:quinone oxidoreductase | -1.858 | 0.01810243 | azoR_1 |
| A0A0D5YIE4 | Dihydroxy-acid dehydratase | -1.852 | 0.04274878 | ABUW_1934 |
| A0A0D5YDZ7 | Amidase | -1.815 | 0.00183863 | ABUW_0176 |
| V5VGI6 | NADH-quinone oxidoreductase subunit K | -1.784 | 0.00012028 | nuoK |
| V5V9G1 | NADPH:quinone oxidoreductase MdaB | -1.769 | 0.0001104 | mdaB |
| A0A0D5YDX2 | Toxin-antitoxin system HicB family antitoxin | -1.759 | 0.00080767 | ABUW_0749 |
| Q2FD44 | ArsB | -1.751 | 0.00268178 | arsB |
| A0A0D5YJS0 | Transcriptional regulator, GntR family | -1.749 | 0.00421276 | ABUW_2406 |
| V5VDJ4 | DUF1275 domain-containing protein | -1.721 | 0.15806856 | A7M90_11535 |
| A0A0D5YE21 | DUF4055 domain-containing protein | -1.714 | 0.00872552 | ABUW_0776 |
| A0A0D5YIM8 | Coenzyme PQQ synthesis protein B | -1.706 | 0.00109943 | pqqB |
| V5VGM5 | Integral membrane protein | -1.704 | 0.00649018 | A7M90_00065 |
| A0A059ZFL9 | Outer membrane porin protein | -1.697 | 0.00597489 | porB |
| A0A0D5YGI5 | Uncharacterized protein | -1.686 | 0.00605235 | ABUW_0788 |
| A0A0D5YIT9 | Phage holin family protein | -1.671 | 0.01013295 | ABUW_1971 |
| A0A0D5YKY7 | NTF2 fold immunity protein domain-containing protein | -1.663 | 0.00029611 | ABUW_3190 |
| A0A0D5YEA0 | Uncharacterized protein | -1.65 | 0.00639969 | ABUW_0543 |
| V5VCZ7 | Uncharacterized protein | -1.628 | 6.40E-06 | A7M90_12425 |
| V5VCB8 | GTP cyclohydrolase 1 type 2 | -1.626 | 0.00019035 | A7M90_09810 |
| A0A0D8GAS1 | AraC family transcriptional regulator | -1.625 | 0.06366883 | ABR2091_0176 |
| A0A0D5YEX3 | Inner membrane protein | -1.619 | 0.07337488 | ABUW_0689 |
| A0A0D5YFW4 | Uncharacterized protein | -1.619 | 4.25E-06 | ABUW_1431 |
| V5VG21 | Uncharacterized protein | -1.606 | 0.00059538 | A7M90_13540 |
| A0A0D8G6G4 | Major facilitator superfamily MFS_1 | -1.6 | 0.11330899 | nreB |
| Q2FCW3 | Transposase | -1.599 | 0.01399398 | tnpA |
| A0A0D5YDE6 | Branched-chain amino acid ABC transporter permease | -1.596 | 0.00012468 | ABUW_0199 |
| A0A0D5YHR8 | Release factor glutamine methyltransferase | -1.567 | 0.0023838 | hemK |
| V5V9Q4 | Holin-like protein CidB | -1.565 | 0.07683985 | cidB_2 |
| V5VGG0 | LysM domain protein | -1.559 | 0.02239399 | ABR2091_0553 |
| A0A0D5YK97 | Glutathione-dependent formaldehyde dehydrogenase | -1.559 | 0.00373953 | ABUW_2594 |
| A0A0D5YIA4 | Fumarylacetoacetate hydrolase | -1.557 | 0.01676898 | ABUW_1935 |
| A0A0C0BS03 | Uncharacterized protein | -1.552 | 0.00046969 | ABUW_0813 |
| A0A0D5YJE3 | Ornithine monooxygenase | -1.533 | 1.97E-07 | ABUW_2188 |
| A0A0D5YGT9 | OmpW family protein | -1.501 | 0.0008433 | ABUW_1233 |
| A0A0D5YF35 | Capsid protein | -1.5 | 0.00698219 | ABUW_0581 |
| A0A0D7TN80 | DUF454 domain-containing protein | -1.5 | 0.00087499 | ybaN_1 |
| A0A0D5YE30 | DNA 5'-3' helicase | -1.497 | 0.00156428 | ABUW_0752 |
| A0A059ZGB1 | HTH-type transcriptional regulator gltC | -1.475 | 0.04560835 | ABR2091_1444 |
| A0A0D5YHU5 | Iron-regulated membrane protein | -1.475 | 3.93E-05 | ABUW_2167 |
| A0A0D5YMU6 | OmpW family protein | -1.472 | 5.24E-05 | ABUW_3583 |
| A0A0D5YJR9 | Signal peptide-containing protein | -1.472 | 0.00038796 | ABUW_2860 |
| A0A0D5YK13 | 3-methyladenine DNA glycosylase | -1.455 | 0.00086023 | ABUW_2512 |
| A0A0D5YF70 | Dipeptide/tripeptide permease | -1.452 | 0.00290215 | ABUW_1206 |
| V5VHB1 | Cation:proton antiporter | -1.444 | 0.00106494 | mrpF |
| A0A0D5YLF2 | Transketolase family protein | -1.437 | 0.01105269 | ABUW_2927 |
| V5V8F5 | BCCT family transporter | -1.434 | 9.28E-05 | betT |
| A0A0D5YJI4 | Aminotransferase | -1.432 | 0.00372572 | ABUW_2352 |
| A0A0D5YFR4 | Uncharacterized protein | -1.408 | 0.066501 | ABUW_1388 |
| V5V989 | Cell division protein FtsL | -1.404 | 0.00031221 | ftsL |
| V5VEZ8 | KTSC domain-containing protein | -1.403 | 0.00334529 | A7M90_06840 |
| A0A0D5YKS5 | Salicylaldehyde dehydrogenase | -1.391 | 0.00027658 | ydcW |
| A0A0D5YGR5 | protein-N(pi)-phosphohistidine--D-fructose phosphotransferase | -1.388 | 0.0003036 | fruA |
| A0A0D5YNJ1 | TetR family transcriptional regulator | -1.383 | 0.00432146 | ABUW_3716 |
| A0A0D5YJ33 | Phenylacetate-CoA oxygenase/reductase subunit PaaK | -1.382 | 0.00711955 | paaK |
| V5VBN5 | Alpha/beta fold hydrolase | -1.373 | 0.08177234 | A7M90_04225 |
| A0A0D5YDT5 | Uncharacterized protein | -1.368 | 0.00147661 | ABUW_0111 |
| A0A0D8ENM3 | Energy transducer TonB | -1.367 | 0.00031335 | ABUW_2982 |
| V5VDA3 | Iron transporter | -1.366 | 0.0899054 | A7M90_12010 |
| A0A0D5YDZ6 | Uncharacterized protein | -1.36 | 0.0006955 | ABUW_0737 |
| A0A0D5YDP6 | Uncharacterized protein | -1.353 | 0.02902897 | ABUW_0236 |
| A0A0D5YG17 | CsuE | -1.343 | 0.01220932 | csuE |
| A0A0D5YE00 | Uncharacterized protein | -1.342 | 0.01509443 | ABUW_0742 |
| A0A059ZNI7 | RNA polymerase factor sigma-70 | -1.34 | 0.00606525 | sigX |
| A0A0D5YLZ1 | CoA transferase | -1.339 | 0.04563214 | caiB2 |
| V5VCS3 | Ben operon transcriptional regulator BenM | -1.339 | 0.0788915 | catM |
| A0A059ZTE1 | Uncharacterized protein | -1.332 | 9.55E-05 | ABUW_4026 |
| A0A0D5YFG2 | Toxin-antitoxin system HicB family antitoxin | -1.328 | 7.86E-06 | ABUW_1267 |
| A0A0D5YIR2 | Negative transcriptional regulator | -1.327 | 0.02295571 | ABUW_2405 |
| A0A0D5YGM1 | SHOCT domain-containing protein | -1.327 | 0.0005632 | ABUW_1724 |
| A0A0D5YK27 | Uncharacterized protein | -1.326 | 0.01251525 | ABUW_2862 |
| A0A0D5YDK8 | Cold-shock DNA-binding domain protein | -1.326 | 0.00147243 | ABUW_0242 |
| A0A0D5YP36 | Uncharacterized protein | -1.325 | 0.00258044 | ABUW_6002 |
| A0A0D5YNG6 | Amino acid adenylation | -1.319 | 0.00030219 | ABUW_3770 |
| V5XVS7 | Epoxyqueuosine reductase | -1.317 | 0.00429127 | queG |
| A0A0D5YDP1 | Acetyltransferase, gnat family | -1.316 | 8.56E-05 | ABUW_0230 |
| A0A077GKG8 | Colicin V producing membrane protein | -1.315 | 0.03413771 | cvpA |
| A0A0D5YJB7 | 1,6-dihydroxycyclohexa-2,4-diene-1-carboxylate dehydrogenase | -1.311 | 0.04109287 | benD |
| V5VH28 | Glutaredoxin | -1.31 | 0.00070189 | grxC |
| A0A0D5YKV7 | DUF4265 domain-containing protein | -1.296 | 0.00048089 | ABUW_2796 |
| A0A0D5YCR0 | Type 4 fimbrial biogenesis protein PilN | -1.284 | 7.21E-08 | comN |
| V5VBG3 | Complement control module protein | -1.282 | 0.00047613 | hemP |
| A0A0D5YGB3 | MFS transporter | -1.282 | 0.00415116 | ABUW_1589 |
| A0A0D5YMB0 | DUF4124 domain-containing protein | -1.272 | 0.00011047 | ABUW_3358 |
| A0A0D5YE68;A0A0D5YFK0 | BppU N-terminal domain-containing protein | -1.262 | 0.09854608 | ABUW_0789;ABUW_1307 |
| A0A0D5YDQ7 | CheW-like domain-containing protein | -1.261 | 0.00381671 | ABUW_0683 |
| A0A0D5YFW2 | Diguanylate cyclase | -1.249 | 0.00060083 | ydaM |
| A0A059ZMQ3 | Cytochrome D ubiquinol oxidase, subunit II | -1.242 | 0.00031 | cydB |
| A0A0D5YD66 | Oxygenase | -1.237 | 0.00440474 | ABUW_0470 |
| A0A059ZFC5 | Phosphopantetheine adenylyltransferase | -1.234 | 0.00336903 | coaD |
| A0A0D5YDV8 | Peptidase S8 | -1.234 | 0.12136033 | ABUW_0358 |
| A0A0C0AGE3 | Endoribonuclease L-PSP family protein | -1.233 | 0.00311071 | A7M90_17615 |
| A0A0D5YNC1 | DNA-processing protein DprA | -1.229 | 0.01701205 | dprA |
| A0A0D5YMX6 | HAD-superfamily hydrolase | -1.226 | 0.00014986 | ABUW_3475 |
| A0A0D5YGW1 | Cytochrome D ubiquinol oxidase subunit I | -1.213 | 0.00073259 | cydA |
| V5VE35 | Uncharacterized protein | -1.209 | 5.75E-05 | A7M90_05605 |
| A0A0D5YMR3 | Heme chaperone HemW | -1.197 | 0.00095824 | hemW |
| V5VB38 | Hotdog fold thioesterase | -1.192 | 0.01554753 | A7M90_09990 |
| A0A0D5YF23 | Uncharacterized protein | -1.177 | 0.00270594 | ABUW_0740 |
| A0A0D5YJP1 | ATP-binding cassette domain-containing protein | -1.17 | 0.0034448 | ABUW_2422 |
| V5VFD0 | Transcriptional regulator | -1.159 | 1.72E-05 | bigR_1 |
| A0A0D5YDX1 | Pilus assembly protein tip-associated adhesin PilY1 | -1.156 | 0.00055242 | pilY |
| A0A0D5YMA4 | Ohr family peroxiredoxin | -1.155 | 0.10467185 | ABUW_3265 |
| A0A0D5YNV3 | Glutathionylspermidine synthase | -1.154 | 0.0091668 | ABUW_3893 |
| V5VDY2 | Glutaredoxin 2 | -1.138 | 0.00319678 | A7M90_11265 |
| A0A0D5YEI5 | Glutamate dehydrogenase | -1.128 | 3.12E-05 | gdh |
| A0A0D5YEM2 | Hydrolase, NUDIX family protein | -1.121 | 0.00066243 | ABUW_0390 |
| A0A059ZIH5 | Dual-specificity RNA pseudouridine synthase RluA | -1.12 | 2.84E-07 | rluA |
| A0A1A1SVI5 | Rubredoxin | -1.116 | 0.00794379 | rubA |
| A0AA90HUQ3 | Carnitine monooxygenase oxygenase subunit | -1.109 | 0.00059731 | cntA |
| V5VHU8 | NADH-quinone oxidoreductase subunit E | -1.106 | 0.00285082 | nuoE |
| A0A059ZLR1 | YaeQ family protein | -1.106 | 0.00386126 | A7M90_10635 |
| A0A059ZPS2 | Uncharacterized protein | -1.104 | 0.06810429 | A7M90_03660 |
| A0A0D5YGX9 | Uncharacterized protein | -1.101 | 0.00399465 | ABUW_1414 |
| A0A090B1U5 | Maleylacetoacetate isomerase | -1.099 | 3.89E-06 | maiA |
| A0A0D5YLN2 | Type IV-A pilus assembly ATPase PilB | -1.098 | 0.00018034 | pilB |
| A0A0D5YED2 | Octanoyltransferase | -1.088 | 0.0072549 | lipB |
| A0A0E1FKE5 | Ureidoglycolate hydrolase | -1.086 | 0.0017237 | allA |
| A0A0D5YCW8 | Fimbrial assembly protein PilQ | -1.085 | 3.25E-07 | comQ |
| A0A0D5YME7 | LysM peptidoglycan-binding domain-containing protein | -1.085 | 8.40E-11 | ABUW_3722 |
| A0A0D5YG83 | Poly-beta-1,6-N-acetyl-D-glucosamine synthase | -1.079 | 0.0214259 | pgaC |
| A0A090B4L5 | Response regulator | -1.073 | 0.03837809 | cusR_1 |
| V5VHV3 | Uncharacterized protein | -1.071 | 0.00033088 | A7M90_13440 |
| A0A0D7R5E0 | PqqA binding protein | -1.065 | 0.00526662 | pqqD |
| A0A0D5YGS8 | Kynureninase | -1.065 | 0.01807245 | kynU |
| V5VEK5 | Methyltransferase | -1.062 | 0.00991528 | A7M90_06535 |
| A0A098SE84 | Pyridoxine/pyridoxamine 5'-phosphate oxidase | -1.058 | 0.00018131 | pdxH |
| A0A0E1FJA9 | LemA family | -1.055 | 1.17E-05 | ABUW_0534 |
| V5VFS6 | Aspartate/glutamate leucyltransferase | -1.052 | 0.00359859 | ate |
| A0A0D8ENG5 | Uncharacterized protein | -1.052 | 0.00030125 | ABUW_3016 |
| A0A0D5YFJ4 | Phage tail protein | -1.049 | 0.16494866 | ABUW_0791 |
| R9RIN9 | TssB | -1.048 | 0.01148717 | tssB |
| A0A0D5YJ98 | Lipoyl synthase | -1.047 | 0.00117967 | lipA2 |
| A0A0D5YF59 | Xanthine dehydrogenase small subunit | -1.045 | 0.00139689 | xdhA |
| V5VIJ9 | RNA pyrophosphohydrolase | -1.043 | 0.00069574 | nudH |
| A0A0D5YI12 | IucA/IucC-family aerobactin siderophore biosynthesis component | -1.041 | 0.00022785 | ABUW_2189 |
| A0A0D5YIJ7 | ABC transporter permease subunit | -1.038 | 0.00268816 | ABUW_2423 |
| A0A0D5YEC9 | Uncharacterized protein | -1.031 | 0.00871731 | ABUW_0902 |
| V5VAA4 | Phosphate regulon transcriptional regulatory protein PhoB | -1.026 | 0.00053873 | phoB |
| A0A0D5YL17 | 5-formyltetrahydrofolate cyclo-ligase | -1.026 | 0.04165889 | ABUW_2863 |
| A0A0D5YCB8 | Polysaccharide deacetylase | -1.022 | 0.00013105 | ABUW_0132 |
| A0A0D5YCF3 | Ribosomal RNA small subunit methyltransferase E | -1.021 | 0.08661082 | ABUW_0172 |
| A0A0D5YGA9 | Xanthine dehydrogenase molybdopterin binding subunit | -1.02 | 0.02148157 | xdhB |
| V5VFV8 | Uncharacterized protein | -1.019 | 0.01890099 | A7M90_10965 |
| A0A059ZH17 | Amidophosphoribosyltransferase | -1.016 | 0.12350538 | ABUW_0391 |
| V5V966 | FAD assembly factor SdhE | -1.014 | 0.00108819 | cptB |
| A0A0D5YGA4 | DUF3015 domain-containing protein | -1.009 | 0.15588702 | ABUW_1192 |
| V5VGB8 | ADP-ribose pyrophosphatase | -1.006 | 0.03853127 | ABUW_3239 |
| A0A0D5YI69 | TetR family transcriptional regulator | -1.006 | 0.05376666 | ABUW_1895 |
| A0A0D5YHL8 | TonB-dependent siderophore receptor | -1.005 | 4.17E-06 | ABUW_1655 |
| V5V8M6 | DUF551 domain-containing protein | -0.999 | 9.37E-05 | A7M90_07695 |
| V5VGS0 | Nuclear transport factor 2 family protein | -0.997 | 0.00190262 | A7M90_09040 |
| V5VDC6 | Small ribosomal subunit protein bS6 | -0.996 | 0.00147125 | rpsF |
| A0A059ZHC0 | Protein tyrosine phosphatase family protein | -0.992 | 0.00442923 | ABUW_1729 |
| A0A0D5YJ03 | Siderophore biosynthesis protein | -0.986 | 2.60E-06 | ABUW_2186 |
| A0A0D5YD81 | Uncharacterized protein | -0.984 | 0.00676696 | ABUW_0134 |
| V5VAR7 | NF038105 family protein | -0.977 | 0.00025642 | A7M90_09740 |
| V5VDG9 | Acetyltransferase | -0.973 | 7.71E-05 | mbtK |
| V5V8X8 | Antibiotic biosynthesis monooxygenase | -0.972 | 0.01259487 | ygiN |
| A0A0D5YIR9 | ADP-ribose pyrophosphatase | -0.971 | 0.01927189 | ABUW_1944 |
| A0A0D5YEF4 | SgcJ/EcaC family oxidoreductase | -0.967 | 0.00760993 | ABUW_0931 |
| A0A0D5YJD9 | Putative 4-hydroxy-4-methyl-2-oxoglutarate aldolase | -0.966 | 6.06E-05 | ABUW_2183 |
| V5VGZ7 | Peptide methionine sulfoxide reductase MsrA | -0.958 | 0.00117305 | msrA |
| A0A0D5YKX4 | ComEA family DNA-binding protein | -0.945 | 3.32E-07 | ABUW_3299 |
| V5VHB7 | DUF2061 domain-containing protein | -0.942 | 7.97E-06 | ABUW_3464 |
| A0A0D5YG16 | CsuC | -0.942 | 0.00013231 | ABUW_1490 |
| A0A0D5YKP6 | Ribonuclease HII | -0.93 | 2.44E-06 | rnhB |
| A0A0D5YK34 | Heme oxygenase | -0.925 | 0.0002324 | ABUW_2981 |
| A0A059ZH19 | L-idonate 5-dehydrogenase (NAD(P)(+)) | -0.918 | 0.1120667 | idnD |
| A0A0D5YLB1 | Cobalamin biosynthesis protein CobW | -0.915 | 0.00104359 | ABUW_2964 |
| V5VB54 | Acyl-CoA desaturase | -0.912 | 0.00021413 | desA3_1 |
| E2FGC2 | PmrA | -0.908 | 0.00257575 | pmrA |
| A0A0D5YMD7 | MazG family protein | -0.904 | 0.00698312 | mazG |
| A0A0D5YML9 | tRNA (guanine(46)-N(7))-methyltransferase | -0.901 | 0.01262091 | ABUW_3495 |
| A0A0D5YHM3 | ABC transporter, ATP-binding protein | -0.899 | 0.00946403 | ABUW_1660 |
| A0A0D5YF88 | Methyltransferase | -0.899 | 0.02584948 | ABUW_0782 |
| V5VCJ3 | Fumarate hydratase class II | -0.897 | 2.54E-05 | fumC |
| A0A0D5YKM9 | diguanylate cyclase | -0.893 | 0.07783009 | ABUW_3178 |
| C5H8J6 | CadR | -0.889 | 0.05076967 | cadR |
| A0A0D5YDE0 | Rha family transcriptional regulator | -0.887 | 0.00014286 | ABUW_0551 |
| A0A086HY23 | Alpha/beta hydrolase | -0.881 | 0.05179341 | aesT |
| A0A0D5YKQ6 | Aspartate 1-decarboxylase | -0.873 | 0.0342357 | panD |
| V5VA86 | Chemotaxis protein | -0.869 | 0.00116679 | pilJ |
| A0A0D5YD19 | N-succinylarginine dihydrolase | -0.866 | 0.00021309 | astB |
| V5VHB6 | Ribosome maturation factor RimP | -0.865 | 0.0122319 | rimP |
| A0A0D5YL33 | p-aminobenzoate synthetase PabB | -0.86 | 0.00200851 | pabB |
| A0A0D5YKU5 | 5'-methylthioadenosine nucleosidase | -0.86 | 7.43E-05 | mtnN |
| A0A0D5YJ36 | Siderophore biosynthesis protein | -0.859 | 1.34E-05 | ABUW_2185 |
| V5VB87 | Sodium/glutamate symporter | -0.859 | 0.00068099 | gltS |
| A0A0D5YEJ2 | DUF4112 domain-containing protein | -0.858 | 0.05394345 | ABUW_0923 |
| A0A059ZM52 | protein-tyrosine-phosphatase | -0.854 | 0.01810592 | yfkJ |
| V5VHJ1 | Cation:proton antiporter | -0.849 | 0.13606476 | phaG |
| A0A0D5YNU8 | UDP-N-acetylmuramate--L-alanyl-gamma-D-glutamyl-meso-2,6-diaminoheptandioate ligase | -0.848 | 0.00129944 | mpl |
| U5QBH4 | Uncharacterized protein | -0.847 | 0.00282855 | ABUW_4041 |
| A0A090C0R2 | Glycerophosphodiester phosphodiesterase | -0.847 | 0.00654835 | ABUW_0613 |
| V5VDN3 | Transcriptional regulator | -0.843 | 0.00103106 | asnC_2 |
| V5VG41 | ImpA-related N-terminal family protein | -0.842 | 0.00176536 | tssA |
| A0A0D5YMY2 | Lipoprotein, putative | -0.838 | 0.00033594 | ABR2091_3691 |
| A0A1A9LAX8 | 2OG-Fe(II) oxygenase superfamily protein | -0.829 | 1.41E-07 | piuC |
| Q5DL40 | CysA | -0.827 | 2.12E-05 | cysA |
| A0A0D5YFA1 | 3'-phosphate/5'-hydroxy nucleic acid ligase | -0.824 | 0.00247225 | ABUW_0651 |
| N0A217 | Branched-chain-amino-acid aminotransferase | -0.822 | 0.0041756 | ilvE |
| A0A059ZGM7 | Ferrochelatase | -0.819 | 0.02907297 | hemH |
| V5VI82 | Putative membrane protein insertion efficiency factor | -0.816 | 0.04122797 | yidD |
| V5VCG2 | 3-phenylpropionate dioxygenase ferredoxin subunit | -0.815 | 3.22E-06 | ABUW_1798 |
| A0A0D5YKX3 | Adenine deaminase | -0.81 | 0.00042719 | add1 |
| A0A0D5YK48 | Class I SAM-dependent methyltransferase | -0.808 | 0.00967587 | ABUW_2449 |
| A0A059ZT14 | Lipoprotein | -0.808 | 0.00096517 | A7M90_19160 |
| A0A0D5YKY4 | Hydroxypyruvate isomerase | -0.802 | 0.02090498 | hyi |
| A0A0D5YL23 | Ribosomal RNA small subunit methyltransferase J | -0.802 | 0.00257047 | rsmJ |
| V5VE58 | Internalin | -0.796 | 0.00329571 | A7M90_02495 |
| A0A0D8GGD3 | Chemotaxis protein CheW | -0.796 | 1.06E-05 | pilI |
| A0A0D8GFP2 | Type III pantothenate kinase | -0.795 | 0.00028886 | coaX |
| A0A0D5YIF1 | Uncharacterized protein | -0.795 | 2.09E-05 | ABUW_1994 |
| A0A0D5YLC1 | Aldehyde dehydrogenase | -0.795 | 0.00035481 | calB |
| A0A0D5YG12 | RNA polymerase-associated protein RapA | -0.794 | 0.00878894 | hepA |
| V5VAI1 | Acidic transcription factor A | -0.794 | 0.05648999 | A7M90_18910 |
| V5VCP1 | DUF3188 domain-containing protein | -0.793 | 5.13E-06 | A7M90_02265 |
| A0A0D5YDK6 | LysR family transcriptional regulator | -0.792 | 0.014887 | ABUW_0214 |
| V5VH71 | DSBA oxidoreductase | -0.792 | 0.00252481 | A7M90_01485 |
| V5VDM2 | 2Fe-2S ferredoxin | -0.79 | 0.00183944 | fdx |
| A0A0D5YDV3 | N-succinylglutamate 5-semialdehyde dehydrogenase | -0.789 | 3.03E-06 | astD |
| A0A059ZLJ4 | Bestrophin | -0.786 | 0.00055458 | A7M90_17495 |
| A0A0D5YK30 | Beta-ketoadipyl-CoA thiolase | -0.784 | 2.77E-05 | paaE |
| Q2FCW2 | STAS domain-containing protein | -0.782 | 0.00419764 | sulI |
| B4YSY0 | histidine kinase | -0.78 | 0.07791506 | pilS |
| V5VAT6 | ABC transporter permease | -0.779 | 3.92E-06 | sbp_1 |
| V5VF25 | Cu(I)-responsive transcriptional regulator | -0.779 | 0.00517602 | cueR |
| V5V8D5 | Iron-sulfur cluster insertion protein ErpA | -0.777 | 1.68E-05 | erpA |
| A0A090C1U7 | Acetyltransferase, GNAT family | -0.775 | 0.059239 | ABUW_2739 |
| A0A090B989 | Peptide chain release factor 1 | -0.775 | 0.00193164 | prfA |
| A0A0D5YIU7 | Surface antigen | -0.771 | 0.00019143 | ABUW_2440 |
| A0A086HW53 | Small ribosomal subunit biogenesis GTPase RsgA | -0.771 | 3.66E-07 | rsgA |
| A0A0D5YK07 | Chemotaxis protein | -0.77 | 4.51E-06 | ABUW_2842 |
| A0A0D5YII5 | Uncharacterized protein | -0.763 | 3.49E-05 | ABUW_1983 |
| A0A0D5YDW9 | Microcin B17 transport protein | -0.761 | 0.00436385 | ABUW_0707 |
| A0A0D5YCJ8 | Alkene reductase | -0.76 | 0.00433674 | xenB |
| V5VHW4 | Uncharacterized protein | -0.758 | 0.02530366 | A7M90_19200 |
| V5VGE8 | Integration host factor subunit alpha | -0.758 | 0.01211731 | ihfA |
| V5VAK1 | Metal binding protein | -0.757 | 0.01048899 | A7M90_18905 |
| V5VH89 | Fe2+ transport system protein A | -0.755 | 0.00352741 | feoA |
| V5V960 | Endonuclease | -0.751 | 6.50E-06 | A7M90_15425 |
| A0A0D5YJK9 | Taurine ABC transporter substrate-binding protein | -0.751 | 1.62E-07 | tauA |
| A0A059ZM24 | UPF0301 protein A7M90_01390 | -0.75 | 0.00472252 | A7M90_01390 |
| A0A0D5YG95 | Histidine decarboxylase | -0.748 | 0.00024508 | basG |
| A0A059ZN00 | Phosphatase NudJ | -0.747 | 0.02323567 | nudJ |
| V5VHC0 | RNA polymerase-binding transcription factor DksA | -0.746 | 0.00057618 | dksA |
| V5VH14 | Efflux transporter periplasmic adaptor subunit | -0.745 | 4.49E-05 | yibH_1 |
| V5VGM2 | Gluconate permease | -0.743 | 8.02E-06 | gntT |
| A0A0D5YDH6 | Uncharacterized protein | -0.741 | 0.15945266 | ABUW_0549 |
| A0A0D5YIF4 | Crossover junction endodeoxyribonuclease RuvC | -0.741 | 0.00368159 | ruvC |
| A0A0D7U5I7 | HAD-IB family hydrolase | -0.74 | 0.0196157 | ABR2091_0413 |
| A0A0D5YLB6 | Twitching mobility protein | -0.736 | 3.54E-08 | pilT |
| A0A0D5YHI6 | phospholipase C | -0.735 | 4.61E-05 | plc1 |
| A0A059ZCX7 | Transcription termination/antitermination protein NusG | -0.735 | 0.01302905 | nusG |
| A0A0D5YK70 | Delta-aminolevulinic acid dehydratase | -0.731 | 0.02358714 | hemB |
| A0A0D5YDY0 | Probable membrane transporter protein | -0.729 | 0.00116021 | yfcA |
| V5VEM0 | Ferredoxin | -0.728 | 0.00074404 | A7M90_06720 |
| V5VI32 | CoA pyrophosphatase | -0.727 | 0.00874044 | ABR2091_0671 |
| V5VD64 | DUF6231 family protein | -0.726 | 0.00194851 | ABUW_2162 |
| A0A0D5YGN7 | Alpha/beta hydrolase | -0.724 | 8.94E-05 | ABUW_1364 |
| V5VIX7 | Phosphodiesterase | -0.716 | 0.0119651 | cpdA |
| V5VHF8 | SirA-like protein | -0.715 | 0.12613016 | tusA |
| A0A0D5YEB6 | Acyl-CoA carboxylase subunit beta | -0.714 | 0.00414435 | ABUW_0486 |
| V5VAC4 | DUF493 domain-containing protein | -0.709 | 0.00123217 | ybeD |
| V5VB25 | Elongation factor 4 | -0.709 | 0.00025051 | lepA |
| V5VF16 | Transcriptional regulator SutA RNAP-binding domain-containing protein | -0.708 | 0.00087101 | ABR2091_1029 |
| A0A0D5YMD8 | Putative hydrolase | -0.707 | 0.02984081 | ABUW_3388 |
| V5VAW9 | Peptidyl-prolyl cis-trans isomerase | -0.705 | 0.02357782 | slyD |
| A0A0D5YG22 | Aldehyde dehydrogenase | -0.704 | 0.00084771 | ABUW_1111 |
| A0A0D5YEH6 | Methyltransferase | -0.702 | 6.37E-06 | prmA2 |
| A0A059ZQ34 | FHA domain protein | -0.701 | 0.00194354 | ABUW_3605 |
| V5VGH1 | MFS transporter | -0.697 | 0.00061837 | emrB_1 |
| V5V9H6 | Arginine N-succinyltransferase | -0.695 | 4.97E-05 | astA |
| A0A0D5YJ32 | PepSY domain-containing protein | -0.692 | 6.76E-05 | ABUW_2180 |
| A0A0D5YF50 | Thioesterase | -0.69 | 0.00717325 | basH |
| A0A0D5YGL9;V5VHD2 | UPF0056 membrane protein | -0.688 | 0.09720103 | ABUW_1155;yhgN |
| V5VI47 | DUF4951 domain-containing protein | -0.686 | 3.14E-05 | ABUW_3263 |
| A0A0D5YD05 | UPF0260 protein ABUW_0054 | -0.682 | 0.01545969 | ABUW_0054 |
| V5V842 | Transcription antitermination protein NusB | -0.681 | 0.00481539 | nusB |
| V5VDP3 | AraC family transcriptional regulator | -0.681 | 0.01253678 | ureR_3 |
| A0A0D5YCB0 | PGAP1 family protein | -0.678 | 0.01024158 | ABUW_0122 |
| V5VC23 | Putative sterol carrier protein | -0.675 | 0.00029187 | A7M90_09420 |
| A0A0D5YL73 | Hemophilin receptor | -0.673 | 0.01961599 | hphR |
| A0A0D5YLD0 | Uncharacterized protein | -0.672 | 5.61E-06 | ABUW_3439 |
| A0A059ZMW7 | Signal peptide | -0.671 | 7.81E-06 | A7M90_02030 |
| A0A0D5YLV9 | DUF2218 domain-containing protein | -0.67 | 0.00328165 | ABUW_3232 |
| A0A098SCQ9 | DUF1289 domain-containing protein | -0.669 | 0.00020203 | ABUW_0866 |
| Q2FD59 | Amino acid permease | -0.662 | 0.00128178 | cycA_3 |
| A0A0E1FDD0 | Electron transport complex, rnfaBcdge type, B subunit | -0.661 | 6.10E-05 | rnfB |
| A0A059ZLL3 | Amino acid synthesis family protein | -0.661 | 0.00497563 | A7M90_05935 |
| V5VI29 | Hypoxanthine phosphoribosyltransferase | -0.66 | 8.00E-06 | hpt |
| A0A081GRI4 | Acetyltransferase | -0.66 | 0.00702141 | ypeA |
| A0A0D5YCJ1 | Flavoprotein | -0.66 | 0.00193158 | ABR2091_3523 |
| A0A0D5YEF3 | PHP domain protein | -0.657 | 0.12151785 | ABUW_0526 |
| C9DPL8 | Heteromeric transposase endonuclease subunit TnsA | -0.654 | 0.01284254 | tniC |
| A0A090BE98 | ImmA/IrrE family metallo-endopeptidase | -0.654 | 0.00468106 | ABUW_3319 |
| A0A0D5YID6 | Transcriptional regulator, TetR family | -0.653 | 0.00579498 | ABUW_1979 |
| V5V9S1 | YCII-related domain protein | -0.651 | 0.02473827 | A7M90_17510 |
| V5V991 | Uncharacterized protein | -0.649 | 0.00410592 | A7M90_15195 |
| A0A059ZJ17 | Phosphate ABC transporter ATP-binding protein | -0.646 | 1.05E-05 | pstB |
| A0A0D5YHP5 | Putative aliphatic sulfonates-binding protein | -0.642 | 0.01120496 | ABUW_2111 |
| A0A081H611 | UPF0125 protein A7M90_16355 | -0.641 | 0.00042391 | ratB |
| A0A0D5YHW2 | DHA2 family efflux MFS transporter permease subunit | -0.641 | 0.00155739 | ABUW_2187 |
| A0A0E1FT68 | Uncharacterized protein | -0.636 | 0.06137416 | ABUW_2200 |
| A0A0D5YIL6 | Oxidoreductase FMN-binding | -0.634 | 0.00943716 | ABUW_1887 |
| A0A0D5YMV7 | tRNA pseudouridine synthase A | -0.631 | 0.0045835 | truA |
| V5VAX6 | Pseudouridine synthase | -0.631 | 0.00200505 | rluE |
| A0A0D5YCA9 | fumarylacetoacetase | -0.629 | 0.00428077 | fahA |
| V5V9E1 | DNA-directed RNA polymerase subunit omega | -0.627 | 0.02561074 | rpoZ |
| V5V8S6 | Pyruvate dehydrogenase E1 component | -0.625 | 8.89E-05 | aceE |
| A0A0D5YCL9 | Transcriptional regulator | -0.624 | 0.02531742 | ABUW_0224 |
| A0A0D5YEQ7 | Intracellular protease, PfpI family | -0.623 | 0.00941798 | ABUW_0711 |
| V5VGJ9 | Oligoribonuclease | -0.62 | 0.00177712 | orn |
| A0A0D5YJT2 | ABC transporter, periplasmic binding protein | -0.619 | 0.00019113 | ABUW_2421 |
| A0A0D5YMD6 | tRNA-dihydrouridine(20/20a) synthase | -0.616 | 0.00014067 | dusA |
| A0A0D5YJZ8 | Aldehyde dehydrogenase | -0.613 | 0.12501588 | ABUW_2496 |
| V5VH01 | HTH-type transcriptional repressor KstR2 | -0.613 | 3.37E-05 | kstR2_2 |
| A0A0D5YLB3 | Urease subunit alpha | -0.612 | 0.00030774 | ureC |
| A0A0D5YC39 | 2,5-didehydrogluconate reductase B | -0.611 | 0.00011205 | alrA |
| A0A0D5YCZ5 | Lipoprotein | -0.608 | 0.00091137 | ABUW_0044 |
| A0A0D5YIN3 | Isochorismatase | -0.606 | 1.40E-05 | ABUW_2075 |
| V5VE65 | ATP-binding cassette domain-containing protein | -0.606 | 0.04286705 | artP |
| A0A0D5YEG4 | S-adenosylmethionine:tRNA ribosyltransferase-isomerase | -0.602 | 0.00729454 | queA |
| V5VEP4 | Copper chaperone | -0.601 | 2.22E-05 | A7M90_06740 |
| A0A090C286 | Chorismate mutase family protein | -0.601 | 0.01202928 | ABUW_3341 |
| V5V9Z8 | Tryptophan--tRNA ligase | -0.599 | 0.00973403 | trpS |
| V5VHD8 | Oxygen-dependent choline dehydrogenase | -0.597 | 1.58E-05 | betA |
| A0A0D5YHW7 | Tryptophan synthase beta chain | -0.597 | 0.00060919 | trpB2 |
| A0A0D5YF28 | Uncharacterized protein | -0.597 | 0.01324417 | ABUW_0745 |
| A0A0D5YDI5 | Uncharacterized protein | -0.596 | 0.00024659 | ABUW_0602 |
| A0A059ZG02 | Threonylcarbamoyl-AMP synthase | -0.595 | 0.06206882 | tsaC |
| A0A0D5YJI7 | GFA family protein | -0.594 | 0.01074927 | ABUW_2705 |
| V5V8Q6 | 6,7-dimethyl-8-ribityllumazine synthase | -0.591 | 0.01882617 | ribE |
| A0A0E1JMW8 | LysR family transcriptional regulator | -0.587 | 0.08575533 | ABUW_2931 |
| A0A0D5YIJ8 | Uncharacterized protein | -0.587 | 0.00015637 | ABUW_2040 |
| A0A0D5YKA1 | Threonine efflux protein | -0.582 | 0.01712355 | rhtC |
| A0A0D5YI08 | (2Fe-2S)-binding protein | -0.582 | 0.09609374 | ABUW_2184 |
| V5V902 | Small-conductance mechanosensitive channel | -0.582 | 0.00538206 | mscS_2 |
| A0A0D5YEI9 | Succinylglutamate desuccinylase | -0.581 | 2.22E-09 | astE |
| V5VB93 | L-lysine 2,3-aminomutase | -0.581 | 0.02106915 | epmB |
| V5V9S3 | UPF0761 membrane protein A7M90_15130 | -0.58 | 0.00833751 | yihY |

***Table S5. Hfq Clash-seq***

***Table S6. iModulion***

***Table S7 iModulion***

***Table S8. Clinical Acinetobacter baumannii isolates identified with cavA F237V mutation.***

| **Strain** | **SRA** | **Biosample** | **Date collected** | **Submission** | **Source** | **MLST** |
| --- | --- | --- | --- | --- | --- | --- |
| 2023DK-00246 | SRS18760969 | SAMN37217195 | 2023 | Massachusetts State Public Health Lab | Wound | MLST79 |
| 2020CK-00433 | SRS7681837 | SAMN16729807 | 2020 | Utah Public Health Laboratory | Wound | MLST78 |
| 2020JQ-00146 | SRS11063236 | 2020JQ-00146 | 2020 | Washington State Department of Health | Urine | MLST79 |
| 2024HL-00055 | SRS20101721 | SAMN39331322 | 2023 | Wadsworth Center - NYS DOH | Sputum | MLST499 |
| 2024HL-00975 | SRS22597251 | SAMN43548122 | 2024 | Wadsworth Center - NYS DOH | Axilla, groin, perianal swab | MLST499 |
| 2023HL-00567 | SRS18464409 | SAMN36781179 | 2023 | Wadsworth Center - NYS DOH | Ulcer | MLST499 |
| 2019CK-00023 | SRS6608260 | SAMN14792028 | 2019 | Utah Public Health Laboratory | Other | MLST78 |
| 2020CK-00065 | SRS6608189 | SAMN14792070 | 2020 | Utah Public Health Laboratory | Urine | MLST78 |
| 2023HL-00687 | SRS18827104 | SAMN37314367 | 2023 | Wadsworth Center - NYS DOH | Perianal swab | MLST499 |
| 2021HL-00891 | SRS12153019 | SAMN26365924 | 2021 | Wadsworth Center - NYS DOH | Tracheal aspirate | MLST499 |
| 2024HL-00971 | SRS22559235 | SAMN43514801 | 2024 | Wadsworth Center - NYS DOH | Sputum | MLST499 |
| s063070 | NA | SAMN41860599 | 2020 | Peking University, China | Hospital | NA |
| 2024HL-00213 | SRS20721414 | SAMN40376014 | 2024 | Wadsworth Center - NYS DOH | Urine | MLST499 |

**Table S9.** *A. baumannii chromosomal differences following serial passage in cefiderocol as identified by Illumina whole genome sequencing. Locus_tags are relevant to the A. baumannii reference genome ASM903584v1 (GCF_009035845.1)*

| ***A. Baumannii* strain M87136** | | | |
| --- | --- | --- | --- |
| **Passage day** | **Locus_tag** | **Gene; Product** | **Mutation** |
| 2 |  |  | None |
| 4 |  |  | None |
| 9 |  |  | None |
| ***A. Baumannii* strain M00376** | | | |
| **Passage day** | **Locus_tag** | **Gene; Product** | **Mutation** |
| 2 | FQU82_RS12695 | *DNA-binding transcriptional regulator Fis* | Leu81* |
| 8 | FQU82_RS12695 | *DNA-binding transcriptional regulator Fis* | Leu81* |
| 10 | FQU82_RS12695 | *DNA-binding transcriptional regulator Fis* | Leu81* |
| ***A. Baumannii* strain M88825** | | | |
| **Passage day** | **Locus_tag** | **Gene; Product** | **Mutation** |
| 2 | - | *-* | None |
| 7 |  |  | None |
| 9 |  |  | None |
| 11 |  |  | None |
| ***A. Baumannii* strain M87341** | | | |
| **Passage day** | **Locus_tag** | **Gene; Product** | **Mutation** |
| 2 |  |  | None |
| 4 |  |  | None |
| 7 |  |  |  |
| 9 | FQU82_RS15440 | *BapA/Bap/LapF family large adhesin* | Ile241Val |
| 11 | FQU82_RS18455 | *Cell division protein FtsZ* | Ala119Val |

**Table S10. TraDIS insertion sites**

| **Locus_tag** | **Name** | **Width** | | **Nsites** | **Normalisation** |
| --- | --- | --- | --- | --- | --- |
| IGR2987 | IGR2987 | 86 | | 19 | 153825 |
| MS7163_RS19940 | *envZ* | 1353 | | 124 | 120234 |
| MS7163_RS07195 | *tonB* | 717 | | 117 | 98379 |
| MS7163_RS11820 | - | 585 | | 17 | 20328 |
| MS7163_RS00405 | *leuL* | 87 | | 11 | 12829 |
| MS7163_RS19180 | - | | 855 | 97 | 11015 |
| MS7163_RS22210 | *cyaA* | 2547 | | 338 | 10221 |
| MS7163_RS19730 | *crp* | 633 | | 159 | 7666 |
| IGR2270 | IGR2270 | 5883 | | 718 | 4384 |
| IGR1744 | IGR1744 | 5583 | | 686 | 4200 |

**Table S11. Cefiderocol MIC of *E. coli* MS7163 and its derivatives**

| Strain | Cefiderocol MIC (μg/ml) | | |
| --- | --- | --- | --- |
|  | Repeat 1 | Repeat 2 | Repeat 3 |
| MS7163∆cyaA | 0.5 | 0.0625 | 0.0625 |
| MS7163∆cyaA + pSU2718 | 0.25 | 0.125 | 0.125 |
| MS7163∆cyaA + pcyaA | 0.25 | 0.125 | 0.125 |
| MS7163 wild-type | 0.25-0.5 | 0.0625 | 0.125 |

**Table S12. Bacterial strains employed in this study**

| ***A. baumannii* isolates** | | | | |
| --- | --- | --- | --- | --- |
| **Strain** | **ST** | **Isolation site** | **Origin** | **Reference** |
| AB5075 | ST1 | Wound | USA | ^1^ |
| M89059 | ST195 | Rectal | Australia | This study |
| M87136 | ST218 | Respiratory | Australia | This study |
| M92179 | ST229 | Blood | Australia | This study |
| M00376 | ST451 | Wound | Australia | This study |
| M810891 | ST784 | Endotracheal aspirate | Australia | This study |
| M86168 | ST878 | Respiratory | Australia | ^2^ |
| M88825 | ST1050 | Environmental | Australia | ^2^ |
| M87341 | ST1050 | Wound / skin / soft tissue | Australia | ^2^ |
| ***E. coli* isolates** | | | | |
| **Strain** | **ST** | **Isolation site** | **Origin** | **Reference** |
| MS7631 | ST95 | Kidneys/Pyelonephritis | Slovakia | ^3^ |

**Table S13. Primers employed for quantitative-real-time PCR.**

| **Gene** | **Primer direction** | **Sequence 5’🡪3’** |
| --- | --- | --- |
| *bauB_F* | Fwd | GGGCAACCTATTTCGAGTGA |
| *bauB_R* | Rev | CGTTTTTCCATGCTTTGGTT |
| *bauE_F* | Fwd | CTCAGGCGCTTATGTGATGA |
| *gyrA_F* | Fwd | GAACCATGCGACCGTACTTT |
| *gyrA_R* | Rev | CAAGATTGCGGTTACGGTTT |
