## Supplementary material for "Disruption of cAMP Signalling and RNA Regulation Confers Cefiderocol Resistance in *Acinetobacter baumannii*": Fig S

**Supplementary Figures**

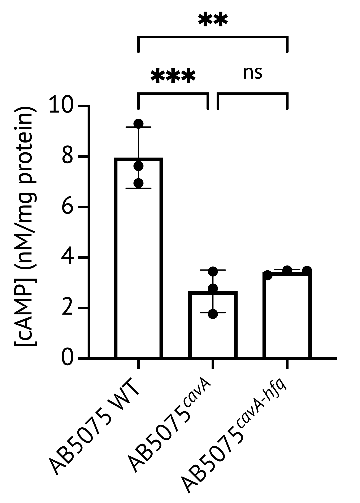

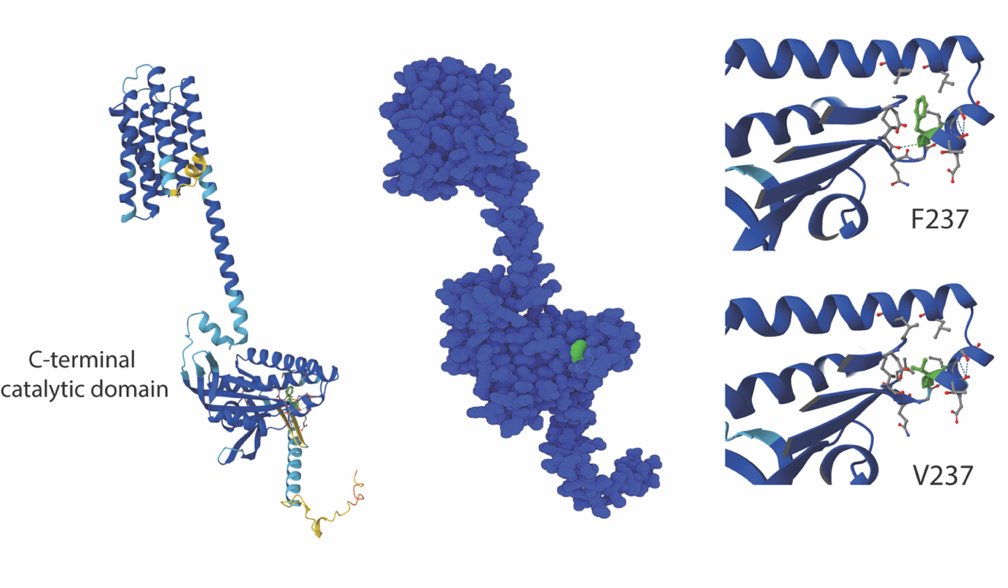

C

D

B

A

***Figure S1.*** *Protein modelling of AB5075 CavA, with ribbon (A) and globular (B) representations highlighting phenylalanine at amino acid position 237 in fluorescent green, and (C) ribbon ball-and-stick modelling illustrating the F237V mutation. Models were generated using AlphaFold, and predicted Local Distance Difference Test (pLDDT) confidence scores are color-coded from 0 to 100 in the ribbon-model as dark blue (>90), cyan (70–90), yellow (50–70), and orange (<50). (D)* cAMP concentrations in AB5075 WT, AB5075*^cavA^*  and AB5075*^cavA-hfq^*. *Data represent biological replicates. Statistical significance is indicated (***, P ≤ 0.001; **, P ≤ 0.01; one-way ANOVA with Tukey’s multiple comparisons).*

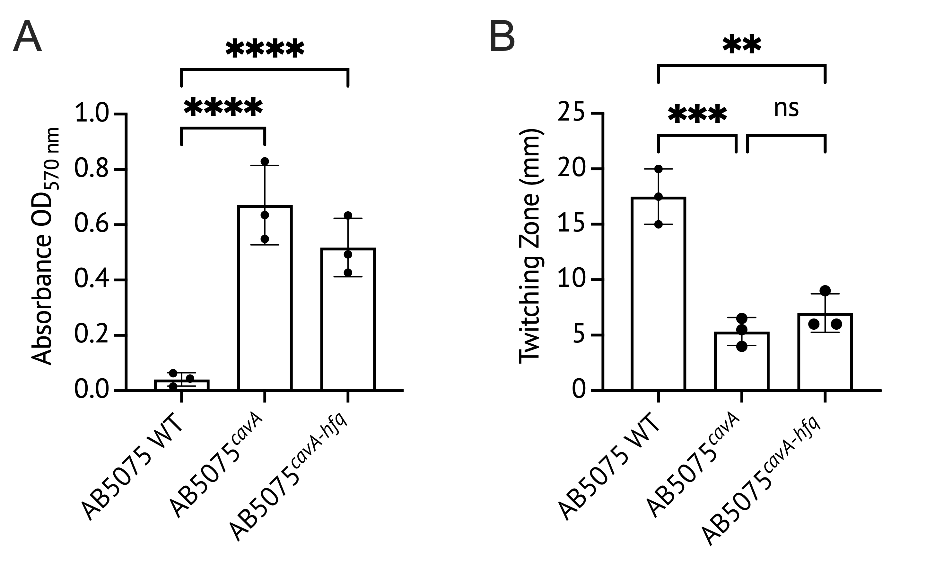

***Figure S2. Effects of cavA and hfq mutations on AB5075 biofilm formation and twitching motility.*** *(A) Biofilm formation for each strain was quantified at 24 hours using crystal violet staining, with absorbance measured at OD570 nm. (B) Twitching motility for each strain was assessed over a 24-hour period. Data represent biological replicates. Statistical significance is indicated (****, P ≤ 0.0001; one-way ANOVA with Tukey’s multiple comparisons).*

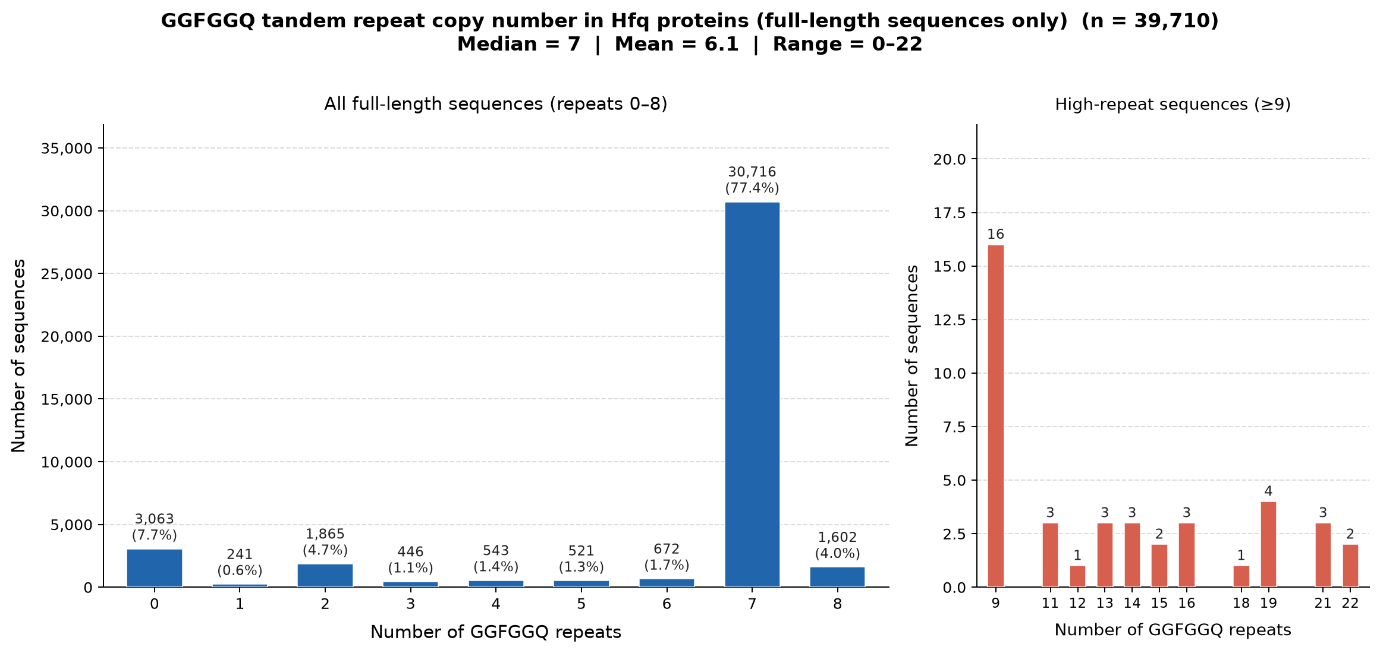

***Figure S3. The distribution of Hfq repeat domains (GGFGGQ) in A. baumannii.*** *Bar charts showing the frequency distribution of GGFGGQ hexapeptide tandem repeat copy number across 39,710 full length A. baumannii* Hfq *protein sequences retrieved from NCBI. Non-overlapping exact-match counts of the GGFGGQ motif were determined for each sequence using Python. The left panel shows the dominant repeat classes (0–8 copies); the right panel expands the rare high-repeat tail (≥9 copies). Among full-length sequences, 7 repeat copies was the overwhelmingly dominant class (30,716 sequences; 77.4%).*

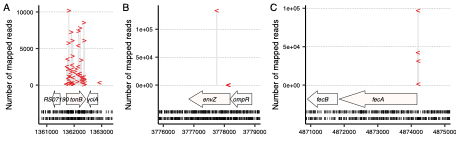

***Figure S4. Location and number of mapped reads per insertion site within the (A) tonB, (B) envZ and (C) fecA genes of E. coli strain MS7163***. Each insertion site is represented by either ‘>’ (insert on the forward strand) or ‘<’ (insert on the reverse strand). Horizontal arrows represent the coding sequences, and the locations of insertion sites in the input pool are shown by short vertical lines beneath; the top row indicates insertion sites on the forward strand, and the bottom row the reverse strand.

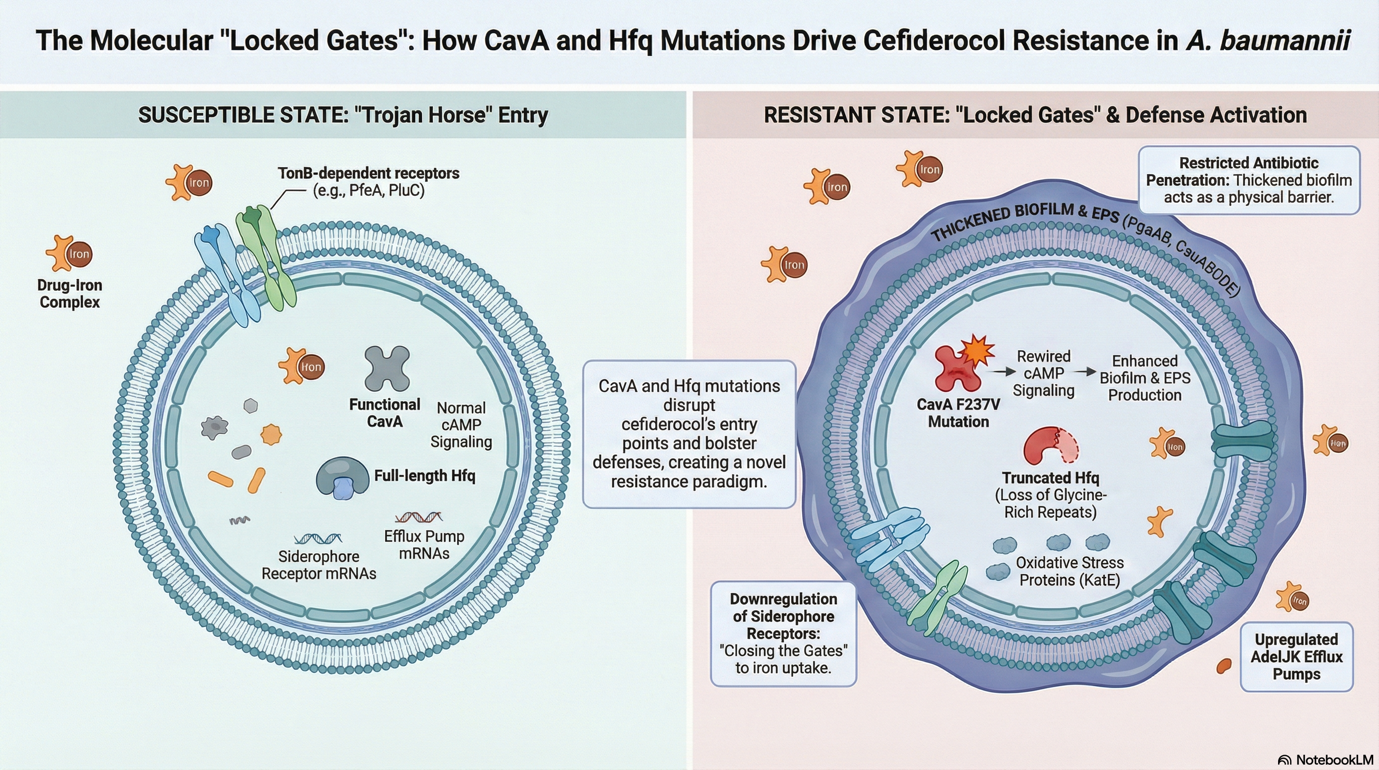

***Figure S5. CavA and Hfq mutations reprogram cefiderocol entry and stress responses to drive resistance in Acinetobacter baumannii.*** In the susceptible “Trojan horse” state (left), cefiderocol chelates iron and is actively transported across the outer membrane through TonB‑dependent siderophore receptors (e.g. PfeA, PiuC), allowing drug–iron complexes to access the periplasm and cytoplasm where canonical targets reside. Functional CavA maintains normal intracellular cAMP signalling, while expression of full‑length Hfq supports balanced transcription of siderophore receptor and efflux pump mRNAs, together permitting efficient cefiderocol uptake while limiting global stress responses. In the resistant “locked gates” state (right), a CavA F237V substitution and truncation of Hfq disrupt cAMP signalling and RNA regulation, leading to downregulation of outer‑membrane siderophore receptors (“gate closing”), upregulation of AdeIJK efflux pumps, and induction of oxidative stress response proteins (e.g. KatE). Concomitantly, activation of the PgaABCD and CsuABCD pathways promotes thickened biofilm and extracellular polysaccharide matrix formation, which physically restricts antibiotic penetration. Together, these CavA‑ and Hfq‑driven changes curtail cefiderocol entry and enhance defensive barriers, establishing a novel, multifactorial resistance mechanism.
